# Demographic histories and genome-wide patterns of divergence in incipient species of shorebirds

**DOI:** 10.1101/559633

**Authors:** Xuejing Wang, Kathryn H. Maher, Nan Zhang, Pingjia Que, Chenqing Zheng, Simin Liu, Biao Wang, Qin Huang, De Chen, Xu Yang, Zhengwang Zhang, Tamás Székely, Araxi O. Urrutia, Yang Liu

## Abstract

Understanding how incipient species are maintained with gene flow is a fundamental question in evolutionary biology. Whole genome sequencing of multiple individuals holds great potential to illustrate patterns of genomic differentiation as well as the associated evolutionary histories. Kentish (*Charadrius alexandrinus*) and the white-faced (*C. dealbatus*) plovers, which differ in their phenotype, ecology and behaviour, are two incipient species and parapatrically distributed in East Asia. Previous studies show evidence of genetic diversification with gene flow between the two plovers. Under this scenario, it is of great importance to explore the patterns of divergence at the genomic level and to determine whether specific regions are involved in reproductive isolation and local adaptation. Here we present the first population genomic analysis of the two incipient species based on the de novo Kentish plover reference genome and resequenced populations. We show that the two plover lineages are distinct in both nuclear and mitochondrial genomes. Using model-based coalescence analysis, we found that population sizes of Kentish plover increased whereas white-faced plovers declined during the Last Glaciation Period. Moreover, the two plovers diverged allopatrically, with gene flow occurring after secondary contact. This has resulted in low levels of genome-wide differentiation, although we found evidence of a few highly differentiated genomic regions in both the autosomes and the Z-chromosome. This study illustrates that incipient shorebird species with gene flow after secondary contact can exhibit discrete divergence at specific genomic regions and provides basis to further exploration on the genetic basis of relevant phenotypic traits.

## Introduction

Understanding the conditions in which speciation occurs is a fundamental question in evolutionary biology. Of equal importance is the understanding of how newly diverged species (incipient species) are maintained, as it is likely that interspecific gene flow is a common occurrence between diverging species [1]. During allopatric speciation, a physical barrier acts to prevent gene flow across the whole genome [2] and pre- and post-zygotic mechanisms of reproductive isolation can evolve to facilitate divergence. However, gene flow across geographical barriers, or secondary contact between diverged populations is possible, which allows gene flow to recommence [3]. Even infrequent gene flow can erode species barriers [4]. In the more contentious geographical context, such as parapatric or sympatric speciation [5, 6], disentangling the relative role of gene flow and other diverging conditions and forces remains challenging [7, 8].

Whether speciation can occur with gene flow has been an area of intense investigation within the last decade [9]. In certain instances, gene flow overcomes the barriers of reproductive isolation and reverse speciation processes occur, while in other species divergence persists in spite of gene flow [10–14]. The homogenising effect of gene flow, on both a genetic and phenotypic level, can be reduced in sympatry and at secondary contact areas if the diverging species vary in niche or mate preference [7, 15–18]. Areas of elevated genetic differentiation found throughout the genome, so called ‘genomic islands’, or a subset of these regions, could be responsible for the phenotypic differences observed between species or contain potential mechanisms for reproductive isolation and hybrid incompatibility [18–22]. These regions of high genetic differentiation are often spread widely throughout the genome and can be either small [e.g. 23] or large in size [e.g. 24]. As gene flow leaves delectable signatures of divergence in the introgressed regions of genome [1], it is possible to study these patterns and infer past gene flow and also the demographic history of a species [1, 25, 26].

Improved estimation of demographic history makes it possible to better understand population differentiation and speciation mechanisms [e.g. 27]. It allows accurate estimates of gene flow, divergence time, effective population size and population changes in size throughout time. It is also possible to obtain information on which is the most likely demographic history of speciation, such as isolation, isolation with migration, early migration or secondary contact. Using whole-genome approaches also makes it possible to screen for fast evolving regions along genomes that techniques using a small number of genetic markers may miss [e.g. 28]. For example, genomic resequencing in carrion (*Corvus corone*) and hooded crows (*C. cornix*) found that distinct differences in phenotype are maintained by variation in less than 1% of the genome [18]. Finally, population genomics has been shown to provide markedly different estimates of effective population sizes compared to the use of a reduced set of molecular markers and thus could better identify species and populations at risk of extinction, or populations with unique genetic structure worthy of conservation effort. Assessing how secondary contact and hybridisation between distinct taxa impacts native populations is of vital importance when considering how to implement effective conservation protocols [29, 30]. Advancements in analysis on the basis of coalescence simulations [31] using unprecedented high-throughput genomic data of a single individual [32] or populations hold great potential to reveal inference about demographic histories [33].

*Charadrius* plovers are model species for investigating breeding system evolution and have been used in numerous studies to better understand mating and parenting behaviours [e.g. 34–39]. Species in the Kentish plover complex (*Charadrius alexandrinus*; KP) are small shorebirds found breeding on saline lakes and coasts throughout Eurasia and North Africa [40]. A previous study found no genetic differentiation between several Eurasian continental populations of Kentish plover [41]. In East Asia, the subspecies *C. a. alexandrinus* has a wide breeding range in the temperate zone, whereas the southern subspecies *C. a. dealbatus*, (known as white-faced plovers; WFP) show distinct phenotypic traits compared to northern populations. They lack the dark eye barring of the KP, and have lighter lower ear coverts, a brighter cinnamon cap and paler plumage with white lores [42]. They also typically have longer wings, beak and tarsus and are more commonly found on sandier substrates, with more active foraging behaviour and a more upright stance [42]. While the KP and WFP share much of the same wintering range, they have largely non-overlapping breeding ranges. WFPs breed exclusively in a restricted coastal range from Fujian to Guangxi, as well as Hainan island in south-east China. KPs nest to the north of this range [43]. Previous work examining mitochondrial DNA and microsatellite markers has shown that although KPs and WFPs are phenotypically well-differentiated [42, 44], genetically they lack differentiation. More extensive microsatellite genotypes and autosomal nuclear sequences, however, showed KPs and WFPs are distinct and young lineages (diverged around 0.6 mya), and bidirectional gene flow occurred between them [43].

Here, we expand previous works by characterizing demographic histories and genomic landscape of divergence in two closely related plovers. Because the isolation-with-migration model (IM) applied in a previous study assumes gene flow throughout the entire divergence history of the two plover [43], it was unknown whether gene flow persisted in the early stages of divergence or/and also occurred after the secondary contact. The current work explores these different scenarios of gene flow using based on advanced modelling on historical demography. Further, under the model of speciation-with-gene-flow model [10–14], it is of importance to investigate the potential heterogeneous genomic landscape of incipient species. In the case of the two study species, It is possible that a small number of genomic regions involved in the phenotypic and ecological differences among them [28, 42]. Hence, we attempted to disentangle the aforementioned questions by applying whole-genome sequencing and assembly of a high-quality *de novo* reference genome of a female KP. We also re-sequenced whole genomes of 21 unrelated male genomes from five populations of KP and six populations of WFP in China as well as full mitochondrion genomes of four KP and two WFP.

## Materials and Methods

### Sampling collection

A single female KP (heterogametic sex) was collected using mist nets in coastal Xitou, Yangjiang county, Guangdong, China in November 2014. A muscle sample was taken from this individual, stored in RNALater (QIAGEN, USA) and transported to the sequencing centre for *de novo* whole genome sequencing (BGI-Shenzhen). In addition, twenty male KP and WFP were collected from breeding colonies at 11 sites for whole genome resequencing (Figure 1 and Supplementary Table 1), including one inland site at Qinghai Lake, and one continental island, Hainan, with the remaining sites located along the Chinese coast, starting from Hebei to Guangxi. Using males avoids systematic biases occurring caused by differences in coverage of the autosomes and Z chromosome [18]. One female WFP was collected from Hainan for resequencing in higher coverage. These individuals were captured on nests using funnel traps during the breeding season between March and July in 2014-2015 [45]. Blood samples taken from these individuals were stored in RNALater (QIAGEN, USA) at −40°C. All bird captures and sampling was performed with permission from the respective authorities (Beijing Normal University to PJQ and Sun Yat-sen University to YL) and blood and tissue collection procedures conform to the regulations of the animal experimental and medical ethics committee of Sun Yat-sen University.

**Figure 1.**
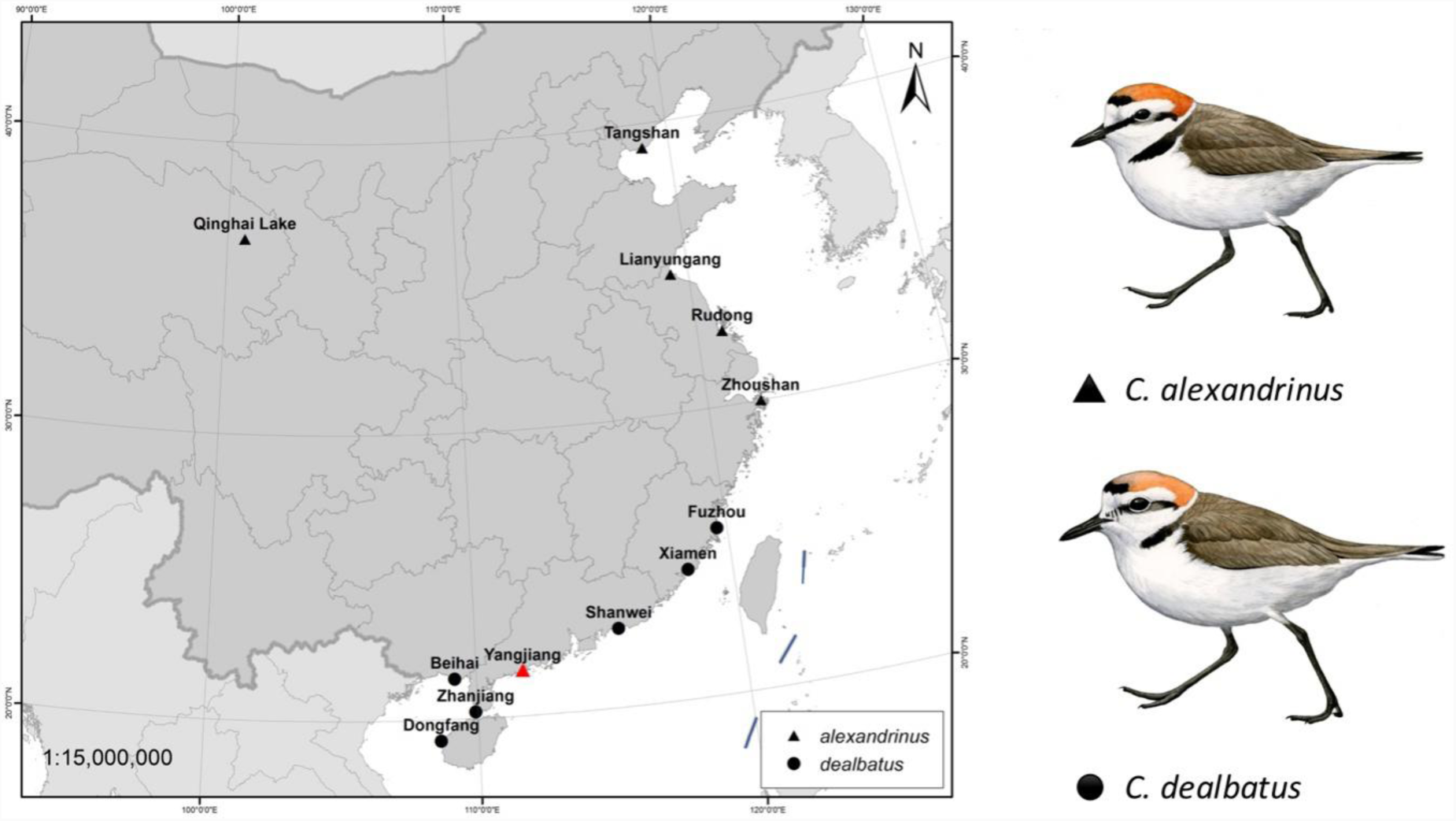
Sampling locations of two plover species, Kentish plover *Charadrius alexandrinus* and white-faced plover *C. dealbatus.* The red triangle represents the location where one individual of Kentish plover for *de novo* sequencing was collected.

### de novo sequencing and assembly of Kentish Plover genome

We isolated DNA from the blood/muscle samples using Qiagen DNeasy Blood and Tissue Kit using the standard manufacturer protocols. Short-insert-sized (170 and 800bp) and mate-pair (2, 5, 10 and 20kb) DNA libraries were constructed for the KP reference genome (Supplementary Table 2). All the libraries were sequenced using Illumina Hiseq 2000 platform on paired-end data. Paired-end sequence data from the genomic DNA libraries were assembled using short oligonucleotide analysis package SOAPdenovo [46]. Final N50 contig and scaffold sizes were calculated based on a minimum length of sequence >100bp. The sequencing coverage, depth and GC content distribution were evaluated by mapping all sequencing reads of the short-insert-sized libraries back to the scaffolds using BWA [47] with the algorithm of BWA-MEM. We also evaluated the genome assembly completeness using BUSCO’s genome mode [48].

### Genome annotation

We combined the homologous prediction method based on the RepBase (http://www.girinst.org/repbase) using the software RepeatMasker and RepeatProteinMask with the *de novo ab initio* prediction method based on self-sequence alignment and repetitive sequence characteristic using the software RepeatModeler (http://www.repeatmasker.org/RepeatModeler), Trf and LTR-FINDER [49]. We used homology, *ab initio* prediction, EST and RNA sequencing to identify protein-coding genes. For homology-based gene prediction, chicken, turkey and zebra finch proteins were downloaded from Ensembl Release 84 (http://www.ensembl.org/info/data/ftp/index.html) and mapped onto the repeat-masked KP genome using tblastn v.2.2.21 [50]. Then we aligned homologous genome sequences against the matching proteins using Genewise (v.wise2.1.23c,10/22/2002) [51] to define gene models. Subsequently, we used the *ab initio* gene prediction software Genescan [52] and Augustus [53] to predict protein-coding genes using parameters trained from a set of high quality homologue prediction proteins. Finally, we carried out RNA-seq on samples of muscle, blood, heart, kidney, liver and brain from the same individual for the *de novo* sequencing. We mapped RNA sequence reads data to the genome using Tophat [54], and obtained transcription-based gene structure using Cufflinks [55]. We also mapped the gene structure and EST sequences to the genome using Genewise. Finally, we merged all genes predicted by the three methods using EvidenceModeler [56], and then removed all genes with a length shorter than 50 amino acids and only with *ab initio* support and FPKM confidence <5 to generate the final gene set. Gene functions were assigned according to the best match of the alignment to the SwissProt, TrEMBL, KEGG and InterPro exogenous protein databases using BLASTP. For the non-coding RNA annotation, we used the software tRNAscan-SE [57] to find the tRNA sequence in Kentish plover genome according to the tRNA architectural feature. Since rRNA sequences are highly conservative, we used similar species rRNA sequences as a reference to find rRNA sequence in the genome using BLASTN. We also used the software INFERNAL [58] with a Rfam family’s covariance model to predict the miRNA and snRNA sequences in the genome.

### Gene family analysis

To define gene families that descended from a single gene in the last common ancestor, we downloaded all protein-coding genes of 15 waterbirds (Emperor penguin, *Aptenodytes forsteri*; Adélie penguin, *Pygoscelis adeliae*; Grey-crowned crane, *Balearica regulorum*; Killdeer, *Charadrius vociferus*; Little egret, *Egretta garzetta*; Sunbittern, *Eurypyga helias*; Northern fulmar, *Fulmarus glacialis*; Red-throated loon, *Gavia stellata*; Crested ibis, *Nipponia nippon*; Dalmatian pelican, *Pelecanus crispus*; White-tailed tropicbird, *Phaethon lepturus*; Common cormorant, *Phalacrocorax carbo*; Greater flamingo, *Phoenicopterus ruber*; Great crested grebe, *Podiceps cristatus*; and Mallard, *Anas platyrhynchos*) from the GigaScience database [59]. Together with the Kentish plover, these species’ genomes represent the major orders in the core waterbird radiation in the avian tree of life [60]. We chose the longest isoform to represent each gene and employed BLASTP to identify potential homologous genes using E-value<1e-10. The raw Blast results were refined using solar (an in-house software) by which the high-scoring segment pairs (HSPs) were conjoined. Similarity between protein sequences were evaluated using bit-score, followed by clustering algorithm in the Treefam pipeline [61].

### Phylogenetic tree construction and divergence time estimate

The protein sequences of one-to-one orthologous genes were aligned using MUSCLE [62] with the default parameters. We then filtered the gap sites from the alignments. The trimmed protein alignments were used as a guide to align corresponding coding sequences (CDS). The phylogenetic tree was reconstructed using RaxML version v8.1.19 [63] and GTRGAMMA model. Divergence times between species were calculated using MCMC tree program implemented in the PAML version 4 [64]. Based on the phylogenetic tree, branch-specific synonymous substitution rate was estimated for KP.

### Whole-genome resequencing

We extracted genomic DNA from blood samples of 20 male individuals of KP and WFP, 10 of each species for 5x depth resequencing and 1 female WFP for 30x depth. For each individual, 1–3 μg of DNA was sheared into fragments of 200–800 bp with the Covaris system. DNA fragments were then treated according to the Illumina DNA sample preparation protocol: fragments were end repaired, A-tailed, ligated to paired-end adaptors and PCR amplified with 500-bp inserts for library construction. Sequencing was performed on the Illumina HiSeq 2000 platform, and 100-bp paired-end reads were generated.

### Read mapping and SNPs calling

After quality control, the reads were mapped to the KP genome using BWA and reads having a mean of approximately 5x depth for each individual and >90% coverage of the KP genome were retained for SNP calling. We used GATK v 3.5 [65] program to call SNPs. SNPs were filtered using VCFtools [66] and GATK by following criteria: 1) missing rate <=0.10; 2) allele frequency >0.05; 3) each 10 bp <=3 SNPs.

### Mitochondrion genome analysis

To infer evolutionary history from mitochondrial DNA (mtDNA), we conducted mitochondrion genome sequencing. Six blood samples were selected, including four KP from four sites (1. Xinbei, Taiwan; 2. Weihai, Shandong; 3. Qinghai Lake, Qinghai; 4. Zhoushan, Zhejiang) and two WFP from two sites (5. Dongfang, Hainan; 6. Minjiang Estuary Fuzhou, Fujian). Gross genomic DNA was extracted by TIANamp Blood Genomic DNA Extraction Kit (TIANGEN, China), following the standard extraction protocol. Paired-end (PE) 150-bp sequence reads were obtained from Illumina MiSeq PE150 sequencing for each sample. Novogene Ltd. (Beijing) performed the library preparation and sequencing. Consequently, we obtained 31,282,372, 29,185,701, 25,092,010, and 24,383,015 clean paired-end reads for the four KP from four sites, respectively; and 30,397,339 and 24,982,573 clean paired-end reads for the two WFP from two sites, respectively. We mapped the clean reads to the mitochondrial genome of the pied avocet, *Recurvirostra avosetta* (GenBank Accession Number: KP757766), using “Map to Reference” tool in Geneious R8 (Biomatters, Auckland, New Zealand) with a medium-low sensitivity and ran 5 iterations. Consensus sequences were saved using a 75% masking threshold, and sites that received insufficient coverage (<5x) were coded using the IUPAC ambiguity symbol N.

We inferred mitochondrial phylogenetic relationship between the two plovers with Bayesian Inference in MrBayes v.3.2.6 [67] and maximum likelihood in RAxML v8 [63] using complete mitochondrial genome sequences including the pied avocet as outgroup. MrBayes was run on the CIPRES science Gateway portal [68] with Metropolis coupling (four chains) set for 10 million generations and sampling every 10000 generations, using HKY nucleotide substitution model which was best-fit model tested by jModelTest 2 [69]. Tracer v1.6 (http://tree.bio.ed.ac.uk/software/tracer/) was used to check the effective sample sizes (ESS) for parameter estimation. RAxML was also run on CIPRES with GTRCAT model and 1,000 bootstrap runs. Maximum-likelihood-bootstrap proportions (MLBS) ≥ 70% were considered strong support [70]. The phylogenetic trees were modified using FigTree v1.4.3 (http://tree.bio.ed.ac.uk/software/figtree).

### Population structure and divergent history between the two plover species

To infer population structuring between KP and WFP, we carried out genetic admixture analysis of the resequenced individuals with ADMIXTURE 1.3 [71]. For K from 1 to 5, each analysis was performed using 200 bootstraps. We applied two approaches to reconstruct the demographic history of KP and WFP. First, we performed a pairwise sequentially Markovian coalescent (PSMC) model to examine changes in historical effective population sizes (*Ne*) of both species [32]. This enabled us to infer demographic dynamics between about 10Ka to 10 Ma. The parameters were set as: “N30 –t5 –r5 –p 4+30*2+4+6+10”, following Nadachowska-Brzyska et al. [72]. Generation time was set to 2.5 years.

The estimated synonymous substitution rate was used as mutation rate, which was 8.11×10^10^ per base pair per year. 100 bootstraps were performed for each analysis. In addition, we carried out a model-based method using an Approximate Bayesian Computational (ABC) approach to infer the divergence history between the two plovers. To achieve this, we first defined four basic demographic models, including: 1) Isolation model, no gene flow during divergence; 2) Isolation with migration model, constant gene flow during divergence; 3) Early migration model, gene flow only exists within the early period of divergence; 4) Secondary contact model, gene flow only exists in the late period of divergence. We performed two groups of simulations with different effective population size (*Ne*) settings (Figure 2). In the first group (A), effective population sizes were hypothesised to be constant in all the four models. In the second group (B), effective population sizes changed based on PSMC results. Model illustrations and priors are shown in Figure 2 and Supplementary Table 17. We used msABC [73] to perform coalescent simulations for these eight models. To obtain the observed data and priors for simulation, 10kbp length loci were randomly chosen from scaffolds with coverage over 80%. The distance between each two loci was higher than 500 Kb to reduce the effect of linkage [27]. Loci with missing data of over 40% in any individual, or over 30% on average, were excluded. *F*_*ST*_, *Tajima’s D* [74], and LD *r*^*2*^ were calculated for each locus with VCFtools 0.1.14. Loci with *F*_*ST*_ higher than 0.159 or *Tajima’s D* >1 or <0 were excluded. 143 10-kbp loci with 14,087 SNPs were left for analysis. 1,000,000 simulations were executed for each model. R package “abc” [75] was used to choose the model which best fitted the observed data with a tolerance of 0.005. Model selection was performed using a multinomial logistic regression method, first in group A and B separately, and then between groups by using two models with highest likelihood in each group. 5,000,000 simulations were used for the best fitted model to estimate population parameters. We used the neural network method with the Epanechnikov kernel to calculate the posterior densities [76]. The number of neural networks was 50. All parameters were log-transformed and medians were used as the as the point estimates.

**Figure 2.**
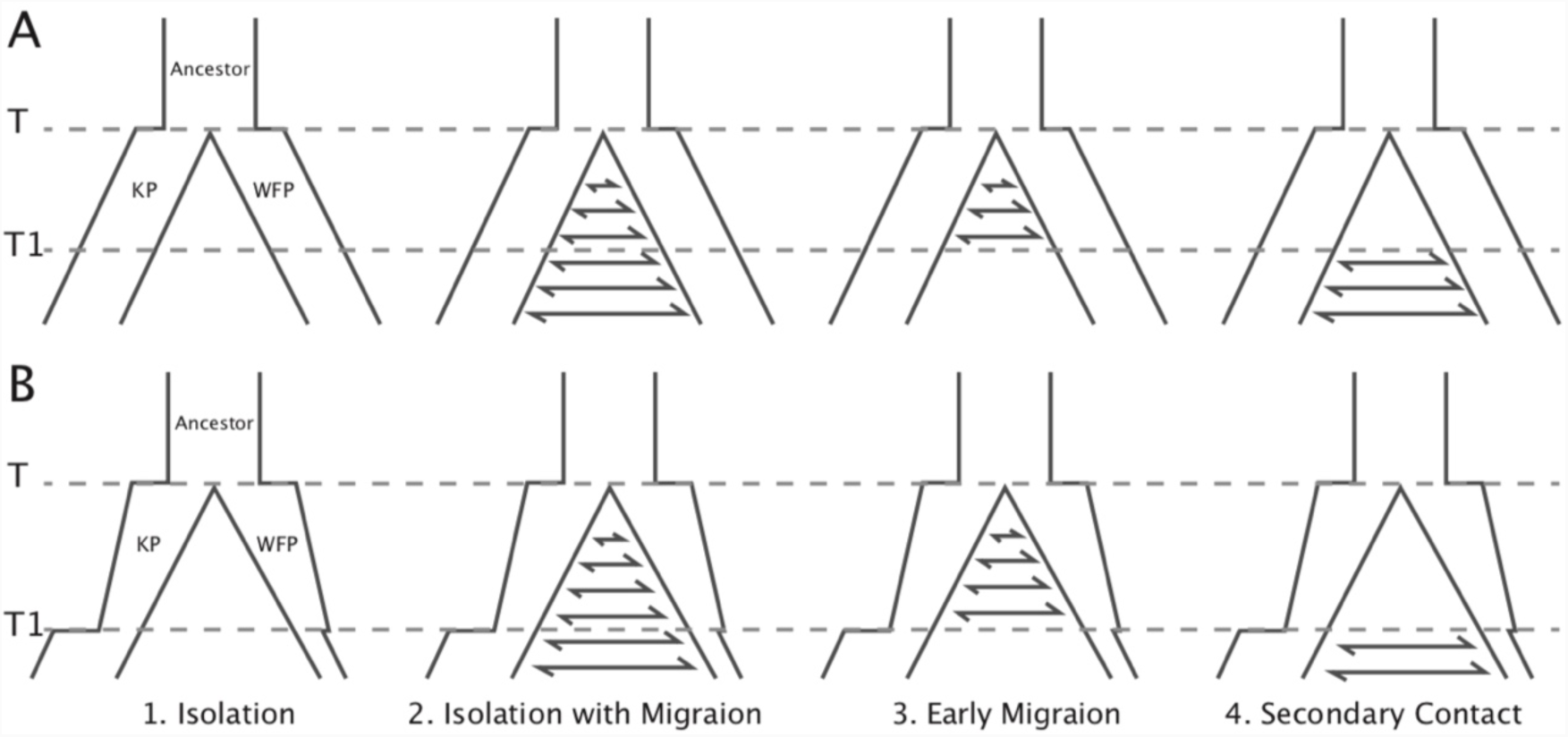
Illustration of the models simulated in ABC analysis. Eight models in two groups were simulated. Effective population sizes in group A were constant. Effective population sizes in group B were based on Ne changes in PSMC. Judged from PSMC, the divergence time was not early than the beginning of population declines 1 million years ago. To simplify the models, population size shifts and changes of gene flow were set to the same time point (*T1*). Prior ranges are available in Table S17.

### Detecting and annotating genomic region under selection

In order to better understand the divergence patterns between the two plover species, genetic parameters *π, Tajima’s D, F*_*ST*_ and *D*_*f*_ (fixed differences) were calculated in 50 kb blocks with VCFtools 0.1.14. Blocks with a length shorter than 25kbp were excluded.

In order to map the KP genome scaffolds onto chromosome coordinates, we downloaded the chicken (*Gallus gallus*) genome from NCBI database (GCF_000002315.4) and computed its alignment with the KP reference genome using Satsuma version 3.1.0 [77]. We divided the genome into non-overlapping windows of 50kb in size with the first window of each scaffold beginning with position 1 of that scaffold, oriented along the chromosome. For each window, we estimated the population genomic parameters calculated above. Finally, we generated the genomic landscape of population divergence in KP and WFP according to the above methods.

To calculate *Tajima’s D* per gene we used the R package PopGenome v2.2.3 [78] in R v3.3.2 [79] for KP and WFP separately. VCF and complementary GFF files were loaded into PopGenome by scaffold with positions with unknown nucleotides excluded (include.unknown=FALSE). Data was then split into genes and neutrality statistics were calculated for coding regions. For calculating *F*_*ST*_, samples were assigned to their two respective species, KP and WFP. *F*_*ST*_ statistics were generated per gene for coding regions.

### Gene ontology annotations

To obtain gene ontology (GO) categories, Kentish plover proteins were BLASTed against the RefSeq protein database using BLASTP v.3.2.0+, with an *E*-value of 1e-5 [80]. GO terms were then assigned using Blast2GO software v.4.1.9 [81] and merged with GO terms obtained from InterProScan v.5.25 (parameters: -f xml, -goterms, -iprlookup) [82]. GO categories were then split into groups associated with biological processes and cellular components. GO categories with fewer than 50 genes were grouped into a single “small” category. Genes with no GO annotation were assigned to an “uncharacterised” category.

### Gene ontology enrichment analyses

Genes were assigned to either autosomes or the Z chromosome, with genes of unknown location excluded from GO enrichment analysis. GO enrichment was performed for genes with high *F*_*ST*_. Genes were considered to have high *F*_*ST*_ if they fell above the 95^th^ percentile of *F*_*ST*_ values, using R quantile(c(.95)). Two cut-off values were used, 0.184 for autosomal genes and 0.262 for genes on the Z chromosome. GO enrichment was also performed on genes with high and low *Tajima’s D* values calculated for KP and WFP separately. Genes with positive *Tajima’s D* could indicate balancing selection. Regions of negative *Tajima’s D* can indicate strong positive selection or selective sweeps. Genes were separated into known autosomal and Z chromosomal genes for positive and negative *Tajima’s D* separately. The 95^th^ percentile of positive values were taken as high *Tajima’s D* genes and the 5^th^ percentile were taken for the negative values. This resulted in cut-off values of ⩾ 1.565 for high and ⩽-1.179 for low value autosomal genes for KP and ⩾ 1.669 and ⩽-1.191 for WFP. ⩾ 1.723 and ⩽-1.234 cut-offs were used for the Z chromosome for KP and ⩾ 1.835 and ⩽-1.445 for WFP. GO enrichment was performed for both high and low *Tajima’s D* values separately. GO enrichment analyses were performed to evaluate if any GO category was over represented in the set of genes of interest compared to equally sized samples of genes drawn randomly. The expected number of genes annotated to each GO was calculated using 1000 equally sized random families. Significance was established using Z-scores and a Benjamini-Hochberg correction was applied to adjust for multiple comparisons. GO categories were significantly enriched if the adjusted p value was < 0.05. Results were filtered to remove any category with an expected number of genes per category of < 1 and an observed number of GOs of 1. This analysis was performed using R v.3.3.2.

## Results

### De novo sequencing the Kentish plover genome

Muscle samples from a heterogametic sex female Kentish plover were collected from a wintering population in Guangdong, China (Figure 1). Short read DNA sequencing (125bp) was carried out using the Illumina platform (see Supplementary Figure 1 for pipeline). After filtering out low quality and clonally duplicated reads, we obtained a genome assembly from 1.81 billion reads in six paired-end and mate-pair libraries that provide 134-fold coverage with a total assembly length of 1.16Gb (Supplementary Table 2). This approximates the genome size estimated using K-mer frequency method (Supplementary Table 3 and Supplementary Figure 2). The GC content versus depth is gathered into a cluster showing the genome sequence is pure and has no pollution from other species (Supplementary Figure 3.

**Figure 3.**
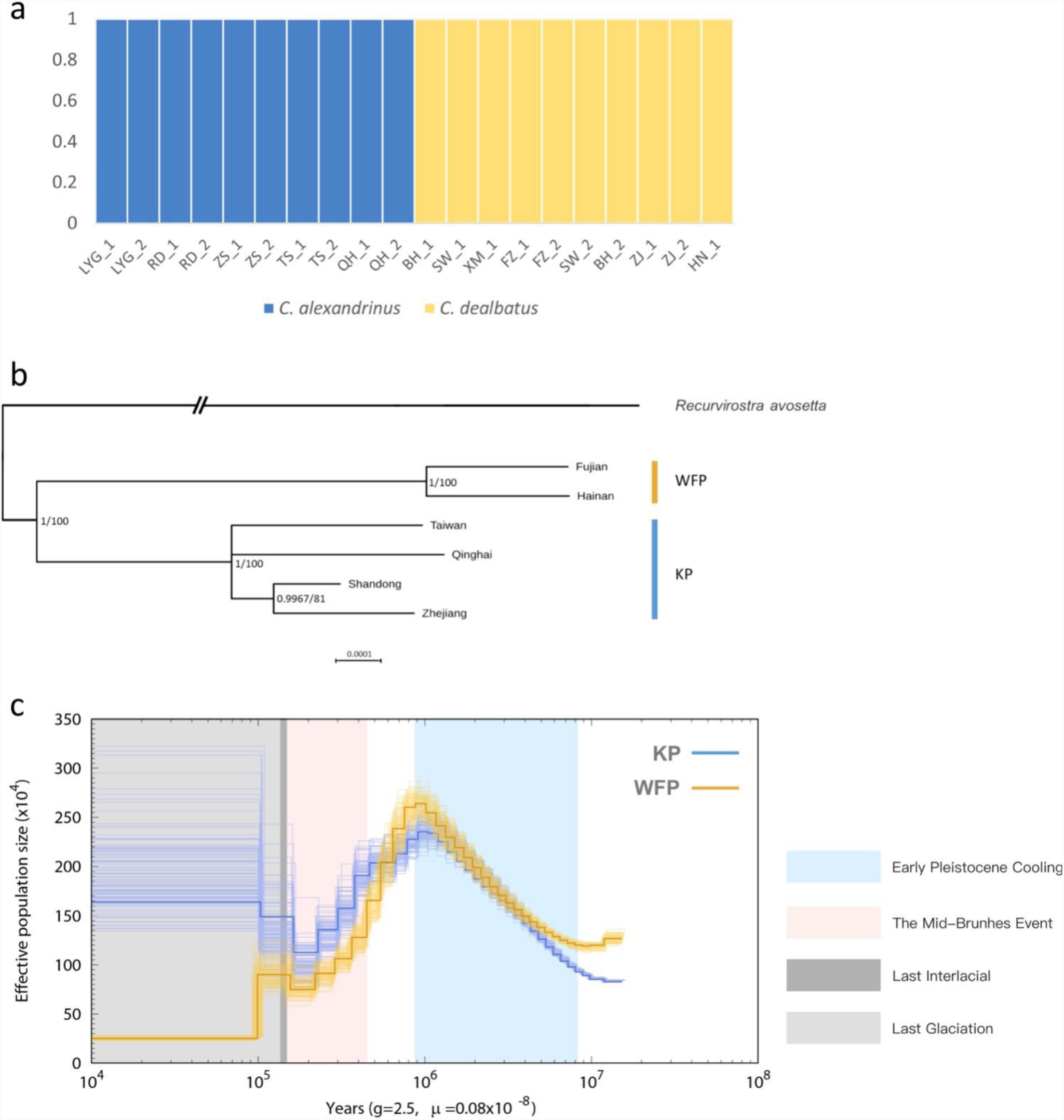
Population genetic structure and historical demography. *C. alexandrinus* marked in blue and *C. dealbatus* in yellow. a) Genetic clustering inferred with ADMIXTURE when K=2. b) Phylogenetic relationship between the *C. dealbatus* (WFP) and different populations of *C. alexandrinus* (KP) using Bayesian and Maximum Likelihood methods based on mitochondrial genome sequences (c.a. 15kb). Posterior probabilities (pp) and bootstrap supports are indicated at each node. White-faced plover and Kentish plover form two independent evolutionary lineages. c) Demographic history of the Kentish plover, blue line, and white-faced plover, yellow line reconstructed from the reference and population resequencing genomes. The line represents the estimated effective population size (*N*e), and the 100 thin blue curves represent the PSMC estimates for 100 sequences randomly resampled from the original sequence. Generation time (*g*) = 2.5 years, and neutral mutation rate per generation (μ) = 8.11 × 10^−10^. The Last Interglacial period (LIG, from approx. 130 to 116 ka) is marked by a grey block.

The contig and scaffold N50 sizes are 38.9 and 3220.7kbp, respectively, with the largest scaffold spanning 15291.1kbp (Supplementary Table 4). Although the number of scaffolds for the Kentish plover is considerably higher than that of the chicken or the zebra finch genomes, the estimated genome size for the Kentish plover (1,245,524,081bp, ~1.25 Gb) is comparable to the sequenced genomes of these two species (Supplementary Table 5). Whole genome alignment reveals that, as expected, a higher proportion of the zebra finch and KP genomes can be aligned against each other than either of them can to the more distantly related chicken with over 900Mbp that can be aligned between the two species (Supplementary Table 6). The BUSCO assessment results indicate that the KP genome assembly has high completeness (C: 94%) (Supplementary Figure 4).

**Figure 4.**
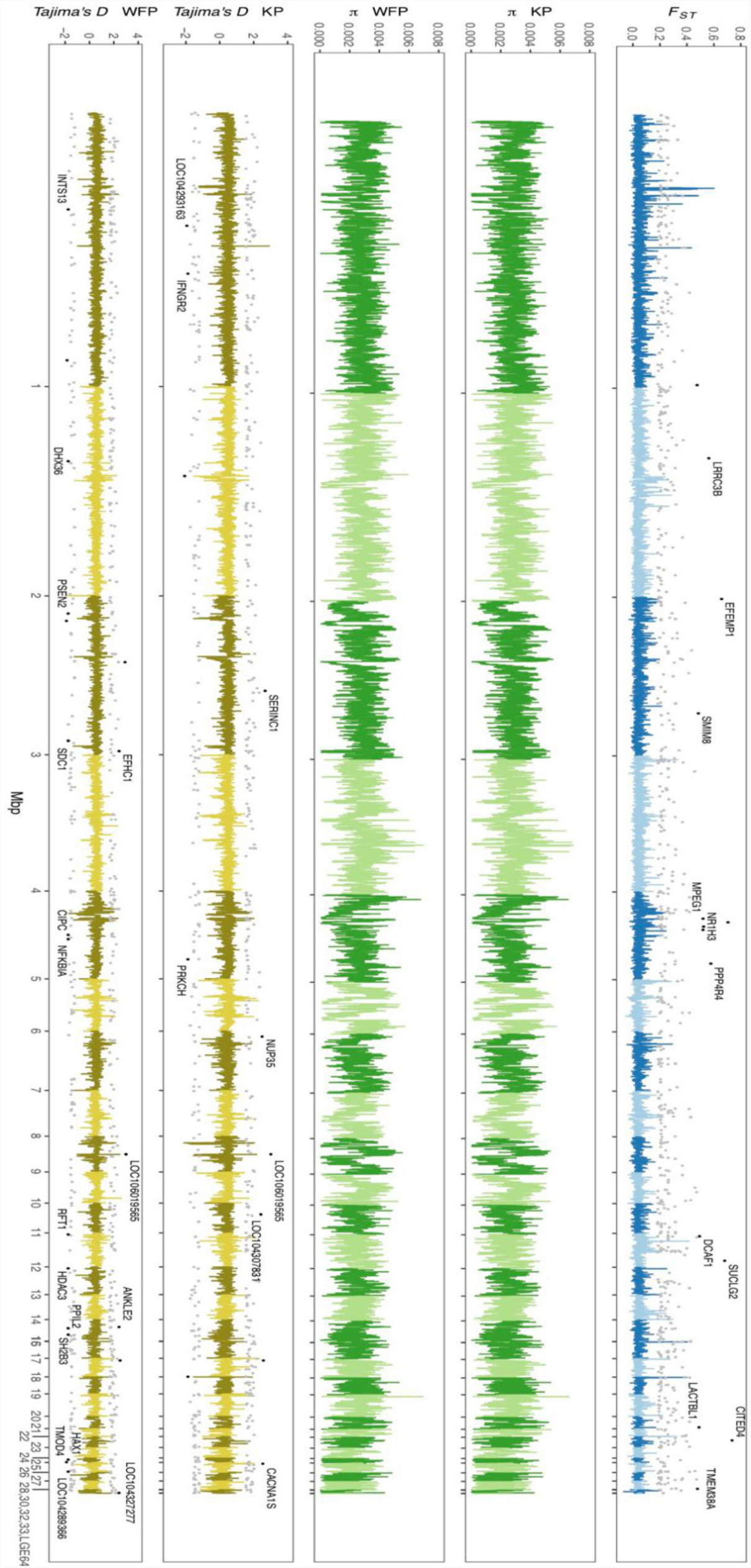
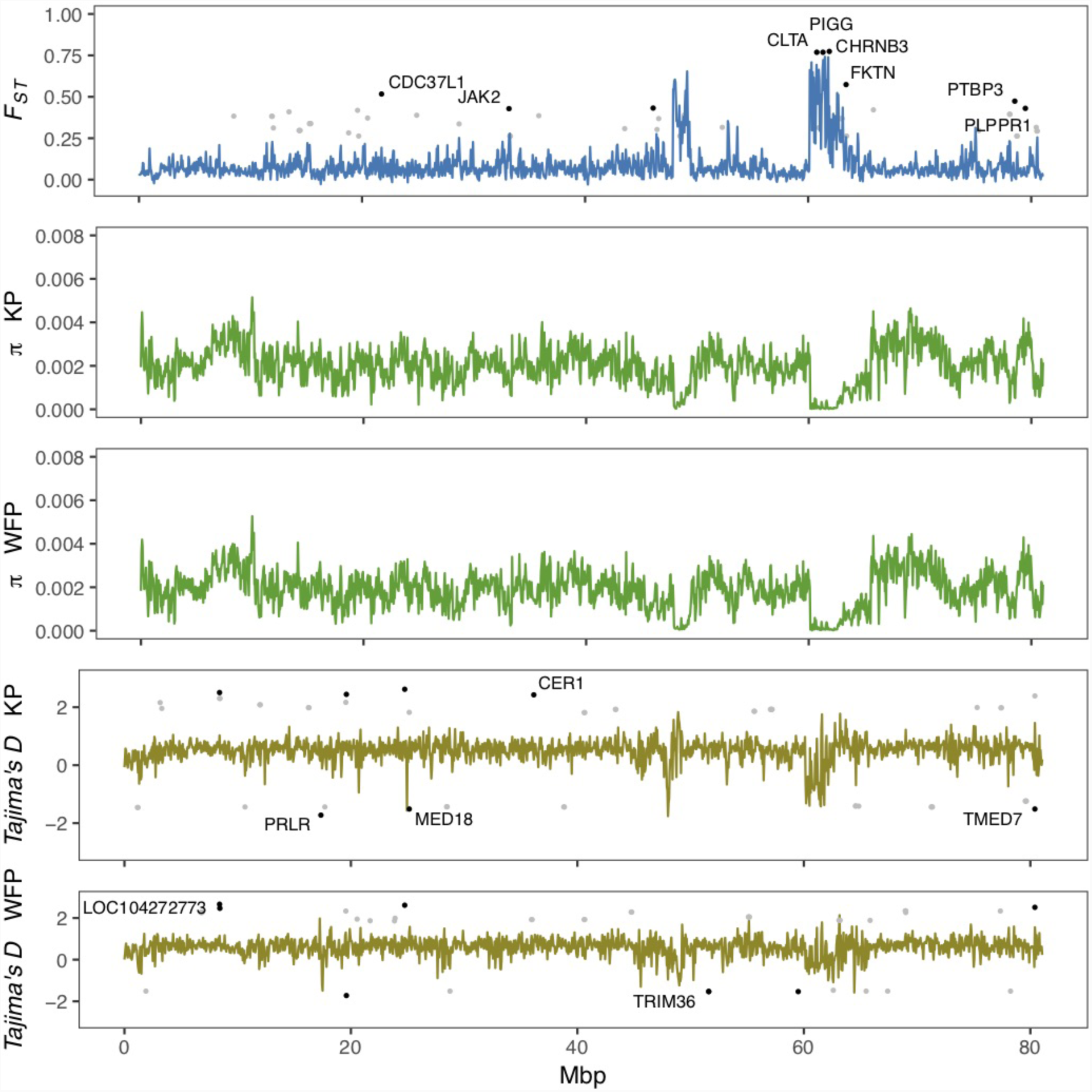
Genome wide landscape of *F*_*ST*_, *π*, and *Tajima’s D* for 50 kb sliding window. Different autosomes are marked with alternating light and dark colors. **a**) Genome wide landscape for autosomes. 95^th^ percentile outliers are plotted for *F*_*ST*_ and 95^th^/5^th^ percentile outliers are plotted for *Tajima’s D* in grey, as calculated per gene. 99.9^th^ percentile outliers for *F*_*ST*_ and 99.9^th^/0.1^th^ percentile outliers for *Tajima’s D* are plotted in black per gene and labelled with gene symbols. **b**) Genome wide landscape for Z chromosome. 95^th^ percentile outliers are plotted for *F*_*ST*_ and 95^th^/5^th^ percentile outliers are plotted for *Tajima’s D* in grey, as calculated per gene. 99^th^ percentile outliers for *F*_*ST*_ and 99^th^/1^st^ percentile outliers for *Tajima’s D* are plotted in black per gene and labelled with gene symbols.

Within the genome sequence of the KP, fewer than 4% of the bases were N bases (Supplementary Table 7). We estimate that repetitive sequences compose about 10% of the genome (Supplementary Table 8) with LINE transposons being the most common, making up about 8% of the genome. LTR transposons are the second most common repetitive sequences in the Kentish plover genome with 2.22% occupancy, with other DNA and LINE transposable elements occupying less than one per cent of the genome (Supplementary Table 9). Annotation of non-protein coding RNA genes revealed around 722 non-coding genes with transfer RNAs and small nuclear RNAs being the most common with over 200 copies each (Supplementary Table 10). Based on the number of RNA genes identified in heavily annotated genomes such as human and mouse, it is likely more will be found in the future as non-coding gene annotation tools improve further.

Initially, annotation of protein coding genes identified 8,893 genes. Further annotation based on homologous sequence alignment against the chicken, the zebra finch and the turkey aided to identify a final set of 15,677 genes in the Kentish plover genome (Supplementary Table 11). The quality of protein coding gene annotation is of similar quality to that of other avian genomes (Supplementary Figure 5). Of these genes, based on ortholog annotations, the vast majority of protein coding genes (15,644) were annotated against at least one functional category in one of several functional annotation databases (Supplementary Table 12).

Protein coding gene content was then compared to that of 15 previously sequenced waterbird species’ genomes. Gene number identified ranged from 13,454 in the Gaviiformes to the egret with 16,585 annotated protein coding genes, with the Kentish plover having a number close to the average (Supplementary Table 13). The number of genes found in the KP genome are close to the average number of genes found in other waterbird species (Supplementary Table 13). Most KP genes have homologs in all other shorebird genomes it was compared to (12,196/15,677; Supplementary Figure 6). Single copy orthologs make up almost 40% of the genes with other orthology relationships making up most of the rest of the genes. Less than 5% of genes did not cluster into orthology groups (Supplementary Figure 7).

Phylogenetic reconstruction from coding sequence alignment places the KP as a closer relative of a previously sequenced plover genome of the killdeer *Charadrius vociferous* than to any other waterbird genome and with the most recent divergence time, as expected (Supplementary Figure 8; Supplementary Figure 9).

### Genome resequencing reveals two diverging plover species with contrasting evolutionary histories

Blood samples were obtained from a total of 21 individuals taken from mainland and Hainan island plover populations along the Chinese coast and from the inland and a high-altitude population of Qinghai Lake (Figure 1 and Supplementary Table 1). Kentish and white-faced plovers have been shown to present distinct phenotypic features, including facial plumage pattern (Figure 1) [44]. For the 20 low depth resequenced samples, a total of 914,529,390 high quality paired-end reads were retained for the further analyses (Supplementary Table 14). Genome resequencing was carried out resulting in over 95% genome coverage with a depth of over 4x for around 70% of the genome per individual (Supplementary Table 15). After filtering, a total of 11,959,725 high quality SNPs were retained. KP and WFP were found to cluster into two distinct groups based on Admixture analysis (Figure 3a). For the high depth sequencing WFP, its genome assembly quality was high enough to be used for the PSMC analysis.

In addition, we obtained 15,613 bp of the complete mitochondrial genome for Kentish and white-faced plovers, except for the D-loop region, which had poor assembly quality. In the phylogenetic analysis, topologies between Bayesian and ML tree were consistent (Figure 3b). Analyses clearly show that the monophyletic relationship of KP and WFP is strongly supported.

Demographic history reconstruction of KP and WFP revealed distinct evolutionary histories of the two species from approximately 10 Ma to 10 ka (Figure 3c). PSMC analysis demonstrated a similar *Ne* history for both species around 1 million years ago during the glaciation, with population sizes of both species rising from 0.8 to 1 million years ago then sharply declining until about 100 thousand years ago during the Last Interglacial Period. KP and WFP then went through steady population size changes separately. The *Ne* of KP increased greatly to 1.65 million and the *Ne* of WFP decreased to about ten thousand.

With ABC simulations, we found that the genomic-wide polymorphism patterns in KP and WFP fit best with the secondary contact model (posterior probability = 0.98, Supplementary Table 17), suggesting that the two plover species experienced gene flow after secondary contact. Model selection showed that changing *Ne* models incorporating the PSMC-inferred Ne fluctuations had much higher posterior possibilities than constant *Ne* models (Bayes factors > 10^3^), which indicates that the secondary contact model based on PSMC results fitted best (Figure 2).

Demographic analyses allowed us to estimate several demographic parameter estimates, including divergence times, effective population sizes and migration rate per generation (Table 1). The two plover populations are estimated to have diverged approximately 606 000 years ago (95% CI, 277 000 ~974 000 years). The effective population size of KP is estimated to be around 2.6 times than that of WFP (median *Ne*_K_ = 296,800 and median *Ne*_W_ = 114,500). The most recent common ancestor is believed to have an effective population size, *Ne*_A_, which is about 10-15 times greater than both modern species. Gene flow between the two groups of plover was approximately symmetric: c. 7.85 individuals per generation immigrate from WFP to KP, compared with c. 3.21 individuals per generation immigrating from KP to WFP. It must be noted however, that the ranges of the probability distributions for migration estimates are broad (Supplementary Figure 10).

**Table 1.** Posterior median, mean, mode and range of 95% highest probability distribution (HPD) of demographic parameters. K represents Kentish Plover *Charadrius alexandrinus* and W represents White-faced Plover *C. dealbatus*. A represents Ancestral population. Ne: recent effective population size; T: population split time; M: migration rate per generation.

|  | **T/10<sup>4</sup>** | **Ne<sub>K</sub>/10<sup>4</sup>** | **Ne<sub>W</sub>/10<sup>4</sup>** | **Ne<sub>A</sub>/10<sup>4</sup>** | **2NM<sub>W→K</sub>** | **2NM<sub>K→W</sub>** |
| --- | --- | --- | --- | --- | --- | --- |
| Median | 60.60 | 29.68 | 11.45 | 72.15 | 7.85 | 3.21 |
| Mean | 61.20 | 137.09 | 20.02 | 98.71 | 7.62 | 3.11 |
| Mode | 54.10 | 16.01 | 5.62 | 26.74 | 7.31 | 3.27 |
| 2.5% HPD | 27.70 | 5.22 | 3.33 | 13.35 | 3.07 | 1.84 |
| 97.5% HPD | 97.40 | 1262.12 | 99.91 | 416.42 | 11.32 | 4.03 |

### Genomic regions associated with divergence between Kentish plover and White-faced Plover

Genome scans showed low genome-wide divergence between KP and WFP (Figure 4). The genome-wide *F*_*ST*_ was 0.046, *π* of KP was 0.00262 and *π* of WFP was 0.00259. 24,065 blocks of 50kbp length were scanned, 22,148 of which were located on autosomes, 1,622 were located on Z chromosome and 295 unassigned. Since the W chromosome is very short (5.16Mb in chicken genome) and only males were used for population analyses, it was not included in the genomic landscape analysis. Examining autosomes and the Z chromosome separately, it was found that the average *F*_*ST*_ of autosomal blocks was 0.043, *π* of KP was 0.00268 and *π* of WFP was 0.00266. The top 1% outlier blocks on autosomes, which have the highest *F*_*ST*_ (average 0.230, peak 0.605), were found to have much lower polymorphism, than when examined at the genome-wide scale (*π*_KP_ = 0.0012, *π*_WFP_ = 0.0011, p < 0.001). The Z chromosome was more divergent than autosomes (average *F*_*ST*_ = 0.089, *π*_KP_ = 0.0020 and *π*_WFP_ = 0.0019). The highest 1% outlier blocks on the Z chromosome had an average *F*_*ST*_ of was 0.664, and the peak value was 0.741. The average *π* was 0.0003.

We identified genes which had high levels of divergence between species, by calculating population statistics per gene (Figure 4). 691 autosomal genes had *F*_ST_ higher than 0.184 and 41 genes on the Z chromosome had *F*_*ST*_ higher than 0.262 (Supplementary Table 18). GO enrichment analyses of autosomal genes with high *F*_*ST*_ found that no categories were enriched for biological processes and integral component of membrane was enriched for cellular components. No categories were enriched when looking at the genes with high *F*_*ST*_ on the Z chromosome. A total of 339 autosomal genes were found with a *Tajima’s D* ≥ 1.565 (Supplementary Table 19) and 235 genes with *Tajima’s D* ≤ −1.179 for KP (Supplementary Table 20). No GOs were enriched after GO enrichment analysis. 325 genes had values of *Tajima’s D* ≥ 1.669 (Supplementary Table 21) and 216 had values of *Tajima’s D* ≤ −1.191 for WFP (Supplementary Table 22). GO enrichment analysis revealed an overrepresentation of high *Tajima’s D* genes associated with microtubule cellular component categories and genes with low *Tajima’s D* had an overrepresentation of genes associated with proteolysis for biological function categories. For KP 18 genes on the Z chromosome had a *Tajima’s D* ≥ 1.723 and 12 genes had *Tajima’s D* ≤ −1.234. GO enrichment was not performed due to the small number of high diversity genes on the Z chromosome. WFP had 19 genes on the Z chromosome had *Tajima’s D* ≥ 1.835 and 10 genes had *Tajima’s D* ≤ −1.445.

## Discussion

We have utilised methods of mitochondria and nuclear whole genome *de novo* sequencing to shed unprecedented insight into the evolution and demographic histories of two small plover species. Although previous studies suggest KP and WFP are sufficiently differentiated at a phenotypic level to justify them as two different species [42, 44], genetic studies have not always agreed [42]. Our results show that KPs and WFPs have low levels of differentiation on average at the genomic level, with moderate to high differentiation in some regions. Our divergence time estimates suggest that Kentish and white-faced plovers diverged relatively recently, less than 0.61 million years ago.

Using genome wide data, we were able to model the demographic histories of the two plovers. By using more complex demographic models it was possible to model demographic histories more comprehensibly than would be possible using traditional population genetics techniques. ABC analysis estimates the divergence time to be about 606,000 years ago during the Pleistocene. Estimates of Ancestral *N*_*e*_ suggests that the most recent common ancestor had a much higher *N*_*e*_ (721,500) than either modern species. Current *N*_*e*_ estimates for KP (296,800) were about 2.5 times that that of WFP (114,500). *N*_*e*_, can be important as an estimate of the health of a population in terms of conservation biology. This can be particularly valuable for species where census data is lacking, such as with the WFP which has no census data available from the IUCN Red List, although the relationship between *N*_*e*_ and census *N* can be difficult to interpret.

We found contrasting demographic histories between the two plover species as demonstrated from the PSMC estimates, especially through their history from 1 million to approximately 100 thousand years ago. The *N*_*e*_ of KP was about seven times larger than WFP towards the end of the last glacial period (LGP) resulting from population increases in KP and declines in WFP. Although declines in population size at the start of the LGP is a common pattern in bird species [72], marked contrasting patterns in population demography may reflect species differences in response to historical climate fluctuation and also population divergence [83]. One possibility is that a decrease in suitable habitat for WFP during the LGP could explain continued declines of the *N*_*e*_ of WFP whereas the KP may have been able to exploit a wider range of habitats. Caution should be exercised when implementing demographic approaches such as ABC and PSMC and when interpreting the results, due to heavy dependence on the parameters and priors [27, 84].

It is unknown whether the breeding grounds of KP and WFP overlap or whether any form of reproductive isolation has occurred. If a hybrid zone exists it has been estimated that this would be found in a narrow range in Fujian Province [42]. It is believed that advanced reproductive isolation occurs relatively late during speciation events in birds, with complete F_1_ hybrid sterility often taking in the order of millions of years [27, but see 85]. The recent split of the two plover species would suggest that reproductive isolation may be limited. The rate of gene flow and recombination occurring throughout the genome can affect the rate at which reproductive isolation occurs. Migratory shorebirds are highly mobile with the potential to disperse large distances to breed, this can lead to high levels of gene flow and weak population structuring in some species [41, 86–88]. We estimated levels of gene flow throughout the history of the two species to determine whether historic gene flow occurred as a result of secondary contact or alternatively that gene flow has occurred continuously as the species diverged. Various scenarios of gene flow were modelled using ABC analysis. ABC simulations best fit the secondary contact model, in which the two plover species diverged allopatrically, with gene flow occurring after secondary contact. There was little support for models of isolation with migration, isolation, or early migration models. Gene flow was found to be approximately symmetric and bidirectional between the two species. How both phenotypic and genomic differentiation is maintained in spite of gene flow is a key question in evolutionary biology. We report levels of gene flow with more than five individuals introgressing between the two species per generation. It is expected that very few individuals are needed to exchange genes between populations to break down differentiation produced by genetic drift [89, 90]. This suggests that some form of selection is acting on these two species to maintain this divergence within the populations. Further study of these species at the hybrid zone would help elucidate which selective forces might be maintaining these differences.

Despite secondary gene flow, we detected highly differentiated genomic regions that may contribute to species divergence [18, 22, 91–93]. We used a window based approach to perform whole genome scans and calculate various population statistics, including *F*_*ST*_, *π* and *Tajima’s D*, to detect areas of high divergence and selection. We found that average *F*_*ST*_ across genome was 0.046. This is slightly higher than levels found between carrion and hooded crows, another taxonomically debated pair of species [18]. It has been suggested that areas of peak divergence contain genes involved in reproductive isolation and that these areas can contain genes responsible for differences in phenotype [18–22]. Higher levels of *F*_*ST*_ and *Tajima’s D* in KP were found in the PPP3CB gene, part of the oocyte meiosis KEGG pathway and could therefore be important for promoting genetic incompatibility in females. Hybrid incompatibility often the heterogametic sex due to Haldane’s Rule [94], which often means in birds that hybrid females are sterile and fertile males allow gene flow to occur between species [95]. The Z chromosome had higher levels of block-average and peak divergence, 0.089 and 0.741 respectively. This pattern is consistent with results from previous studies [18, e.g. 22]. Mean divergence levels of sex chromosomes are often higher than on the autosomes. This could be caused by the lower *Ne* of the Z chromosome compared with the autosomes, which can result in faster lineage sorting and higher *F*_*ST*_ [9]. There is often an overabundance of highly differentiated loci on the sex chromosomes [1]. In light of this pattern, Cruickshank and Hahn [96] proposed that speciation is not necessarily the sole force contributing to divergence in genomic regions. Population bottlenecks, i.e. in WFP (Figure 2c), and divergence in regions of low *Ne*, i.e. Z chromosome, can also contribute to heterogeneity of genomic differentiation [9].

This study also provides a greater understanding about the demographic histories of two shorebird species, especially linking their current population status with evolutionary context [29, 30, 97]. Although KP have a large census population size and are widely distributed throughout Eurasia and North Africa, there is evidence that populations are in decline in East Asia. The decline in Chinese plovers could, at least in part, be due to a reduction in suitable breeding and feeding locations along the Chinese coast due to land reclamation and development [98]. Extremely low nest survival in Kentish plovers from Bohai Bay has been reported and linked to anthropogenic disturbance [99]. This work also emphasises the lower effective population size of WFP when compare to KP, which has resulted from population declines since the LGM (Figure 2c). Thus, increased effort to monitor population trends for this species is warranted in order to accurately assess any potential threats to this species and for conservation status evaluation for the IUCN.

In conclusion, we produced the first high quality genome of the KP and performed whole genome resequencing of two plover species relatively early in their divergence. We found multiple pieces of evidence to support that the WFP and KP are distinct lineages with complex demographic histories. We found evidence for gene flow between these two species due to secondary contact. Our results further reveal a heterogeneous pattern of genomic differentiation with elevated divergence in the Z chromosome. This suggests that some form of selection is working to maintain genetic and phenotypic differences between the two species. Overall, this study provides new insights into the genomic patterns between a species pair at an early stage of speciation. Further analyses of populations at the hybrid zone would increase our understanding of the specific selective forces maintaining this divergence.

## Acknowledgements

The authors thank Qiaoyi Liang, Zhechun Zhang, Xuecong Zhang, Xin Lin and Demeng Jiang for their assistance during sampling, and Shaochong Peng and Chunfa Zhou for their help in the preparation of Figure 1. This work was supported by National Natural Science Foundation of China (31301875, 31572251 to YL, and 31600297 to PJQ) and Special Program for Applied Research on Super Computation of the NSFC-Guangdong Joint Fund (the second phase) under Grant No. U1501501; a National Environment Research Council Great Western Four+ Doctoral Training Partnership studentship (grant number NE/L002434/1), Korner Travelling Award and a British Council and Chinese Scholarship Council Newton Fund PhD Placement awarded to KHM; a Royal Society Dorothy Hodgkin Research Fellowship (grant number DH071902), Royal Society research grant (grant number RG0870644), a Royal Society research grant for fellows (grant number RG080272) and a NERC grant (NE/P004121/1) to AOU. All sequencing data will be deposited in NCBI databases upon acceptance.

## Data Availability

This information will become available upon the acceptance of the manuscript.

## Author Contributions

YL and TS conceived of the study. XJW and YL designed the study. PQ, QH, BW, ZWZ and YL collected the samples. XW, NZ and KHM, CZ, SL, BW, DC, XY analysed data. KHM, AOU and YL wrote the manuscript with contributions from all authors. All authors read and approved the final version of the manuscript.

## Supplementary Material

Supplementary information, Supplementary Figures 1-10 and Supplementary Tables S1-S22 are available online.

