## Supplementary information for "Demographic histories and genome-wide patterns of divergence in incipient species of shorebirds"

**Running title:** Population genomics of two plover species

**Abbreviation：**

KP：Kentish plover (*Charadrius alexandrinus*)

WFP: White-faced plover (*Charadrius dealbatus*)

**Supplementary Table 1** Sampling sites of KP and WFP

| Species | site | amount |
|---|---|---|
| **Kentish plover** | Yangjiang, Guangdong* | 1 |
| | Qinghai Lake, Qinghai | 2 |
| | Tangshan, Hebei | 2 |
| | Lianyungang, Jiangsu | 2 |
| | Rudong, Jiangsu | 2 |
| | Zhoushan, Zhejiang | 2 |
| **White-faced plover** | Minjiang Estuary, Fuzhou, Fujian | 2 |
| | Xiamen, Fujian | 1 |
| | Shanwei, Guangdong | 2 |
| | Zhanjiang, Guangdong | 2 |
| | Beihai, Guangxi | 2 |
| | Dongfang, Hainan | 2 |
| **Total** | | 22 |

* The wintering female individual used for *de novo* sequencing.

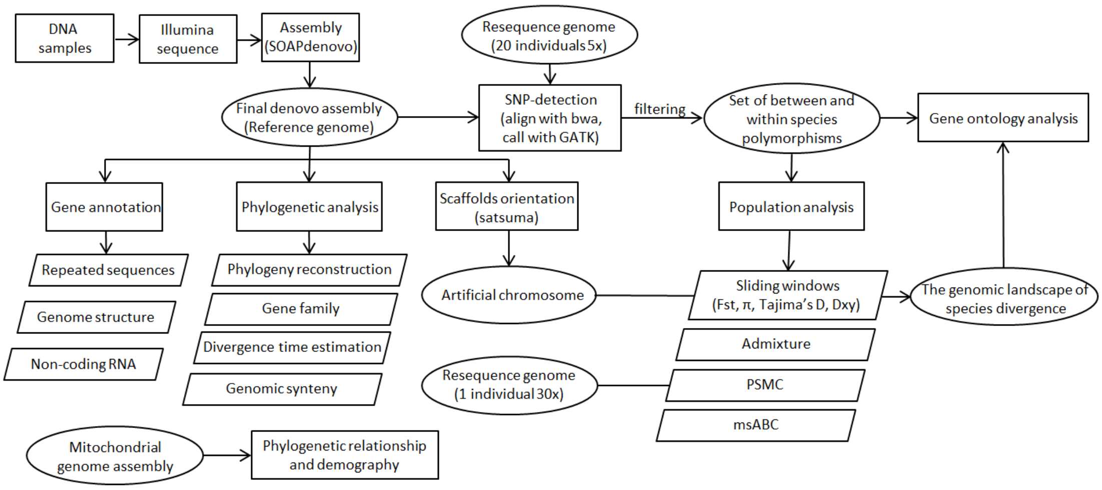


**Supplementary Figure 1** Schematic overview of the different analyses of the study

### 1. Genome sequencing and assembly

#### 1.1 *De novo* sequencing

**Supplementary Table 2** Summary of DNA libraries and sequencing data of the KP genome

| Library insert size | Read Length (bp) | Data (Gb) | Sequencing Depth (×) | Physical Depth (×) |
|---|---|---|---|---|
| 170bp | 125 | 54.99 | 40.72 | 27.69 |
| 800bp | 125 | 24.68 | 18.28 | 58.49 |
| 2kb | 125 | 30.25 | 22.4 | 179.21 |
| 5kb | 125 | 23.32 | 17.27 | 345.32 |
| 10kb | 125 | 24.57 | 18.19 | 727.62 |
| 20kb | 125 | 23.82 | 17.64 | 1411.54 |
| Total | | 181.63 | 134.5 | 2749.86 |

### 1.2 Quality control of raw sequencing reads

To prepare high quality data for *de novo* genome assembly of KP genome, the raw sequencing data meeting the following conditions were filtered using a combined strategy.

1) Reads in which N constitutes more than 2% (for the short-insert libraries), 5% or 10% (for the mate-paired libraries) of read length or polyA structure reads.
2) Low quality reads. Reads of short-insert libraries that have 40% bases with quality scores ≤7; reads of mate-paired libraries that have more than 30% or 40% bases with quality scores ≤7.
3) Reads with adapter contamination. Reads with more than 10 bp aligned to the adapter sequences (allowing less than or equal to 3 bp mismatches).
4) Short insert-size libraries (250 bp, 500 bp, 800 bp insertion size) in which forward and reverse reads overlapped >=10 bp allowing 10% mismatches and Read1 and Read2 are both ends of one paired end reads.
5) PCR duplicates.
6) The raw reads were also corrected based on K-mer spectrum.

Finally, 181.63 Gb (around 134.5x coverage of KP genome) reads were obtained for following procedures.

### 1.3 Estimate the Genome Size using K-mer spectrum

The genome size of KP was estimated around 1.35 Gb based on K-mer spectrum [1]. More details are available in Supplementary Table 3.

**Supplementary Table 3** Estimated genome size of the KP genome and parameters related to sequencing quantity and depth.

| K-mer | K-mer Number | Peak Depth | Genome Size (bp) | Used Bases (bp) | Used Reads | depth (×) |
|---|---|---|---|---|---|---|
| 17 | 54,018,385,172 | 40 | 1,350,459,629 | 63,729,555,540 | 606,948,148 | 47.19 |

### Supplementary Figure 3

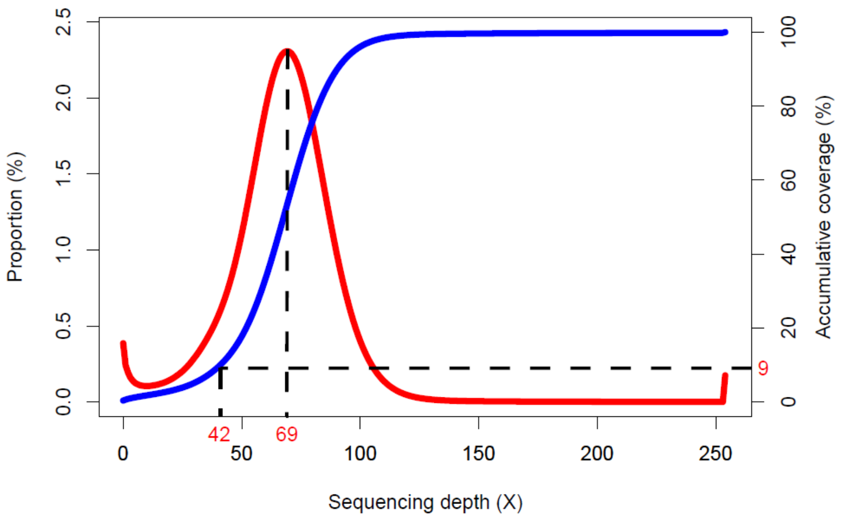


**Supplementary Figure 2** The depth coverage of the KP genome assembly

The cleaned reads from 170 bp, 800 bp libraries were mapped to the genome assembly using bwa. The X-axis is the depth of coverage; red curve indicates the percentage of depth of coverage (left Y-axis) and blue curve is the accumulated percentage of depth of coverage (right Y-axis). From the figure, we can see the average depth coverage is around 69 and 90% of the bases' sequencing depth larger than 42. This indicates the assembly result has a high sequencing depth.

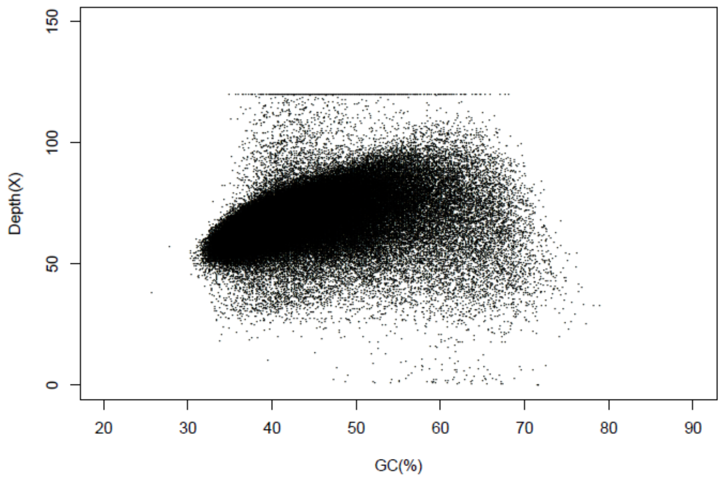


**Supplementary Figure 3** Depth versus GC content of assembled genome sequence of KP genome

We scanned the KP genome with a sliding window of a size equal to the mean fragment length and with the step size set as half the window size. In each window, we calculated the GC content (the percentage of G and C bases in the window), as well as the average read depth which was obtained by aligning Illumina paired-end reads to the reference genome. This resulted in the data points of GC contents and read depth in Supplementary Figure 4.

### 1.4 Genome assembly

The KP genome was assembled by SOAPdenovo2 [2] using the default parameters. Firstly, the contigs were constructed using the filtered reads of paired-end libraries (170bp, 800bp); the mate-paired reads were used to bridge the contigs; finally the assembly gaps were filled using the reads of paired-end libraries by GapCloser1.10. The final KP assembly was around 1.24 Gb.

**Supplementary Table 4** Statistics of the assembly quality of the KP genome as represented by length for number of contigs and scaffolds.

| | contig | | scaffold | |
|---|---|---|---|---|
| | **Length (bp)** | **number** | **Length (bp)** | **number** |
| Total | - | 1,207,355,267 | - | 1,245,524,081 |
| Max | - | 290,908 | - | 15,291,072 |
| number>=100 | - | 194,903 | - | 126,040 |
| number>=2000 | - | 50,293 | - | 3,299 |
| N50 | 38,982 | 9,113 | 3,220,723 | 107 |
| N60 | 30,946 | 12,582 | 2,333,970 | 153 |
| N70 | 23,532 | 17,058 | 1,539,735 | 217 |
| N80 | 16,467 | 23,162 | 881,669 | 322 |
| N90 | 8,903 | 32,942 | 360,831 | 540 |

#### 1.5 Evaluating the assembly quality and completeness

To compare the KP genome assembly with two other bird species whose genome have been relatively thoroughly assembled, i.e., chicken (*Gallus gallus*) and zebra finch (*Taeniopygia guttata*), we aligned their whole genomes pairwise using LASTZ [3] with parameters as "T=2 C=2 H=2000 Y=3400 L=6000 K=2200". The genome base content was calculated by running an in-house Perl script.

**Supplementary Table 5** Comparisons of genome size and sequencing quality between KP and two published bird genomes, the domestic chicken (*Gallus gallus*) and the zebra finch (*Taeniopygia guttata*).

| Genome | # scaffolds | Genome size (bp, with N) | Genome Size (bp, without N) | Masked Size (bp) | %masked |
|---|---|---|---|---|---|
| Chicken *Gallus gallus* | 55 | 1,108,466,630 | 948,566,554 | 101,764,878 | 9.15 |
| Zebra finch *Taeniopygia guttata* | 68 | 1,235,794,146 | 1,137,254,349 | 88,175,047 | 7.14 |
| Kentish Plover *Charadrius alexandrinus* | 126040 | 1,245,524,081 | 1,122,176,016 | 85,218,351 | 6.84 |

**Supplementary Table 6** Statistics of alignment between the three avian genomes (chicken, zebra finch and KP)

| Species vs Species | Query Aligned (bp) | Query Coverage (%) | Target Aligned (bp) | Target Coverage (%) |
|---|---|---|---|---|
| *G. gallus* vs *T. guttata* | 482,904,967 | 39.08 | 481,584,252 | 43.45 |
| *G. gallus* vs *C. alexandrinus* | 836,915,653 | 67.19 | 811,084,521 | 73.17 |
| *T. guttata* vs *C. alexandrinus* | 922,005,778 | 74.03 | 893,365,194 | 72.29 |

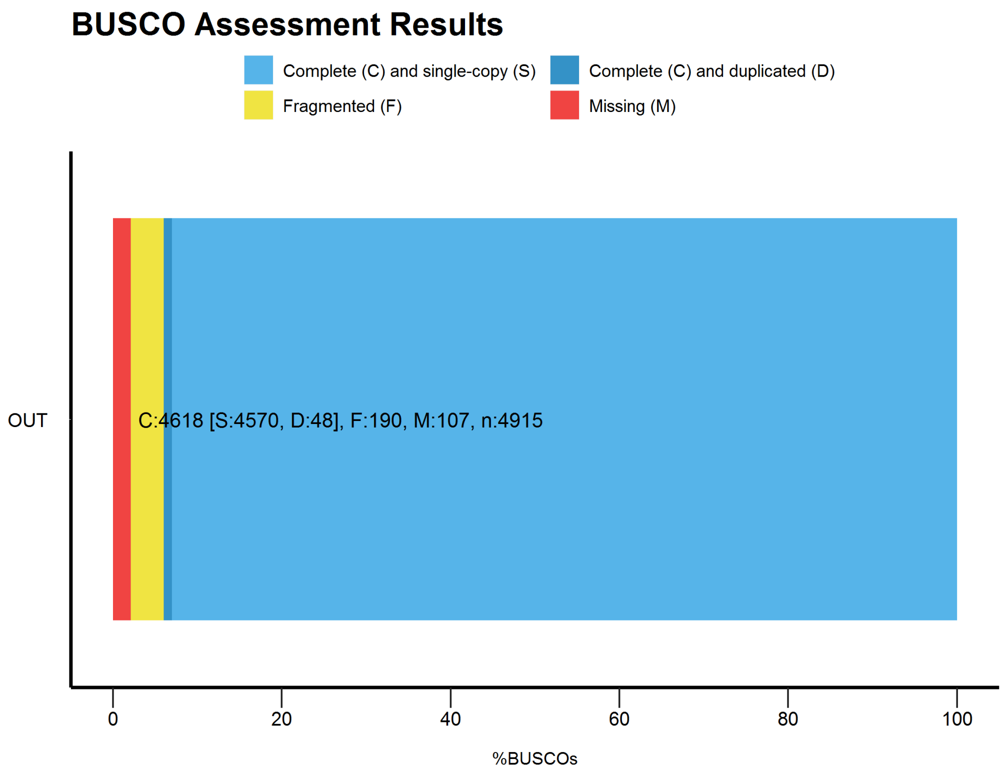


**Supplementary Figure 4** BUSCO Assessment results of KP assembly

To evaluate the KP genome assembly completeness, we used BLAST v2.6.0, HMMER v3.1, AUGUSTUS v3.2.3 and BUSCO v2.0.1 [4] with parameters: "python BUSCO.py -l aves_odb9 -m genome -c -sp chicken". The results were: 'C:94.0%[S:93.0%,D:1.0%],F:3.9%,M:2.1%,n:4915'.

**Supplementary Table 7** Statistics of genome content of the *de novo* KP genome, calculated by running an in-house Perl script.

| Sample ID | Number (bp) | % of genome |
|---|---|---|
| A | 341,442,006 | 27.85 |
| T | 340,760,358 | 27.80 |
| C | 252,389,362 | 20.59 |
| G | 253,091,044 | 20.65 |
| N | 38,135,881 | 3.11 |
| GC | 505,480,406 | 42.56* |
| Total(bp) | 1,225,818,651 | |

* GC content of the genome without N

### 2. RNA sequencing

The RNA-seq libraries were sequenced 90 bp at each end using Illumina Hiseq 2000 platform. The RNA-seq reads were mapped to the KP genome using Tophat with default parameters, and subsequently analyzed using in-house Perl scripts. The RNA-seq results were validated using qRT-PCR, with five biological replicates for each stage. All data were expressed as mean ± standard error of mean and were evaluated by one-way ANOVA followed by Tukey's honestly significant difference test for adjusting *P* values from multiple comparisons. Results were considered to be statistically significant for *P* values <0.05.

### 3. Genome analysis

#### 3.1 Transposable element analysis of KP genome

We constructed a transposable element (TE) library of KP genome using a combination of homology-based and *de novo* approaches.

1) Tandem repeats were identified using Tandem Repeats Finder (v4.05, http://tandem.bu.edu/trf/trf.html) [5].
2) RepeatMasker (v3.3.0, http://www.repeatmasker.org/) and RepeatProteinMask were employed to identify TE based on homologous search against a library of Repbase [6] (Release 16.03) using the parameters "-nolow -no_is -norna -parallel 1" and "-noLowSimple -pvalue 1e-4".
3) *Ab initio* TE library was constructed using RepeatModeler (v1.08, http://www.repeatmasker.org/RepeatModeler.html) with default parameters. RepeatModeler identifies repeat elements by integrating two repeat finding programs RECON [7] and RepeatScout [8]. Using the repeat library constructed with RepeatModeler, we estimated the repeat content of KP genome using RepeatMasker v4.0.5 with the sensitive mode (-s). The TE expansion history was constructed by first recalculating the divergence of the identified TE copies in the genome with the corresponding consensus sequence in the TE library using Kimura distance [9] and then the percentage of TE in the genome was estimated at difference divergence levels.

**Supplementary Table 8** Summary of the proportion of repeats in the KP genome as estimated by different methods, i.e. Tandem Repeats Finder, RepeatMasker and Repeat Protein Mask.

| **Type** | **Repeat Size (bp)** | **% of genome** |
|---|---|---|
| Tandem Repeats Finder | 15,064,689 | 1.209506 |
| Repeat Masker | 85,263,191 | 6.845567 |
| Repeat Protein Mask | 53,067,439 | 4.260651 |
| De novo | 95,269,107 | 7.648917 |
| Total | 132,695,100 | 10.653756 |

**Supplementary Table 9** Summary of the KP genome's transposable element content and their characteristics within the KP genome

| | **RepBase TEs** | | **TE Proteins** | | **De novo** | | **Combined TEs** | |
|---|---|---|---|---|---|---|---|---|
| | **Length (Kbp)** | **%in Genome** | **Length (Kbp)** | **% in Genome** | **Length (Kbp)** | **% in Genome** | **Length (Kbp)** | **% in Genome** |
| DNA | 7,002 | 0.56 | 239 | 0.02 | 655 | 0.05 | 7,703 | 0.62 |
| LINE | 6,897 | 5.54 | 5,143 | 4.13 | 7,824 | 6.28 | 101,594 | 8.16 |
| SINE | 1,300 | 0.10 | 0 | 0.00 | 74 | 0.01 | 1,360 | 0.11 |
| LTR | 9,156 | 0.74 | 1,410 | 0.11 | 2,051 | 1.65 | 27,712 | 2.22 |
| Other | 3 | 0.00 | 0 | 0.00 | 0 | 0.00 | 3 | 0.00 |
| Unknown | 0 | 0.00 | 0 | 0.00 | 596 | 0.05 | 596 | 0.05 |
| Total | 85,263 | 6.85 | 5,307 | 4.26 | 9,441 | 7.58 | 121,000 | 9.71 |

For the non-coding RNA annotation, we used the software tRNAscan-SE [10] to find the tRNA sequence in Kentish plover genome according to the tRNA architectural feature. Since rRNA sequences are highly conservative, we used similar species rRNA sequences as a reference to find rRNA sequence in the genome using BLASTN. We also used the software INFERNAL [11] with a Rfam family's covariance model to predict the miRNA and snRNA sequences in the genome.

**Supplementary Table 10** Summary of predicted RNA genes and their characteristics within the KP genome

| Type | | Copy | Average length (bp) | Total length (bp) | % of genome |
|---|---|---|---|---|---|
| miRNA | | 195 | 85.1538 | 16,605 | 0.001333 |
| tRNA | | 229 | 74.9039 | 17,153 | 0.001377 |
| rRNA | | 62 | 138.0322 | 8,558 | 0.000687 |
| rRNA | 18S | 13 | 147.6153 | 1,919 | 0.000154 |
| | 28S | 40 | 146.8250 | 5,873 | 0.000472 |
| | 5.8S | 2 | 155.0000 | 310 | 0.000025 |
| | 5S | 7 | 65.1429 | 456 | 0.000037 |
| snRNA | | 236 | 123.3475 | 29,110 | 0.002337 |
| snRNA | CD-box | 108 | 95.1574 | 10,277 | 0.000825 |
| | HACA-box | 73 | 141.5068 | 10,330 | 0.000829 |
| | splicing | 42 | 153.3095 | 6,439 | 0.000517 |

#### 3.2 Gene prediction and annotation

The RNA-seq reads were assembled into transcripts using the following steps.

1) The raw reads were aligned to the genome using TopHat [12] using default parameters and assembled into transcripts using Cufflinks [13].
2) The assembled transcripts were used to refine the gene models of homology-based approaches. Overlapping gene models of RNA-seq and homology-based approaches were merged.
3) We also predicted gene models using the transcripts that didn't overlap with the homology-based models. A fifth-order Markov model was trained using 1,000 intact gene models of the homology-based approach and was used to predict ORF of RNA transcripts.

Finally, 15,677 gene models were produced.

**Supplementary Table 11** Summary of predicted protein coding genes and their characteristics in the KP genome as compared with the chicken, turkey and zebra finch genomes

| Gene set | | Number | Average gene length (bp) | Average CDS length (bp) | Average exon per gene | Average exon length (bp) | Average intron length (bp) |
|---|---|---|---|---|---|---|---|
| Transcripts | TopHat+ cufflinks | 8,893 | 19,394.48 | 1,281.39 | 7.995 | 160.268 | 2,103.57 |
| Homolog | *Gallus gallus* | 19,284 | 16,281.10 | 1,364.75 | 7.292 | 187.148 | 2,370.57 |
| | *Meleagris gallopavo* | 33,186 | 10,110.78 | 9,24.346 | 4.747 | 194.736 | 2,451.89 |
| | *Taeniopygia guttata* | 15,921 | 17,760.18 | 1,354.04 | 7.668 | 176.588 | 2,460.50 |
| Integrate | | 26,097 | 15,359.36 | 1,169.26 | 6.174 | 189.388 | 2,621.98 |
| Final set | | 15,677 | 22,632.24 | 1,601.90 | 9.01 | 177.783 | 2,495.63 |

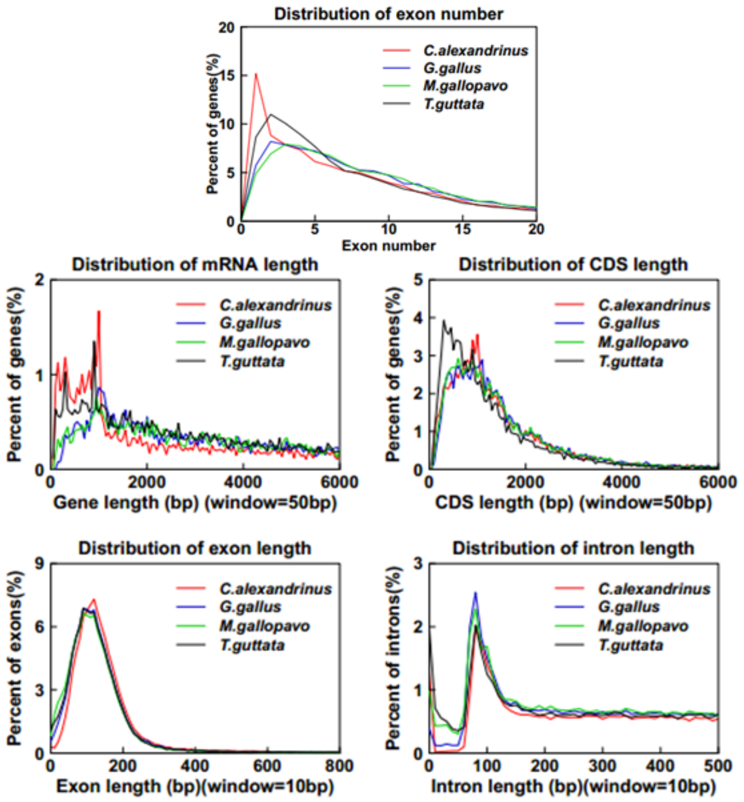


**Supplementary Figure 3** Comparison of gene parameters, i.e. distribution of 1) exon number, 2) mRNA length, 3) CDS length, 4) exon length, 5) intron length, among four avian genomes (KP, the domestic chicken *Gallus gallus*, turkey *Meleagris gallopavo* and zebra finch *Taeniopygia guttata* )

We annotated the predicted gene models using InterPro [14] KEGG [15], Swiss-Prot [16] and TrEMBL [16] databases using the following steps.

1) The gene symbol and pathway were assigned based on the best blast hit against Swiss-Prot and KEGG databases.
2) The motifs and domains in protein sequences were annotated using InterProScan [17] by searching publicly available databases, including Pfam, PRINTS, PANTHER, PROSITE, ProDom, and SMART.
3) Gene Ontology [18] terms were assigned using Blast2GO [19, 20].

**Supplementary Table 12** Number of KP genes assigned to functional categories using different databases

| | | Number | Percent (%) |
|---|---|---|---|
| Total | | 15,677 | |
| Annotated | InterPro | 14,502 | 92.504944 |
| | GO | 11,035 | 70.389743 |
| | KEGG | 13,685 | 87.293487 |
| | Swissprot | 15,265 | 97.371946 |
| | TrEMBL | 15,639 | 99.757607 |
| | Annotated | 15,644 | 99.789501 |
| Unannotated | | 33 | 0.210499 |

### 4. Gene family evolution

#### 4.1 Gene family analysis

The gene family analysis was conducted by Treefam using the following steps.

1) Protein sequences of KP and other 15 waterbirds (Adélie penguin, *Pygoscelis adeliae*; Common cormorant, *Phalacrocorax carbo*; Crested ibis, *Nipponia nippon*; Dalmatian pelican, *Pelecanus crispus*; Emperor penguin, *Aptenodytes forsteri*; Great crested grebe, *Podiceps cristatus*; Greater flamingo, *Phoenicopterus ruber*; Grey-crowned crane, *Balearica regulorum*; Killdeer, *Charadrius vociferus*; Little egret, *Egretta garzetta*; Mallard, *Anas platyrhynchos*; Northern fulmar, *Fulmarus glacialis*; Red-throated loon, *Gavia stellata*; Sunbittern, *Eurypyga helias* and White-tailed tropicbird, *Phaethon lepturus*) were downloaded from the GigaScience database [21]. BLASTP [22] was employed to identify potential homologous genes using E-value<1e-10.
2) The raw Blast results were refined using solar (an in-house software, version0.9.6) by which the high-scoring segment pairs (HSPs) were conjoined.
3) Similarity between protein sequences were evaluated using bit-score, followed by clustering protein sequences into gene families using hcluster_sg, a hierarchical clustering algorithm in the Treefam pipeline (version 0.50) with the parameters "-w 5 -s 0.33 -m 100000".

**Supplementary Table 10** Summary of unigene and unclustered gene of predicted RNA genes and their characteristics among 16 waterbird species

| Species | Genes number | Unclustered genes | Family number | Unique families | Average genes per family |
|---|---|---|---|---|---|
| Adélie penguin (*Pygoscelis adeliae*) | 15,270 | 33 | 13,508 | 1 | 1.13 |
| Common cormorant (*Phalacrocorax carbo*) | 13,479 | 69 | 12,210 | 5 | 1.1 |
| Crested ibis (*Nipponia nippon*) | 16,756 | 96 | 14,164 | 3 | 1.18 |
| Dalmatian pelican (*Pelecanus crispus*) | 14,813 | 79 | 12,668 | 3 | 1.16 |
| Emperor penguin (*Aptenodytes forsteri*) | 16,070 | 42 | 14,104 | 0 | 1.14 |
| Great crested Grebe (*Podiceps cristatus*) | 13,913 | 79 | 12,040 | 12 | 1.15 |
| Greater Flamingo (*Phoenicopterus ruber*) | 14,024 | 61 | 12,385 | 3 | 1.13 |
| Grey-crowned crane (*Balearica regulorum*) | 14,173 | 46 | 12,838 | 5 | 1.1 |
| Kentish Plover (*Charadrius alexandrinus*) | 15,677 | 514 | 12,435 | 43 | 1.22 |
| Killdeer (*Charadrius vociferus*) | 16,860 | 59 | 13,874 | 29 | 1.21 |
| Little egret (*Egretta garzetta*) | 16,585 | 27 | 12,843 | 32 | 1.29 |
| Mallard (*Anas platyrhynchos*) | 16,521 | 306 | 14,452 | 65 | 1.12 |
| Northern fulmar (*Fulmarus glacialis*) | 14,306 | 48 | 12,714 | 4 | 1.12 |
| Red-throated loon (*Gavia stellata*) | 13,454 | 63 | 12,278 | 3 | 1.09 |
| Sunbittern (*Eurypyga helias*) | 13,974 | 107 | 12,117 | 19 | 1.14 |
| White-tailed Tropicbird (*Phaethon lepturus*) | 14,970 | 65 | 12,851 | 13 | 1.16 |

The gene family cluster result of 16 species is shown in Supplementary Figure 6 and 7.

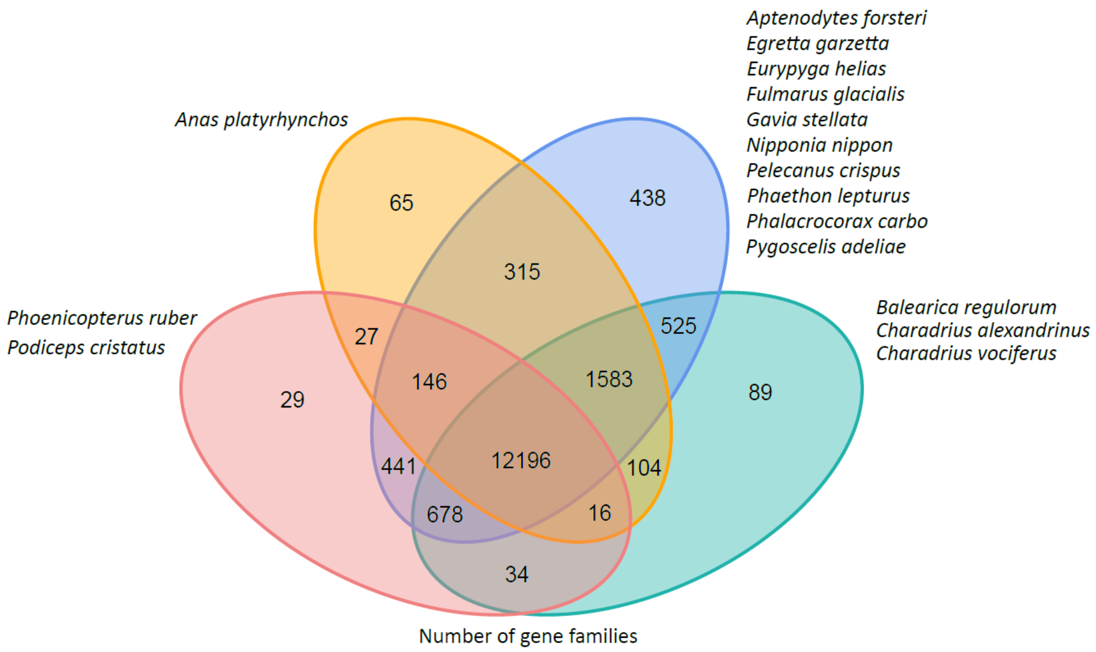


**Supplementary Figure 6** Orthologous gene clusters shared between the KP genome and the genomes of 16 species of waterbirds

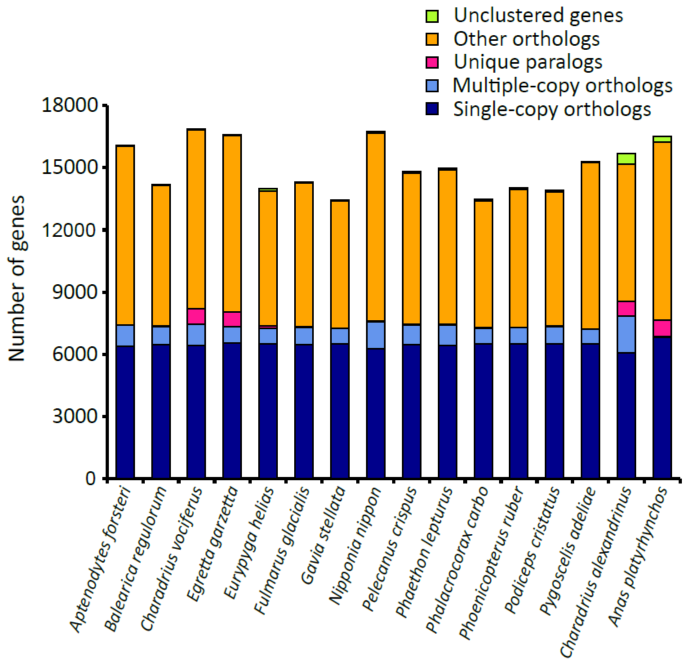


**Supplementary Figure 7** Characteristics of gene family evolution in KP and other 15 species of waterbirds

Other orthologs: orthologous genes that could only be found in some species but not in all 16 species.
Unique paralogs: lineage specific genes.
Multiple-copy orthologs: species contains multiple copies of orthologous gene.
Single-copy orthologs: species contains one copy of orthologous gene.

**4.2 Phylogenetic tree construction and divergence time estimate**

12,196 one-to-one orthologous genes were obtained from the gene family analysis using the pipeline described previously. The protein sequences of one-to-one orthologous genes were aligned using MUSCLE with the default parameters [23]. We then filtered the saturated sites and poorly aligned regions using trimAl [24] with the parameters “-gt 0.8 -st 0.001 -cons 60”. After trimming the saturated sites and poorly aligned regions in the concatenated alignment, amino acids were used in the phylogenomics reconstruction. The trimmed protein alignments were used as a guide to align corresponding coding sequences (CDS). The aligned protein and the four-fold degenerated sites in the CDS sequences were each concatenated into a super gene using an in-house Perl script.

The phylogenomic tree was reconstructed using RaxML version 8.1.19 [25] based on the concatenated protein sequences. Specifically, we used PROTGAMMAAUTO parameter to select the optimal amino acid substitution model, specified the spotted gar as the outgroup, and evaluated the robustness of the result using 100 bootstraps. To compare the neutral mutation rate of different species, we also generated a phylogeny based on the four-fold degenerate sites. The phylogenomics topology was used as the input to optimize the branch lengths of alignment of four-fold degenerate sites using the "-f e" parameter in Raxml under the general time reversible (GTR) model, suggested by modelgenerator version 0.8537. We calculated the pairwise distances to the outgroup [26]. We calculated the pairwise distances to the outgroup (duck) based on the optimized branch length of neutral tree using the ‘cophenetic.phylo’ module in the R-package ‘ape’ [27].

We used the CDS sequences alignment and its tree from the phylogenetic tree analysis as the input, and used MCMCTree implemented in the PAML version 4 [28] to estimate the divergence time between KP and other waterbirds.

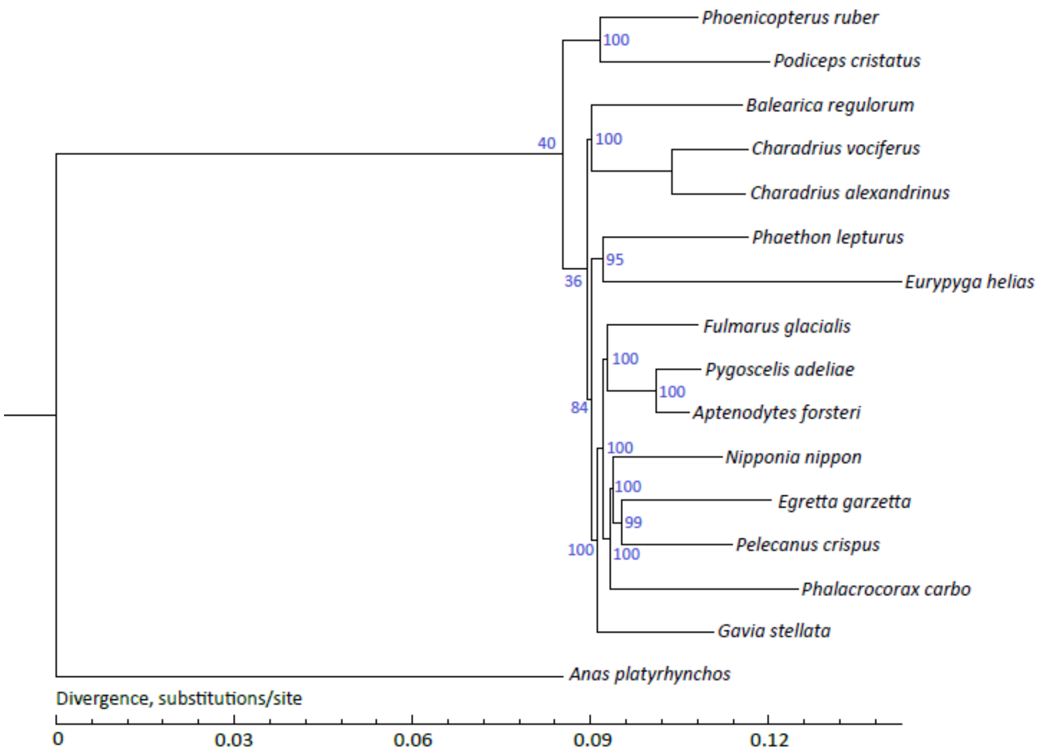


**Supplementary Figure 4** Evolutionary relationships between KP and 15 other published waterbird species using coding DNA sequence. The phylogeny was reconstructed using the alignment of four-fold degenerate sites. The robustness of the result was calculated using 100 bootstraps and is represented in blue for each node.

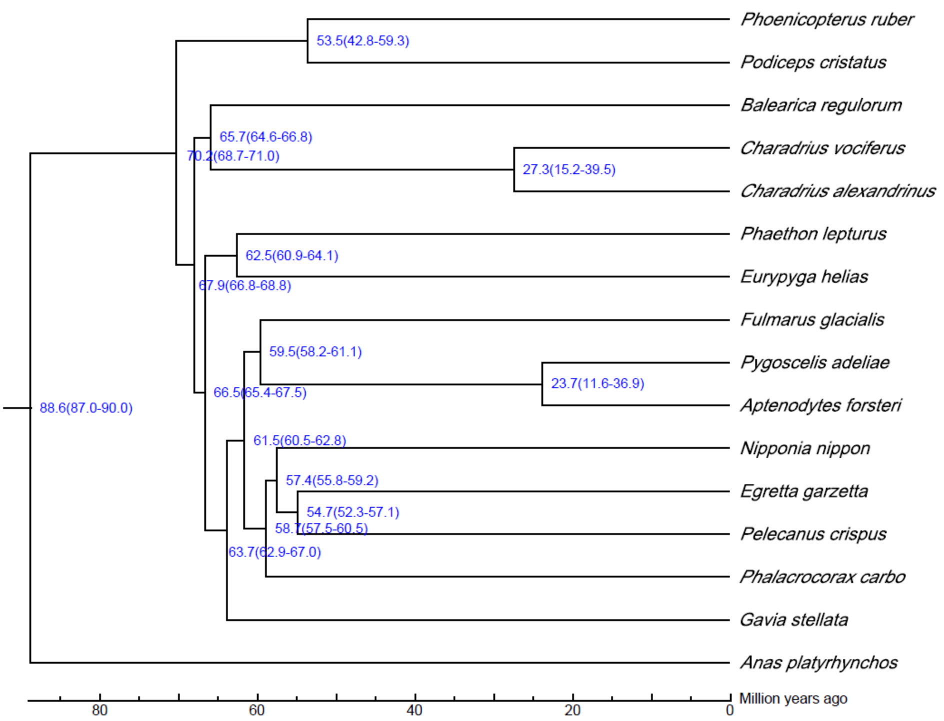


**Supplementary Figure 5** Divergence times between the KP and 15 waterbird species. The numbers on each node represent the divergence time in millions of years, and the corresponding 95% credible intervals of the divergence times are indicated in brackets.

### 5. Whole genome resequencing for KP and WFP

#### 5.1 Samples quality checking

We detected the DNA samples from 22 individuals in three methods.

1) Agarose gel electrophoresis: to analysis the degree of DNA degradation and whether polluted by RNA.
2) Nanodrop: to measure the purity of DNA (OD260/OD280).
3) Qubit: to measure the concentration of DNA.

The DNA samples with OD260/OD280 ratios 1.8 to 2.0 and concentrations higher than 1.5ug were chosen to build DNA libraries.

#### 5.2 High-throughput sequencing and quality control

The DNA libraries were preliminarily quantified using Qubit2.0 and diluted to 1ng/ul. The insert size was detected using Agilent 2100 to obtain the expectant libraries. The effective concentration of the database was accurately quantified (>2nM) using Q-PCR to ensure the libraries' quality. After the quality control, the qualified libraries were sequencing by Illumina Hiseq platform.

We carried the following bioinformatics analyses on every resequencing samples after obtaining the raw data.

Firstly, we filtered out the raw data according to the following criteria:

1) The reads containing adapter sequences.
2) The reads containing higher than 50% low quality bases (quality score<5).
3) The reads containing higher than 10% N bases.

**Supplementary Table 11** Statistics of 21 re-sequenced genomes of KP (samples in bold) and WFP (samples in italic)

| Sample | Clean Reads Number | Clean Bases Number | Low-quality Reads Rate(%) | Ns Reads Rate(%) | Clean Q30 Bases Rate(%) | Clean GC(%) |
|---|---|---|---|---|---|---|
| **QH_1** | 46,551,514 | 6.98 Gb | 4.29 | 0 | 85.82 | 44.88 |
| **QH_2** | 47,461,352 | 7.12 Gb | 4.15 | 0 | 86.00 | 45.95 |
| **TS_1** | 43,934,860 | 6.59 Gb | 0 | 0 | 88.19 | 44.07 |
| **TS_2** | 41,963,880 | 6.29 Gb | 0 | 0 | 88.35 | 44.39 |
| **LYG_1** | 44,028,162 | 6.6 Gb | 4.36 | 0 | 86.10 | 43.02 |
| **LYG_2** | 43,536,758 | 6.53 Gb | 0 | 0 | 88.63 | 41.79 |
| **RD_1** | 48,336,134 | 7.25 Gb | 4.03 | 0 | 86.51 | 42.07 |
| **RD_2** | 40,801,248 | 6.12 Gb | 0 | 0 | 88.42 | 41.82 |
| **ZS_1** | 46,851,774 | 7.0 Gb | 5.51 | 0 | 86.40 | 45.09 |
| **ZS_2** | 47,753,350 | 7.13 Gb | 4.48 | 0 | 87.12 | 45.47 |
| *FZ_1* | 51,757,222 | 7.73 Gb | 4.81 | 0 | 86.69 | 44.45 |
| *FZ_2* | 48,968,132 | 7.32 Gb | 4.67 | 0 | 86.91 | 46.28 |
| *XM_1* | 47,992,388 | 7.17 Gb | 6.47 | 0 | 85.87 | 44.88 |
| *SW_1* | 51,379,250 | 7.68 Gb | 6.07 | 0 | 86.11 | 43.53 |
| *SW_2* | 46,003,762 | 6.87 Gb | 4.03 | 0 | 87.40 | 42.43 |
| *ZJ_1* | 42,983,608 | 6.45 Gb | 0 | 0 | 88.71 | 42.35 |
| *ZJ_2* | 39,715,194 | 5.96 Gb | 0 | 0 | 88.35 | 42.38 |
| *BH_1* | 45,453,048 | 6.82 Gb | 4.68 | 0 | 85.84 | 43.04 |
| *BH_2* | 46,553,268 | 6.98 Gb | 4.24 | 0 | 86.03 | 43.08 |
| *HN_1* | 42,504,486 | 6.38 Gb | 0 | 0 | 88.28 | 43.95 |
| *HN_2* | 251,843,604 | 37.77Gb | 0.44 | 0 | 93.43 | 45.01 |

**5.3 Read mapping and SNPs calling**

We used BWA to align the high-quality data to the reference genome with the parameters “-R -a -M -C” and outputted the alignment in BAM format. Then we used Picard MarkDuplicates to identify and filter out the PCR duplication with parameters “VALIDATION_STRINGENCY=
LENIENT REMOVE_DUPLICATES=false”. The filtered alignments were then used to call the SNPs and InDels for all of the samples jointly using GATK HaplotypeCaller (v3.3-0) [29] with parameters “-ERC GVCF -variant_index_type LINEAR -variant_index_parameter 128000 --dbsnp -L”.

The SNPs were filtered using inhouse Perl script by following criteria:

1) Missing rate <= 0.10
2) Allele frequency > 0.05
3) Each 10 bp <=3 SNPs

After the SNP calling and filtering, we obtained a total of 11,959,725 high quality SNPs to carry the downstream analyses.

**Supplementary Table 12** The average coverage of 20 re-sequenced genomes (5x) of KP and WFP

| Sample ID | Kentish plover | White-faced plover |
|---|---|---|
| Average mapping depth | 5.42 | 5.57 |
| Mapping rate (%) | 99.11 | 98.99 |
| Coverage (%) | 97.08 | 97.38 |
| Coverage at least 4× (%) | 69.93 | 72.17 |
| Coverage at least 10× (%) | 8.86 | 9.39 |

### 6. Inference of demographic history of the two species

#### 6.1 Settings of ABC models

**Supplementary Table 13** Priors of the models for the divergence pattern between the KP and WFP

| **model** | **T1/10^6** | ***Ne_K*/10^6** | ***Ne_W*/10^6** | ***Ne_A*/10^6** | ***N1_K*/10^6** | ***N1_W*/10^6** | ***G_K*** | ***G_W*** | ***M*** |
|---|---|---|---|---|---|---|---|---|---|
| **A1** | - | 0.01-20 | 0.01-20 | 0.01-20 | - | - | - | - | - |
| **A2** | - | 0.01-20 | 0.01-20 | 0.01-20 | - | - | - | - | 0-500 |
| **A3** | 0-1 | 0.01-20 | 0.01-20 | 0.01-20 | - | - | - | - | 0-500 |
| **A4** | 0-1 | 0.01-20 | 0.01-20 | 0.01-20 | - | - | - | - | 0-500 |
| **B1** | 0.008-0.015 | 0.05-10 | 0.01-1 | 0.05-10 | 0.05-5 | 0.05-5 | 1-14 | 2-25 | - |
| **B2** | 0.008-0.015 | 0.05-10 | 0.01-1 | 0.05-11 | 0.05-5 | 0.05-5 | 1-14 | 2-25 | 0-500 |
| **B3** | 0.008-0.015 | 0.05-10 | 0.01-1 | 0.05-12 | 0.05-5 | 0.05-5 | 1-14 | 2-25 | 0-500 |
| **B4** | 0.008-0.015 | 0.05-10 | 0.01-1 | 0.05-13 | 0.05-5 | 0.05-5 | 1-14 | 2-25 | 0-500 |

K: Kentish plover; W: white-faced plover; A: ancestral population; *Ne*: effective population size; T1: migration/Ne change time; M: migration rate. Population split time range was 0-1 million year and N0 was $10^6$ in every model. Model illustrations are in Supplementary Figure 8.

### 6.2 ABC model choice and posterior density

**Supplementary Table 14** Models of divergence patterns and their corresponding posterior probabilities for model selection in ABC analyses.

| **Group** | **Models** | **%** | **Bayes Factors** | | | |
|---|---|---|---|---|---|---|
| | | | 1 | 2 | 3 | 4 |
| A. Constant Ne | 1. Isolation | 0.00 | 1.00 | 0.00 | 0.00 | 0.00 |
| | 2. Isolation with migration | 0.00 | 3.26E+09 | 1.00 | 0.00 | 0.00 |
| | **3. Early gene flow** | **1.00** | 1.44E+22 | 4.43E+12 | 1.00 | 1.97E+07 |
| | 4. Secondary contact | 0.00 | 7.33E+14 | 2.25E+05 | 0.00 | 1.00 |
| B. Changing Ne | 1. Isolation | 0.00 | 1.00 | 0.00 | 0.00 | 0.00 |
| | 2. Isolation with migration | 0.00 | 7.97E+33 | 1.00 | 0.00 | 0.00 |
| | 3. Early gene flow | 0.02 | 6.81E+40 | 8.54E+06 | 1.00 | 0.02 |
| | **4. Secondary contact** | **0.98** | 3.26E+42 | 4.09E+08 | 47.89 | 1.00 |
| A & B | A3 | 0.00 | 1.00 | 4.03E+04 | 3.00E-04 | 0.00 |
| | A4 | 0.00 | 0.00 | 1.00 | 0.00 | 0.00 |
| | B3 | 0.08 | 3.71E+03 | 1.50E+08 | 1.00 | 0.08 |
| | **B4** | **0.92** | 4.49E+04 | 1.81E+09 | 12.09 | 1.00 |

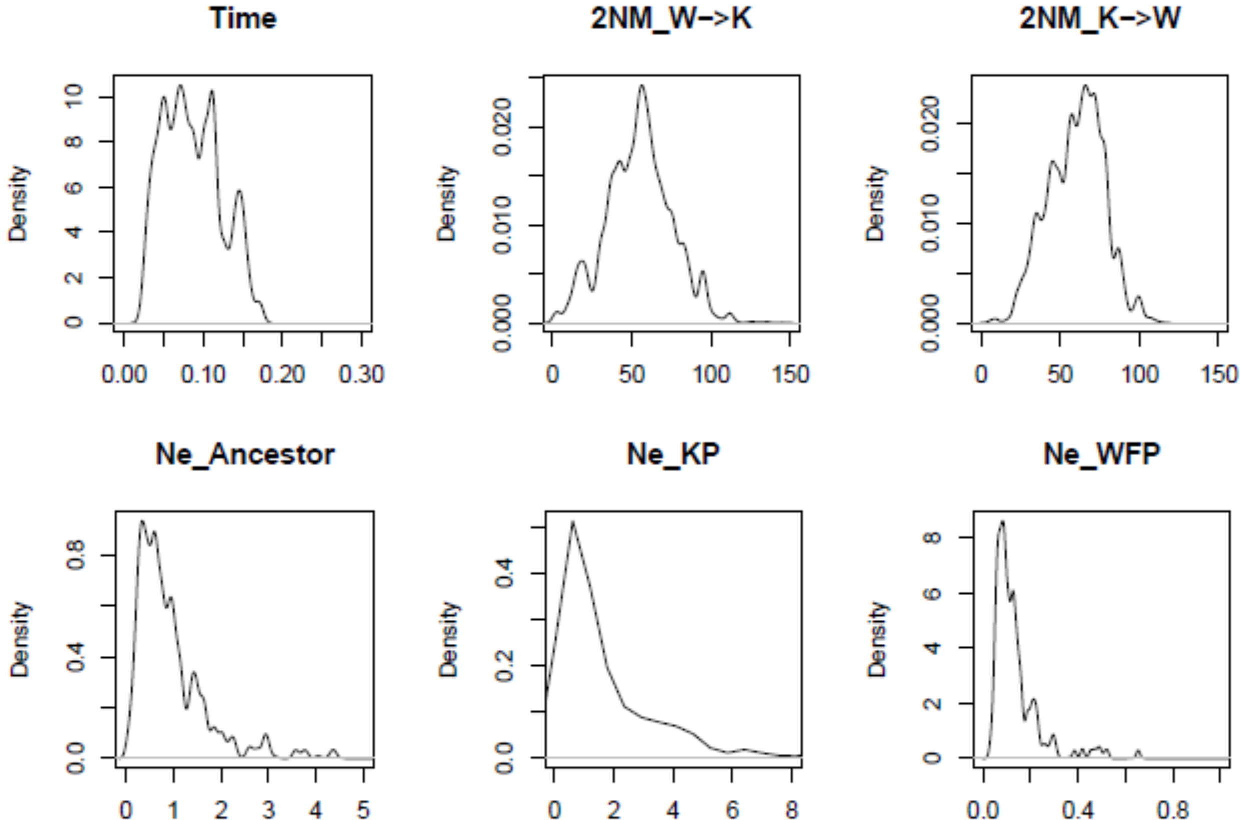


**Supplementary Figure 6** Posterior densities of population parameters of model B4, secondary contact model with changing effective population sizes based on PSMC. Divergent time was measured in $10^7$ years and Ne in $10^6$. K or KP represents the KP *Charadrius alexandrinus* and W or WFP represents the white-faced Plover *C. dealbatus*. **Upper left:** Population divergence times (*T*) of *C. alexandrinus* and *C. dealbatus*. Population migration rates (*2NM*) represent the average number of individuals in each group that had previously migrated from the other group. **Upper middle:** gene flow from *C. dealbatus* to *C. alexandrinus* and vice versa for **Upper right**. **Bottom:** Effective population sizes (*Ne*) of the two species (Ne_KP & Ne_WFP) and their most recent common ancestor (Ne_Ancestor).

### 7. Detection of genomic region under selection

Gene symbols were assigned based on best reciprocal BLAST matches. The BLAST XML file containing outputs from the Kentish plover proteins BLASTed against the RefSeq protein database was split into smaller files using Biostar165777 (http://dx.doi.org/10.6084/m9.figshare.1425030). The resulting files were then parsed using blastxmlparser (http://dx.doi.org/10.1093/bioinformatics/btq475). Parsed files were combined and top hits were extracted first by lowest E-value and then by highest bit score. Where multiple hits had equal E-values and bit scores, multiple best hits were used. Protein sequences for top hits were obtained using Entrez Direct (https://www.ncbi.nlm.nih.gov/books/NBK179288). Sequences were BLASTed against a database of Kentish plover proteins and top hits extracted. Gene symbols were obtained for reciprocal best hits using Entrez Direct.

**Supplementary Table 18** High diversity genes and gene descriptions. Genes were considered to have high $F_{ST}$ if they fell above the 95$^{th}$ percentile of $F_{ST}$ values (autosomes: $F_{ST} \geq 0.184$, Z: $F_{ST} \geq 0.262$). Gene symbols were assigned based on best reciprocal BLAST matches from the RefSeq protein database.

| Gene Name | Gene Symbol | *FST* | Chromosome |
|---|---|---|---|
| CHAAL00000012431 | CHRNB3 | 0.773974814 | Z |
| CHAAL00000011746 | PIGG | 0.768934531 | Z |
| CHAAL00000012272 | CLTA | 0.768934531 | Z |
| CHAAL00000012462 | CITED4 | 0.736842105 | Autosome |
| CHAAL00000002440 | | 0.706477733 | Autosome |
| CHAAL00000013705 | SUCLG2 | 0.682261209 | Autosome |
| CHAAL00000010671 | EFEMP1 | 0.658802178 | Autosome |
| CHAAL00000005765 | PPP4R4 | 0.578947368 | Autosome |
| CHAAL00000012451 | FKTN | 0.57375834 | Z |
| CHAAL00000000807 | LRRC3B | 0.564327485 | Autosome |
| CHAAL00000011583 | NR1H3 | 0.525416104 | Autosome |
| CHAAL00000001248 | | 0.5215311 | Autosome |
| CHAAL00000003458 | MPEG1 | 0.5215311 | Autosome |
| CHAAL00000003501 | CDC37L1 | 0.516900756 | Z |
| CHAAL00000004224 | DCAF1 | 0.495906433 | Autosome |
| CHAAL00000013419 | LACTBL1 | 0.491525424 | Autosome |
| CHAAL00000003675 | SMIM8 | 0.48706512 | Autosome |
| CHAAL00000010452 | TMEM38A | 0.480290685 | Autosome |
| CHAAL00000014940 | | 0.477339181 | Autosome |
| CHAAL00000015124 | LOC104387136 | 0.474769398 | Autosome |

| | | | |
|---|---|---|---|
| CHAAL00000000208 | HACD3 | 0.473684211 | Autosome |
| CHAAL00000013108 | RPL23 | 0.473684211 | Autosome |
| CHAAL00000008264 | PTBP3 | 0.473684211 | Z |
| CHAAL00000004132 | OLFML3 | 0.460980036 | Autosome |
| CHAAL00000014470 | DTWD1 | 0.460950764 | Autosome |
| CHAAL00000000774 | IL13RA1 | 0.460274891 | Autosome |
| CHAAL00000009899 | RNF144A | 0.435406699 | Autosome |
| CHAAL00000008463 | | 0.43192853 | Z |
| CHAAL00000009613 | RABGAP1L | 0.43062201 | Autosome |
| CHAAL00000004714 | IQCH | 0.430226944 | Autosome |
| CHAAL00000008274 | PLPPR1 | 0.430226944 | Z |
| CHAAL00000008625 | LOC104291089 | 0.430034416 | Autosome |
| CHAAL00000014048 | LOC105412744 | 0.428409734 | Autosome |
| CHAAL00000006239 | JAK2 | 0.428409734 | Z |
| CHAAL00000011709 | FKBP6 | 0.423323788 | Autosome |
| CHAAL00000005176 | HDAC1 | 0.421239933 | Autosome |
| CHAAL00000001970 | PLIN2 | 0.421052632 | Autosome |
| CHAAL00000005544 | CPTP | 0.421052632 | Autosome |
| CHAAL00000007430 | | 0.421052632 | Autosome |
| CHAAL00000008811 | MGAT4B | 0.421052632 | Autosome |
| CHAAL00000009839 | PRICKLE4 | 0.421052632 | Autosome |
| CHAAL00000011446 | DHCR7 | 0.421052632 | Autosome |
| CHAAL00000014341 | LOC103920693 | 0.421052632 | Autosome |
| CHAAL00000014916 | RHOBTB3 | 0.421052632 | Z |
| CHAAL00000003441 | | 0.418350277 | Z |
| CHAAL00000005626 | UAP1 | 0.418282548 | Autosome |
| CHAAL00000011770 | LRP11 | 0.415563448 | Autosome |
| CHAAL00000007052 | | 0.409338393 | Z |
| CHAAL00000000117 | CARS | 0.401088929 | Autosome |
| CHAAL00000000544 | SFN | 0.401088929 | Autosome |
| CHAAL00000013102 | CHCHD6 | 0.401088929 | Autosome |
| CHAAL00000000116 | OSBPL5 | 0.398877905 | Autosome |
| CHAAL00000015549 | TACR1 | 0.394736842 | Autosome |
| CHAAL00000008256 | SMC2 | 0.394522893 | Z |
| CHAAL00000011178 | FEM1B | 0.391660971 | Autosome |
| CHAAL00000013794 | | 0.389348743 | Z |
| CHAAL00000010692 | HAO1 | 0.389275074 | Autosome |
| CHAAL00000001818 | GABRR3 | 0.388663968 | Autosome |
| CHAAL00000005262 | CKAP2L | 0.388663968 | Autosome |
| CHAAL00000003538 | HEXB | 0.388663968 | Z |
| CHAAL00000004256 | SYNPR | 0.385964912 | Autosome |
| CHAAL00000011679 | PSIP1 | 0.385964912 | Z |
| CHAAL00000011699 | STX1A | 0.383458647 | Autosome |

| | | | |
|---|---|---|---|
| CHAAL00000014608 | RNF165 | 0.383458647 | Z |
| CHAAL00000000125 | C1QTNF7 | 0.382340148 | Autosome |
| CHAAL00000007031 | PLPP1 | 0.382340148 | Z |
| CHAAL00000011611 | | 0.379305594 | Autosome |
| CHAAL00000007871 | LOC106888475 | 0.376566416 | Autosome |
| CHAAL00000000716 | HDX | 0.373684211 | Autosome |
| CHAAL00000004903 | ARL4A | 0.373684211 | Autosome |
| CHAAL00000008070 | MAEA | 0.373684211 | Autosome |
| CHAAL00000002480 | ZNF131 | 0.371816638 | Z |
| CHAAL00000010078 | HSD17B1 | 0.369806094 | Autosome |
| CHAAL00000001817 | DCAF6 | 0.368421053 | Autosome |
| CHAAL00000001860 | PLA2G12A | 0.368421053 | Autosome |
| CHAAL00000002364 | MAFG | 0.368421053 | Autosome |
| CHAAL00000002863 | DDX11 | 0.368421053 | Autosome |
| CHAAL00000002867 | TMTC1 | 0.368421053 | Autosome |
| CHAAL00000003777 | LOC106893686 | 0.368421053 | Autosome |
| CHAAL00000005263 | SLC20A1 | 0.368421053 | Autosome |
| CHAAL00000006115 | TRIM41 | 0.368421053 | Autosome |
| CHAAL00000007112 | GHSR | 0.368421053 | Autosome |
| CHAAL00000007429 | GOLGA7B | 0.368421053 | Autosome |
| CHAAL00000009165 | MPPE1 | 0.368421053 | Autosome |
| CHAAL00000010145 | ARID3B | 0.368421053 | Autosome |
| CHAAL00000012358 | COMMD6 | 0.368421053 | Autosome |
| CHAAL00000014097 | TACR2 | 0.368421053 | Autosome |
| CHAAL00000015139 | SLC35F3 | 0.368421053 | Autosome |
| CHAAL00000002735 | | 0.368421053 | Z |
| CHAAL00000012446 | PSD3 | 0.368421053 | Z |
| CHAAL00000008624 | DPP7 | 0.366709457 | Autosome |
| CHAAL00000001679 | ZPBP2 | 0.362430466 | Autosome |
| CHAAL00000003410 | DDX23 | 0.358974359 | Autosome |
| CHAAL00000012601 | RPL32 | 0.355410999 | Autosome |
| CHAAL00000015131 | B3GALNT2 | 0.354066986 | Autosome |
| CHAAL00000011353 | LOC106899559 | 0.35356409 | Autosome |
| CHAAL00000004573 | LOC101790034 | 0.352987732 | Autosome |
| CHAAL00000008636 | LOC104050098 | 0.350095887 | Autosome |
| CHAAL00000008764 | VDAC1 | 0.345864662 | Autosome |
| CHAAL00000012305 | PNOC | 0.343859649 | Autosome |
| CHAAL00000006173 | ATG3 | 0.342105263 | Autosome |
| CHAAL00000013793 | | 0.342105263 | Z |
| CHAAL00000013795 | HINT1 | 0.342105263 | Z |
| CHAAL00000012582 | CCDC71 | 0.341317365 | Autosome |
| CHAAL00000009070 | GPR139 | 0.340592861 | Autosome |
| CHAAL00000013156 | PDE4D | 0.338345865 | Z |

| | | | |
|---|---|---|---|
| CHAAL00000013157 | | 0.338345865 | Z |
| CHAAL00000003589 | NLN | 0.336454634 | Z |
| CHAAL00000001528 | LOC104056080 | 0.334014648 | Autosome |
| CHAAL00000010743 | OCSTAMP | 0.332900152 | Autosome |
| CHAAL00000014119 | LRRTM3 | 0.332706767 | Autosome |
| CHAAL00000009195 | MAD2L1BP | 0.33076225 | Autosome |
| CHAAL00000012873 | GPR18 | 0.330481681 | Autosome |
| CHAAL00000009738 | BICRAL | 0.329602286 | Autosome |
| CHAAL00000007258 | LOC104536907 | 0.328718819 | Autosome |
| CHAAL00000003743 | CASK | 0.325162221 | Autosome |
| CHAAL00000003745 | GPR82 | 0.325162221 | Autosome |
| CHAAL00000014882 | HDAC9 | 0.324492333 | Autosome |
| CHAAL00000007187 | KDSR | 0.32132964 | Autosome |
| CHAAL00000010442 | RAB8A | 0.32132964 | Autosome |
| CHAAL00000013387 | PRDM1 | 0.319673229 | Autosome |
| CHAAL00000005519 | PRKCZ | 0.319533142 | Autosome |
| CHAAL00000004833 | LOC104285445 | 0.3189274 | Autosome |
| CHAAL00000011954 | LOC104286445 | 0.317833418 | Autosome |
| CHAAL00000007068 | GRM2 | 0.317524581 | Autosome |
| CHAAL00000013600 | MAPKAP1 | 0.317524581 | Autosome |
| CHAAL00000014789 | | 0.317524581 | Autosome |
| CHAAL00000015463 | | 0.317524581 | Autosome |
| CHAAL00000006604 | CFAP65 | 0.317234048 | Autosome |
| CHAAL00000003373 | ABRA | 0.316770186 | Autosome |
| CHAAL00000008644 | AJM1 | 0.316595059 | Autosome |
| CHAAL00000000109 | LOC104292600 | 0.316520468 | Autosome |
| CHAAL00000000551 | LIN28A | 0.315789474 | Autosome |
| CHAAL00000001028 | TYW3 | 0.315789474 | Autosome |
| CHAAL00000001681 | LOC104313862 | 0.315789474 | Autosome |
| CHAAL00000001950 | EHBP1 | 0.315789474 | Autosome |
| CHAAL00000002153 | LOC104295326 | 0.315789474 | Autosome |
| CHAAL00000003353 | PARM1 | 0.315789474 | Autosome |
| CHAAL00000005614 | RGS2 | 0.315789474 | Autosome |
| CHAAL00000006584 | TMBIM1 | 0.315789474 | Autosome |
| CHAAL00000006677 | NUDT13 | 0.315789474 | Autosome |
| CHAAL00000007827 | DPM2 | 0.315789474 | Autosome |
| CHAAL00000009399 | MTURN | 0.315789474 | Autosome |
| CHAAL00000009429 | CMIP | 0.315789474 | Autosome |
| CHAAL00000010388 | E2F2 | 0.315789474 | Autosome |
| CHAAL00000010944 | LOC104138862 | 0.315789474 | Autosome |
| CHAAL00000012588 | QRICH1 | 0.315789474 | Autosome |
| CHAAL00000013060 | RRAD | 0.315789474 | Autosome |
| CHAAL00000013391 | RTN4IP1 | 0.315789474 | Autosome |

| | | | |
|---|---|---|---|
| CHAAL00000015315 | MASP1 | 0.315789474 | Autosome |
| CHAAL00000001389 | LOC103914478 | 0.315789474 | Z |
| CHAAL00000005738 | LOC104069106 | 0.315789474 | Z |
| CHAAL00000012599 | IFT122 | 0.31547619 | Autosome |
| CHAAL00000009858 | AMFR | 0.313113292 | Autosome |
| CHAAL00000012276 | | 0.313113292 | Z |
| CHAAL00000005773 | CLMN | 0.312886176 | Autosome |
| CHAAL00000007035 | MCIDAS | 0.312030075 | Z |
| CHAAL00000008554 | SLITRK4 | 0.310304586 | Autosome |
| CHAAL00000008626 | MAN1B1 | 0.309409888 | Autosome |
| CHAAL00000012409 | MRM1 | 0.308859337 | Autosome |
| CHAAL00000000714 | LOC104283506 | 0.308652988 | Autosome |
| CHAAL00000012046 | ESD | 0.308652988 | Autosome |
| CHAAL00000015172 | PDCD2 | 0.308603612 | Autosome |
| CHAAL00000008754 | SLC22A5 | 0.308522352 | Autosome |
| CHAAL00000000191 | ZCCHC6 | 0.30778032 | Z |
| CHAAL00000001484 | SYCP2L | 0.307627357 | Autosome |
| CHAAL00000007893 | MED27 | 0.307479224 | Autosome |
| CHAAL00000004511 | DCLRE1C | 0.304565448 | Autosome |
| CHAAL00000006877 | NPY5R | 0.304132815 | Autosome |
| CHAAL00000010813 | ZNF335 | 0.302631579 | Autosome |
| CHAAL00000002733 | NFIL3 | 0.302631579 | Z |
| CHAAL00000009603 | SMG7 | 0.302131912 | Autosome |
| CHAAL00000004878 | ZC3H12A | 0.301075269 | Autosome |
| CHAAL00000006768 | GCM1 | 0.301075269 | Autosome |
| CHAAL00000009642 | TBX4 | 0.300965592 | Autosome |
| CHAAL00000005233 | MUC2 | 0.300875878 | Autosome |
| CHAAL00000007589 | PEX26 | 0.300825593 | Autosome |
| CHAAL00000011302 | | 0.300205822 | Autosome |
| CHAAL00000012704 | SLC17A9 | 0.299579748 | Autosome |
| CHAAL00000008017 | | 0.298885901 | Autosome |
| CHAAL00000003647 | DDR2 | 0.298245614 | Autosome |
| CHAAL00000006192 | ZDHHC23 | 0.298245614 | Autosome |
| CHAAL00000009649 | VMP1 | 0.298245614 | Autosome |
| CHAAL00000003146 | | 0.297195084 | Autosome |
| CHAAL00000003490 | E2F3 | 0.296936371 | Autosome |
| CHAAL00000007245 | LOC104286679 | 0.296650718 | Autosome |
| CHAAL00000013023 | PENK | 0.296650718 | Autosome |
| CHAAL00000013149 | ZSWIM6 | 0.296650718 | Z |
| CHAAL00000005701 | QSOX1 | 0.296398892 | Autosome |
| CHAAL00000008223 | | 0.295921924 | Z |
| CHAAL00000004228 | MON1A | 0.295587453 | Autosome |
| CHAAL00000009194 | GTPBP2 | 0.294871795 | Autosome |

| | | | |
|---|---|---|---|
| CHAAL00000012584 | USP19 | 0.294153752 | Autosome |
| CHAAL00000008050 | SH3BP2 | 0.294043492 | Autosome |
| CHAAL00000001348 | CRMP1 | 0.293921423 | Autosome |
| CHAAL00000005742 | BDP1 | 0.293410692 | Z |
| CHAAL00000010954 | TOP3B | 0.292382628 | Autosome |
| CHAAL00000005126 | NUAK1 | 0.292318634 | Autosome |
| CHAAL00000008443 | LOC106890872 | 0.291696826 | Autosome |
| CHAAL00000008305 | BIRC5 | 0.291146943 | Autosome |
| CHAAL00000010451 | MED26 | 0.288651316 | Autosome |
| CHAAL00000000061 | LOC106855934 | 0.287925697 | Autosome |
| CHAAL00000013736 | | 0.287925697 | Autosome |
| CHAAL00000003890 | DRD1 | 0.287509819 | Autosome |
| CHAAL00000007352 | DUSP5 | 0.285714286 | Autosome |
| CHAAL00000003094 | LOC106895940 | 0.284210526 | Autosome |
| CHAAL00000004434 | COL15A1 | 0.284210526 | Autosome |
| CHAAL00000005658 | ITPA | 0.284210526 | Autosome |
| CHAAL00000011166 | CALCRL | 0.284210526 | Autosome |
| CHAAL00000012598 | RHO | 0.284210526 | Autosome |
| CHAAL00000010906 | TMEM132C | 0.283911377 | Autosome |
| CHAAL00000007716 | UTP20 | 0.282416568 | Autosome |
| CHAAL00000013199 | LIFR | 0.282296651 | Z |
| CHAAL00000001997 | LOC103540408 | 0.281464531 | Autosome |
| CHAAL00000005440 | SPAG6 | 0.279581934 | Autosome |
| CHAAL00000006340 | LOC101913524 | 0.278515438 | Autosome |
| CHAAL00000005235 | MUC6 | 0.275541796 | Autosome |
| CHAAL00000013760 | TNRC6A | 0.27548998 | Autosome |
| CHAAL00000013483 | CCDC148 | 0.274498401 | Autosome |
| CHAAL00000001935 | SPRED2 | 0.274272638 | Autosome |
| CHAAL00000000450 | BNIP1 | 0.271929825 | Autosome |
| CHAAL00000006515 | DCAF5 | 0.271929825 | Autosome |
| CHAAL00000014250 | METTL16 | 0.271929825 | Autosome |
| CHAAL00000009517 | NOB1 | 0.271895153 | Autosome |
| CHAAL00000004770 | | 0.271539504 | Autosome |
| CHAAL00000014957 | | 0.269680464 | Autosome |
| CHAAL00000000695 | | 0.268925739 | Autosome |
| CHAAL00000007274 | | 0.268782644 | Autosome |
| CHAAL00000006862 | NEK1 | 0.268368942 | Autosome |
| CHAAL00000005700 | LHX4 | 0.268292683 | Autosome |
| CHAAL00000000233 | ADAMTS17 | 0.26715523 | Autosome |
| CHAAL00000010210 | OTUD7A | 0.265143992 | Autosome |
| CHAAL00000003991 | LOC104289750 | 0.264599856 | Autosome |
| CHAAL00000005662 | LOC104293403 | 0.264321024 | Autosome |
| CHAAL00000000093 | PLEKHA7 | 0.264132554 | Autosome |

| | | | |
|---|---|---|---|
| CHAAL00000000602 | LOC101911571 | 0.264132554 | Autosome |
| CHAAL00000000689 | IGBP1 | 0.264132554 | Autosome |
| CHAAL00000004880 | CSF3R | 0.263733103 | Autosome |
| CHAAL00000015123 | DNAH1 | 0.263335404 | Autosome |
| CHAAL00000000039 | PIH1D2 | 0.263157895 | Autosome |
| CHAAL00000000094 | RPS13 | 0.263157895 | Autosome |
| CHAAL00000000232 | CERS3 | 0.263157895 | Autosome |
| CHAAL00000000736 | SYTL4 | 0.263157895 | Autosome |
| CHAAL00000000746 | FGF13 | 0.263157895 | Autosome |
| CHAAL00000001326 | | 0.263157895 | Autosome |
| CHAAL00000001485 | ELOVL2 | 0.263157895 | Autosome |
| CHAAL00000002018 | GRPR | 0.263157895 | Autosome |
| CHAAL00000002224 | HNRNPR | 0.263157895 | Autosome |
| CHAAL00000002794 | JPT2 | 0.263157895 | Autosome |
| CHAAL00000002819 | HOXA7 | 0.263157895 | Autosome |
| CHAAL00000003193 | DNAJA2 | 0.263157895 | Autosome |
| CHAAL00000003672 | SLC35A1 | 0.263157895 | Autosome |
| CHAAL00000003727 | CACNA2D2 | 0.263157895 | Autosome |
| CHAAL00000003821 | VGLL4 | 0.263157895 | Autosome |
| CHAAL00000003903 | DBN1 | 0.263157895 | Autosome |
| CHAAL00000004552 | GFRA1 | 0.263157895 | Autosome |
| CHAAL00000004633 | KAZN | 0.263157895 | Autosome |
| CHAAL00000005379 | GAD1 | 0.263157895 | Autosome |
| CHAAL00000005450 | | 0.263157895 | Autosome |
| CHAAL00000005506 | LOC104295600 | 0.263157895 | Autosome |
| CHAAL00000005706 | FAM163A | 0.263157895 | Autosome |
| CHAAL00000006632 | INHA | 0.263157895 | Autosome |
| CHAAL00000007518 | WNT5B | 0.263157895 | Autosome |
| CHAAL00000007884 | FIBCD1 | 0.263157895 | Autosome |
| CHAAL00000007898 | | 0.263157895 | Autosome |
| CHAAL00000007922 | FAM163B | 0.263157895 | Autosome |
| CHAAL00000009833 | PIFO | 0.263157895 | Autosome |
| CHAAL00000009903 | MBOAT2 | 0.263157895 | Autosome |
| CHAAL00000010832 | LOC101911060 | 0.263157895 | Autosome |
| CHAAL00000011123 | CPSF6 | 0.263157895 | Autosome |
| CHAAL00000011138 | LRRTM1 | 0.263157895 | Autosome |
| CHAAL00000013062 | CA7 | 0.263157895 | Autosome |
| CHAAL00000013084 | LOC106888755 | 0.263157895 | Autosome |
| CHAAL00000013242 | LOC102090344 | 0.263157895 | Autosome |
| CHAAL00000013587 | NEK6 | 0.263157895 | Autosome |
| CHAAL00000013716 | LOC104283874 | 0.263157895 | Autosome |
| CHAAL00000014334 | TSPAN32 | 0.263157895 | Autosome |
| CHAAL00000015202 | TNIP3 | 0.263157895 | Autosome |

| | | | |
|---|---|---|---|
| CHAAL00000015290 | COPS7B | 0.263157895 | Autosome |
| CHAAL00000003442 | PTGER4 | 0.263157895 | Z |
| CHAAL00000008267 | INIP | 0.263157895 | Z |
| CHAAL00000012452 | TAL2 | 0.263157895 | Z |
| CHAAL00000013790 | CHSY3 | 0.263157895 | Z |
| CHAAL00000010238 | ALDH1A2 | 0.262858852 | Autosome |
| CHAAL00000010056 | IFI35 | 0.26274831 | Autosome |
| CHAAL00000011964 | RMDN2 | 0.262527576 | Autosome |
| CHAAL00000001536 | LAMB1 | 0.262207437 | Autosome |
| CHAAL00000004144 | RHOC | 0.262174127 | Autosome |
| CHAAL00000004638 | LOC103907554 | 0.262174127 | Autosome |
| CHAAL00000013117 | CRHR2 | 0.262174127 | Autosome |
| CHAAL00000013606 | | 0.262174127 | Autosome |
| CHAAL00000006240 | RCL1 | 0.262174127 | Z |
| CHAAL00000000732 | TNMD | 0.261874198 | Autosome |
| CHAAL00000002240 | LOC104292733, LOC104056669 | 0.261874198 | Autosome |
| CHAAL00000002897 | BRAF | 0.261874198 | Autosome |
| CHAAL00000008105 | LOC104020516 | 0.261874198 | Autosome |
| CHAAL00000008467 | TCIM | 0.261874198 | Autosome |
| CHAAL00000014927 | LOC101878833 | 0.261874198 | Autosome |
| CHAAL00000008691 | PROSER2 | 0.261369985 | Autosome |
| CHAAL00000002219 | RPA2 | 0.260792431 | Autosome |
| CHAAL00000004519 | EXOC4 | 0.26057475 | Autosome |
| CHAAL00000002113 | GGACT | 0.259868421 | Autosome |
| CHAAL00000000637 | ICE1 | 0.259734703 | Autosome |
| CHAAL00000008763 | LOC104291590 | 0.259649123 | Autosome |
| CHAAL00000001916 | IRAK4 | 0.259485924 | Autosome |
| CHAAL00000001974 | | 0.258977769 | Autosome |
| CHAAL00000002872 | DENND5B | 0.258960284 | Autosome |
| CHAAL00000001058 | N4BP2L1 | 0.258373206 | Autosome |
| CHAAL00000003895 | N4BP3 | 0.258373206 | Autosome |
| CHAAL00000001010 | EHHADH | 0.257492172 | Autosome |
| CHAAL00000002884 | SYT10 | 0.257240735 | Autosome |
| CHAAL00000001537 | DLD | 0.256965944 | Autosome |
| CHAAL00000009778 | LOC104289550 | 0.255898367 | Autosome |
| CHAAL00000015357 | LARP4B | 0.255715045 | Autosome |
| CHAAL00000010519 | ADSL | 0.254639489 | Autosome |
| CHAAL00000007908 | RALGDS | 0.253056885 | Autosome |
| CHAAL00000003293 | ARFIP1 | 0.252631579 | Autosome |
| CHAAL00000006153 | TBC1D23 | 0.252631579 | Autosome |
| CHAAL00000008140 | MESD | 0.252631579 | Autosome |
| CHAAL00000010723 | TOMM34 | 0.252631579 | Autosome |
| CHAAL00000012580 | KLHDC8B | 0.252631579 | Autosome |

| | | | |
|---|---|---|---|
| CHAAL00000006310 | PLBD1 | 0.25229741 | Autosome |
| CHAAL00000002638 | | 0.250398724 | Autosome |
| CHAAL00000003280 | FGG | 0.250355619 | Autosome |
| CHAAL00000013621 | GDI2 | 0.250116442 | Autosome |
| CHAAL00000011428 | LRRC6 | 0.24984876 | Autosome |
| CHAAL00000012733 | | 0.249512671 | Autosome |
| CHAAL00000014345 | CTSD | 0.249011858 | Autosome |
| CHAAL00000003462 | LPXN | 0.248120301 | Autosome |
| CHAAL00000004801 | ADTRP | 0.248120301 | Autosome |
| CHAAL00000008287 | LOC104285906 | 0.248120301 | Autosome |
| CHAAL00000013255 | CRCP | 0.248120301 | Autosome |
| CHAAL00000003274 | GUCY1A1 | 0.24769551 | Autosome |
| CHAAL00000013445 | LOC104023457 | 0.247439867 | Autosome |
| CHAAL00000012099 | | 0.247076023 | Autosome |
| CHAAL00000010946 | CDC45 | 0.246923919 | Autosome |
| CHAAL00000000524 | LOC101870611 | 0.246920493 | Autosome |
| CHAAL00000013526 | E2F7 | 0.246088193 | Autosome |
| CHAAL00000008494 | SPERT | 0.245961794 | Autosome |
| CHAAL00000006411 | ABCD4 | 0.2457949 | Autosome |
| CHAAL00000000803 | | 0.24548456 | Autosome |
| CHAAL00000007719 | GAS2L3 | 0.24534413 | Autosome |
| CHAAL00000001968 | DENND4C | 0.245215311 | Autosome |
| CHAAL00000009258 | PLEKHH2 | 0.245069715 | Autosome |
| CHAAL00000001512 | ARHGAP25 | 0.244912281 | Autosome |
| CHAAL00000015501 | LOC104290743 | 0.244748643 | Autosome |
| CHAAL00000001962 | HACD4 | 0.244482173 | Autosome |
| CHAAL00000007568 | PLCZ1 | 0.244457849 | Autosome |
| CHAAL00000003908 | | 0.244455478 | Autosome |
| CHAAL00000015665 | LOC104630663 | 0.244326412 | Autosome |
| CHAAL00000004147 | ST7L | 0.243978591 | Autosome |
| CHAAL00000008814 | TBC1D9B | 0.243521648 | Autosome |
| CHAAL00000011721 | RNF43 | 0.243421053 | Autosome |
| CHAAL00000013437 | HMOX2 | 0.24297044 | Autosome |
| CHAAL00000009092 | FOPNL | 0.242881795 | Autosome |
| CHAAL00000004705 | HADHB | 0.242470338 | Autosome |
| CHAAL00000004799 | EDN1 | 0.24099723 | Autosome |
| CHAAL00000009943 | CABLES1 | 0.24099723 | Autosome |
| CHAAL00000006018 | SLC25A24 | 0.240551042 | Autosome |
| CHAAL00000011584 | MADD | 0.240414055 | Autosome |
| CHAAL00000004923 | CLVS2 | 0.2402746 | Autosome |
| CHAAL00000008903 | PRELID1 | 0.2402746 | Autosome |
| CHAAL00000014485 | LOC104287055 | 0.2402746 | Autosome |
| CHAAL00000004328 | NCLN | 0.239766082 | Autosome |

| | | | |
|---|---|---|---|
| CHAAL00000005772 | DICER1 | 0.239489256 | Autosome |
| CHAAL00000012016 | ACTR5 | 0.239486275 | Autosome |
| CHAAL00000009935 | USP14 | 0.23923445 | Autosome |
| CHAAL00000011206 | EPS15 | 0.23923445 | Autosome |
| CHAAL00000014107 | TET1 | 0.239104549 | Autosome |
| CHAAL00000009441 | ATP2C2 | 0.238721805 | Autosome |
| CHAAL00000004033 | RAB44 | 0.238693063 | Autosome |
| CHAAL00000000670 | LOC104282587 | 0.238296747 | Autosome |
| CHAAL00000008752 | LOC104292002 | 0.237867396 | Autosome |
| CHAAL00000004838 | TMEM179 | 0.237749546 | Autosome |
| CHAAL00000007617 | MAK16 | 0.237355863 | Autosome |
| CHAAL00000013413 | LOC104294499 | 0.236842105 | Autosome |
| CHAAL00000011585 | MYBPC3 | 0.236744999 | Autosome |
| CHAAL00000010164 | SNX33 | 0.23647146 | Autosome |
| CHAAL00000005623 | | 0.235944976 | Autosome |
| CHAAL00000005061 | FAM46B | 0.235518965 | Autosome |
| CHAAL00000002458 | RIPK4 | 0.234992243 | Autosome |
| CHAAL00000013013 | TMEM68, LOC106894137 | 0.234851835 | Autosome |
| CHAAL00000013116 | MINDY4 | 0.234851835 | Autosome |
| CHAAL00000004394 | LOC101867830 | 0.234834073 | Autosome |
| CHAAL00000003971 | LOC105412838 | 0.234708393 | Autosome |
| CHAAL00000007132 | LOC101915456 | 0.234106772 | Autosome |
| CHAAL00000014767 | PICK1 | 0.233808382 | Autosome |
| CHAAL00000001019 | STAG1 | 0.233798783 | Autosome |
| CHAAL00000003841 | SORBS2 | 0.232813304 | Autosome |
| CHAAL00000000699 | LOC103906158 | 0.23231646 | Autosome |
| CHAAL00000014149 | NFXL1 | 0.232096635 | Autosome |
| CHAAL00000015360 | | 0.232096635 | Autosome |
| CHAAL00000008690 | ECHDC3 | 0.230769231 | Autosome |
| CHAAL00000003873 | LOC101870911 | 0.230628655 | Autosome |
| CHAAL00000006531 | FZD6 | 0.230549199 | Autosome |
| CHAAL00000002096 | FKBP9 | 0.229916898 | Autosome |
| CHAAL00000008012 | UROC1 | 0.229808267 | Autosome |
| CHAAL00000012694 | SOD3 | 0.229323308 | Autosome |
| CHAAL00000005275 | LOC104842678 | 0.228070175 | Autosome |
| CHAAL00000001539 | CBLL1 | 0.227892065 | Autosome |
| CHAAL00000004041 | PNPLA1 | 0.227814061 | Autosome |
| CHAAL00000012786 | WDR12 | 0.227814061 | Autosome |
| CHAAL00000000707 | | 0.227688787 | Autosome |
| CHAAL00000002360 | ARHGDIA | 0.227688787 | Autosome |
| CHAAL00000009364 | EXOSC7 | 0.227688787 | Autosome |
| CHAAL00000009740 | PRPH2 | 0.227688787 | Autosome |
| CHAAL00000011292 | ARMT1 | 0.227688787 | Autosome |

| | | | |
|---|---|---|---|
| CHAAL00000013734 | DHRSX | 0.227688787 | Autosome |
| CHAAL00000014821 | CCDC9B | 0.227688787 | Autosome |
| CHAAL00000014019 | NF2 | 0.227395412 | Autosome |
| CHAAL00000008789 | LOC104291817 | 0.227121859 | Autosome |
| CHAAL00000005577 | | 0.226315789 | Autosome |
| CHAAL00000005578 | LOC104072859 | 0.226315789 | Autosome |
| CHAAL00000011045 | PRKAB1 | 0.226147561 | Autosome |
| CHAAL00000002033 | | 0.223832052 | Autosome |
| CHAAL00000000673 | | 0.223766965 | Autosome |
| CHAAL00000013894 | HLCS | 0.222939424 | Autosome |
| CHAAL00000002981 | ANXA7 | 0.22225947 | Autosome |
| CHAAL00000001840 | LOC104282992 | 0.221926658 | Autosome |
| CHAAL00000009951 | OSBPL1A | 0.221599792 | Autosome |
| CHAAL00000003311 | DUSP4 | 0.221491228 | Autosome |
| CHAAL00000014772 | TMEM184B | 0.221298861 | Autosome |
| CHAAL00000015280 | LOC104284949 | 0.221113128 | Autosome |
| CHAAL00000014529 | BNIPL | 0.221052632 | Autosome |
| CHAAL00000011882 | EPHX2 | 0.220519654 | Autosome |
| CHAAL00000003012 | CTDSPL2 | 0.22034564 | Autosome |
| CHAAL00000013723 | | 0.22008547 | Autosome |
| CHAAL00000013660 | | 0.219497608 | Autosome |
| CHAAL00000012539 | PRPF39 | 0.219278796 | Autosome |
| CHAAL00000008453 | RNF219 | 0.218623482 | Autosome |
| CHAAL00000004359 | TEX30 | 0.218298912 | Autosome |
| CHAAL00000014674 | | 0.218117409 | Autosome |
| CHAAL00000002868 | IPO8 | 0.217865381 | Autosome |
| CHAAL00000006371 | LOC104288274 | 0.217638691 | Autosome |
| CHAAL00000005247 | SUPT20H | 0.217415266 | Autosome |
| CHAAL00000009057 | ZP2 | 0.217284456 | Autosome |
| CHAAL00000002686 | | 0.216900382 | Autosome |
| CHAAL00000014401 | ATG16L1 | 0.216882787 | Autosome |
| CHAAL00000008020 | | 0.216696915 | Autosome |
| CHAAL00000001094 | SPATA13 | 0.215631672 | Autosome |
| CHAAL00000006579 | TNS1 | 0.21557318 | Autosome |
| CHAAL00000000401 | | 0.215097297 | Autosome |
| CHAAL00000011608 | ALKBH3 | 0.21484038 | Autosome |
| CHAAL00000002957 | THSD1 | 0.214058637 | Autosome |
| CHAAL00000000365 | | 0.213775179 | Autosome |
| CHAAL00000012501 | FAM20A | 0.213775179 | Autosome |
| CHAAL00000000106 | SAAL1 | 0.213735558 | Autosome |
| CHAAL00000006678 | ECD | 0.213586291 | Autosome |
| CHAAL00000000128 | BST1 | 0.213528117 | Autosome |
| CHAAL00000009485 | SPG7 | 0.213483146 | Autosome |

| | | | |
|---|---|---|---|
| CHAAL00000012107 | | 0.213390186 | Autosome |
| CHAAL00000005635 | | 0.212598425 | Autosome |
| CHAAL00000010708 | ESF1 | 0.212436025 | Autosome |
| CHAAL00000005387 | FASTKD1 | 0.212181397 | Autosome |
| CHAAL00000003691 | ME1 | 0.212121212 | Autosome |
| CHAAL00000005207 | WNK2 | 0.212065251 | Autosome |
| CHAAL00000011039 | PXN | 0.212013084 | Autosome |
| CHAAL00000011601 | TSPAN18 | 0.21172795 | Autosome |
| CHAAL00000000058 | TIGAR | 0.210526316 | Autosome |
| CHAAL00000000355 | DMAP1 | 0.210526316 | Autosome |
| CHAAL00000000571 | MAN1C1 | 0.210526316 | Autosome |
| CHAAL00000000585 | TRAF6 | 0.210526316 | Autosome |
| CHAAL00000000664 | ZC4H2 | 0.210526316 | Autosome |
| CHAAL00000000684 | EFNB1 | 0.210526316 | Autosome |
| CHAAL00000000791 | MYD88 | 0.210526316 | Autosome |
| CHAAL00000001090 | SHISA2 | 0.210526316 | Autosome |
| CHAAL00000001235 | DNAJA3 | 0.210526316 | Autosome |
| CHAAL00000001495 | LOC104289569 | 0.210526316 | Autosome |
| CHAAL00000001509 | TMEM230 | 0.210526316 | Autosome |
| CHAAL00000001612 | SNF8 | 0.210526316 | Autosome |
| CHAAL00000001615 | GNGT2 | 0.210526316 | Autosome |
| CHAAL00000001656 | SERBP1 | 0.210526316 | Autosome |
| CHAAL00000001683 | PSMD3 | 0.210526316 | Autosome |
| CHAAL00000001908 | DBX2 | 0.210526316 | Autosome |
| CHAAL00000001954 | COMMD1 | 0.210526316 | Autosome |
| CHAAL00000002314 | UBXN7 | 0.210526316 | Autosome |
| CHAAL00000002811 | SNX10 | 0.210526316 | Autosome |
| CHAAL00000003350 | AP1AR | 0.210526316 | Autosome |
| CHAAL00000003797 | MCM3AP | 0.210526316 | Autosome |
| CHAAL00000003885 | LSM6 | 0.210526316 | Autosome |
| CHAAL00000004324 | GNA11 | 0.210526316 | Autosome |
| CHAAL00000004715 | AAGAB | 0.210526316 | Autosome |
| CHAAL00000005085 | TSEN54 | 0.210526316 | Autosome |
| CHAAL00000005157 | DLGAP3 | 0.210526316 | Autosome |
| CHAAL00000005726 | SREBF1 | 0.210526316 | Autosome |
| CHAAL00000005849 | XKR9 | 0.210526316 | Autosome |
| CHAAL00000005917 | LOC104286147 | 0.210526316 | Autosome |
| CHAAL00000006031 | NR5A2 | 0.210526316 | Autosome |
| CHAAL00000006094 | LOC103907365 | 0.210526316 | Autosome |
| CHAAL00000006165 | TBX19 | 0.210526316 | Autosome |
| CHAAL00000006335 | LOC107312618 | 0.210526316 | Autosome |
| CHAAL00000006359 | ADCK1 | 0.210526316 | Autosome |
| CHAAL00000006650 | FIGN | 0.210526316 | Autosome |

| | | | |
|---|---|---|---|
| CHAAL00000006688 | | 0.210526316 | Autosome |
| CHAAL00000007209 | NQO2 | 0.210526316 | Autosome |
| CHAAL00000007392 | TWNK | 0.210526316 | Autosome |
| CHAAL00000007566 | AEBP2 | 0.210526316 | Autosome |
| CHAAL00000007673 | LOC104282668 | 0.210526316 | Autosome |
| CHAAL00000007774 | SHANK3 | 0.210526316 | Autosome |
| CHAAL00000008994 | GRXCR2 | 0.210526316 | Autosome |
| CHAAL00000009084 | ARL6IP1 | 0.210526316 | Autosome |
| CHAAL00000009438 | HSBP1 | 0.210526316 | Autosome |
| CHAAL00000009466 | BANP | 0.210526316 | Autosome |
| CHAAL00000009597 | TSEN15 | 0.210526316 | Autosome |
| CHAAL00000009605 | LAMC2 | 0.210526316 | Autosome |
| CHAAL00000009695 | ADORA2B | 0.210526316 | Autosome |
| CHAAL00000009812 | | 0.210526316 | Autosome |
| CHAAL00000009852 | WDR92 | 0.210526316 | Autosome |
| CHAAL00000009929 | LOC101876247 | 0.210526316 | Autosome |
| CHAAL00000009945 | RMC1 | 0.210526316 | Autosome |
| CHAAL00000009998 | MYCL | 0.210526316 | Autosome |
| CHAAL00000010036 | MUL1 | 0.210526316 | Autosome |
| CHAAL00000010141 | SEMA7A | 0.210526316 | Autosome |
| CHAAL00000010336 | ICMT | 0.210526316 | Autosome |
| CHAAL00000010613 | VIPR1 | 0.210526316 | Autosome |
| CHAAL00000010644 | LOC104533797 | 0.210526316 | Autosome |
| CHAAL00000010693 | TMX4 | 0.210526316 | Autosome |
| CHAAL00000010709 | NDUFAF5 | 0.210526316 | Autosome |
| CHAAL00000010932 | RANBP1 | 0.210526316 | Autosome |
| CHAAL00000011049 | SRRM4 | 0.210526316 | Autosome |
| CHAAL00000011174 | NOX5 | 0.210526316 | Autosome |
| CHAAL00000011257 | JUN | 0.210526316 | Autosome |
| CHAAL00000011551 | CHAC1 | 0.210526316 | Autosome |
| CHAAL00000011560 | CHST1 | 0.210526316 | Autosome |
| CHAAL00000011567 | CREB3L1 | 0.210526316 | Autosome |
| CHAAL00000011807 | LOC105402709 | 0.210526316 | Autosome |
| CHAAL00000012106 | SOCS6 | 0.210526316 | Autosome |
| CHAAL00000012203 | TRIB1 | 0.210526316 | Autosome |
| CHAAL00000012240 | TNFRSF11B | 0.210526316 | Autosome |
| CHAAL00000012311 | DNAJC27 | 0.210526316 | Autosome |
| CHAAL00000012581 | KLHDC8B | 0.210526316 | Autosome |
| CHAAL00000012811 | MAIP1 | 0.210526316 | Autosome |
| CHAAL00000013053 | GPI | 0.210526316 | Autosome |
| CHAAL00000013098 | ABTB1 | 0.210526316 | Autosome |
| CHAAL00000013316 | PIPOX | 0.210526316 | Autosome |
| CHAAL00000013339 | GATSL2 | 0.210526316 | Autosome |

| | | | |
|---|---|---|---|
| CHAAL00000013578 | INPP5A | 0.210526316 | Autosome |
| CHAAL00000013591 | NR6A1 | 0.210526316 | Autosome |
| CHAAL00000013846 | LOC104291507 | 0.210526316 | Autosome |
| CHAAL00000013898 | VPS26C | 0.210526316 | Autosome |
| CHAAL00000014055 | DPAGT1 | 0.210526316 | Autosome |
| CHAAL00000015391 | ANKRD40 | 0.210526316 | Autosome |
| CHAAL00000015576 | LOC104285043 | 0.210526316 | Autosome |
| CHAAL00000015668 | | 0.210526316 | Autosome |
| CHAAL00000003860 | CDKN2AIP | 0.209996823 | Autosome |
| CHAAL00000006708 | TMEM200A | 0.209634255 | Autosome |
| CHAAL00000000085 | COPB1 | 0.209544383 | Autosome |
| CHAAL00000013375 | LOC106894421 | 0.209356725 | Autosome |
| CHAAL00000001483 | GCM2 | 0.208850151 | Autosome |
| CHAAL00000004089 | TMEM130 | 0.208605455 | Autosome |
| CHAAL00000000693 | ARR3 | 0.208386821 | Autosome |
| CHAAL00000002185 | MMAB | 0.208386821 | Autosome |
| CHAAL00000005439 | PIP4K2A | 0.208066896 | Autosome |
| CHAAL00000014714 | LOC104031724 | 0.208008058 | Autosome |
| CHAAL00000011090 | LOC104282471 | 0.207767847 | Autosome |
| CHAAL00000010630 | ATP5F1C | 0.207741576 | Autosome |
| CHAAL00000002557 | LRIT2 | 0.207393484 | Autosome |
| CHAAL00000008795 | WNT8A | 0.207393484 | Autosome |
| CHAAL00000001014 | DBR1, LOC106886629 | 0.207277453 | Autosome |
| CHAAL00000003398 | SCN8A | 0.206993995 | Autosome |
| CHAAL00000001171 | GTDC1 | 0.206921413 | Autosome |
| CHAAL00000001912 | ARRDC5 | 0.206921413 | Autosome |
| CHAAL00000001953 | B3GNT2 | 0.206921413 | Autosome |
| CHAAL00000002218 | SUSD2 | 0.206921413 | Autosome |
| CHAAL00000004072 | LOC104573748 | 0.206921413 | Autosome |
| CHAAL00000004120 | NGF | 0.206921413 | Autosome |
| CHAAL00000004125 | CSDE1 | 0.206921413 | Autosome |
| CHAAL00000005032 | MRPS22 | 0.206921413 | Autosome |
| CHAAL00000005384 | SSB | 0.206921413 | Autosome |
| CHAAL00000005599 | ATP13A5 | 0.206921413 | Autosome |
| CHAAL00000007952 | CUTA | 0.206921413 | Autosome |
| CHAAL00000010156 | SCAMP5 | 0.206921413 | Autosome |
| CHAAL00000011698 | BUD23 | 0.206921413 | Autosome |
| CHAAL00000011789 | TAGLN2 | 0.206921413 | Autosome |
| CHAAL00000013386 | PREP | 0.206921413 | Autosome |
| CHAAL00000015068 | LOC104282741 | 0.206921413 | Autosome |
| CHAAL00000005586 | GPR132 | 0.206721623 | Autosome |
| CHAAL00000013106 | FBXO47 | 0.206644931 | Autosome |
| CHAAL00000002565 | LOC104046051 | 0.206539075 | Autosome |

| | | | |
|---|---|---|---|
| CHAAL00000003017 | CTSH | 0.206539075 | Autosome |
| CHAAL00000010686 | XPO1 | 0.206539075 | Autosome |
| CHAAL00000000048 | NCK2 | 0.20602789 | Autosome |
| CHAAL00000010408 | CFAP57 | 0.205949657 | Autosome |
| CHAAL00000005217 | LOC104264865 | 0.205798928 | Autosome |
| CHAAL00000005469 | FAM171A1 | 0.205719779 | Autosome |
| CHAAL00000004278 | INTS4 | 0.205709782 | Autosome |
| CHAAL00000007958 | GSN | 0.205706574 | Autosome |
| CHAAL00000013351 | ANKFY1 | 0.205626333 | Autosome |
| CHAAL00000007759 | DCN | 0.205607477 | Autosome |
| CHAAL00000009115 | LOC103917624 | 0.205523314 | Autosome |
| CHAAL00000001135 | POLE2 | 0.205263158 | Autosome |
| CHAAL00000001628 | TMPPE | 0.205263158 | Autosome |
| CHAAL00000010409 | EBNA1BP2 | 0.205263158 | Autosome |
| CHAAL00000012556 | LSM3 | 0.205263158 | Autosome |
| CHAAL00000012761 | CD28 | 0.205263158 | Autosome |
| CHAAL00000015268 | ABCF3 | 0.205263158 | Autosome |
| CHAAL00000007811 | CCDC102A | 0.204742626 | Autosome |
| CHAAL00000001752 | OTULINL | 0.204645692 | Autosome |
| CHAAL00000009335 | CLRN1 | 0.204499799 | Autosome |
| CHAAL00000009397 | METTL6 | 0.204334365 | Autosome |
| CHAAL00000014299 | SLC4A11 | 0.204334365 | Autosome |
| CHAAL00000010775 | ATP9A | 0.203778677 | Autosome |
| CHAAL00000002702 | ALAS1 | 0.203708754 | Autosome |
| CHAAL00000000522 | GUK1 | 0.203375286 | Autosome |
| CHAAL00000002576 | ANXA11 | 0.203040424 | Autosome |
| CHAAL00000010974 | | 0.203007519 | Autosome |
| CHAAL00000012594 | RBSN | 0.202620285 | Autosome |
| CHAAL00000006135 | ZPLD1 | 0.202361512 | Autosome |
| CHAAL00000002966 | SERPINE3 | 0.202037351 | Autosome |
| CHAAL00000015494 | MYO1G | 0.201575367 | Autosome |
| CHAAL00000014056 | VPS11 | 0.201499792 | Autosome |
| CHAAL00000015196 | PDE5A | 0.201451906 | Autosome |
| CHAAL00000005863 | MTM1 | 0.200932018 | Autosome |
| CHAAL00000009959 | TAF4B | 0.200827622 | Autosome |
| CHAAL00000006805 | PRADC1 | 0.200779727 | Autosome |
| CHAAL00000007445 | LOC107206690 | 0.200779727 | Autosome |
| CHAAL00000009832 | LOC104289366 | 0.200469326 | Autosome |
| CHAAL00000009383 | NPHP3 | 0.200316185 | Autosome |
| CHAAL00000012413 | LOC101877420 | 0.200150915 | Autosome |
| CHAAL00000009181 | TTC27 | 0.199893674 | Autosome |
| CHAAL00000012197 | ADCY8 | 0.19984463 | Autosome |
| CHAAL00000013809 | MYNN | 0.19982884 | Autosome |

| | | | |
|---|---|---|---|
| CHAAL00000003218 | | 0.199561404 | Autosome |
| CHAAL00000002893 | KIAA1147 | 0.199207697 | Autosome |
| CHAAL00000011691 | SERPINH1 | 0.199207697 | Autosome |
| CHAAL00000003925 | ALKBH8 | 0.199174407 | Autosome |
| CHAAL00000003384 | LOC104258112 | 0.199127763 | Autosome |
| CHAAL00000011147 | MYO1B | 0.199124327 | Autosome |
| CHAAL00000001884 | LOC103894282 | 0.199084668 | Autosome |
| CHAAL00000008629 | ENTPD2 | 0.197994987 | Autosome |
| CHAAL00000006042 | KCNT2 | 0.197909156 | Autosome |
| CHAAL00000011014 | PISD | 0.197644461 | Autosome |
| CHAAL00000001321 | LOC104288922 | 0.197584124 | Autosome |
| CHAAL00000006568 | PECR | 0.197584124 | Autosome |
| CHAAL00000009374 | CXCR6 | 0.197584124 | Autosome |
| CHAAL00000009993 | HEYL | 0.197584124 | Autosome |
| CHAAL00000012366 | DACH1 | 0.197584124 | Autosome |
| CHAAL00000006092 | LOC104264308 | 0.197407007 | Autosome |
| CHAAL00000007930 | MRPS2 | 0.197251288 | Autosome |
| CHAAL00000005386 | PPIG | 0.197031039 | Autosome |
| CHAAL00000002996 | MPP7 | 0.196936002 | Autosome |
| CHAAL00000012007 | LPIN3 | 0.196756426 | Autosome |
| CHAAL00000012032 | CBFA2T2 | 0.196528555 | Autosome |
| CHAAL00000007564 | SLCO1C1 | 0.195996125 | Autosome |
| CHAAL00000002327 | INTS8 | 0.195906433 | Autosome |
| CHAAL00000010059 | G6PC | 0.195700519 | Autosome |
| CHAAL00000012851 | DCT | 0.195557702 | Autosome |
| CHAAL00000012623 | | 0.195488722 | Autosome |
| CHAAL00000003089 | LOC104293991 | 0.195302305 | Autosome |
| CHAAL00000015667 | | 0.19504644 | Autosome |
| CHAAL00000005965 | FAM69A | 0.194414608 | Autosome |
| CHAAL00000010806 | ELMO2 | 0.194414608 | Autosome |
| CHAAL00000005963 | EVI5 | 0.194331984 | Autosome |
| CHAAL00000010371 | PTPRU | 0.194331984 | Autosome |
| CHAAL00000014032 | RNF215 | 0.194331984 | Autosome |
| CHAAL00000001609 | CALCOCO2 | 0.194248508 | Autosome |
| CHAAL00000001558 | KMT2E | 0.194224499 | Autosome |
| CHAAL00000015470 | MTG2 | 0.194192377 | Autosome |
| CHAAL00000008751 | RAD50 | 0.194132873 | Autosome |
| CHAAL00000009628 | SLC9A4 | 0.193963931 | Autosome |
| CHAAL00000006631 | | 0.193919264 | Autosome |
| CHAAL00000005954 | TGFBR3 | 0.19375645 | Autosome |
| CHAAL00000000787 | PLCD1 | 0.193684211 | Autosome |
| CHAAL00000011493 | SLC39A6 | 0.193666618 | Autosome |
| CHAAL00000012072 | LOC101871754 | 0.19365722 | Autosome |

| | | | |
|---|---|---|---|
| CHAAL00000014807 | LOC104293019 | 0.193600387 | Autosome |
| CHAAL00000014220 | ARHGAP31 | 0.193340494 | Autosome |
| CHAAL00000011419 | ZFAT | 0.193322289 | Autosome |
| CHAAL00000009172 | PPP4R1 | 0.192982456 | Autosome |
| CHAAL00000003955 | | 0.192309848 | Autosome |
| CHAAL00000010680 | PEX13 | 0.192307692 | Autosome |
| CHAAL00000012488 | CACNG1 | 0.19221968 | Autosome |
| CHAAL00000013666 | LOC104373218 | 0.192166463 | Autosome |
| CHAAL00000001578 | CREG2 | 0.191729323 | Autosome |
| CHAAL00000013236 | RAD51D | 0.191456903 | Autosome |
| CHAAL00000015346 | GJD4 | 0.191333276 | Autosome |
| CHAAL00000000510 | PGR | 0.190918473 | Autosome |
| CHAAL00000008345 | | 0.190887667 | Autosome |
| CHAAL00000001344 | PPP2R2C | 0.190665343 | Autosome |
| CHAAL00000001707 | COMMD8 | 0.190665343 | Autosome |
| CHAAL00000002632 | LPCAT3 | 0.190665343 | Autosome |
| CHAAL00000009587 | TPR | 0.190665343 | Autosome |
| CHAAL00000010666 | LOC106902692 | 0.190665343 | Autosome |
| CHAAL00000014886 | TSPAN13 | 0.190665343 | Autosome |
| CHAAL00000000721 | CHM | 0.190407211 | Autosome |
| CHAAL00000007903 | SPACA9 | 0.190283401 | Autosome |
| CHAAL00000012760 | RAPH1 | 0.190283401 | Autosome |
| CHAAL00000010768 | RIPOR3 | 0.190003637 | Autosome |
| CHAAL00000007586 | | 0.189967105 | Autosome |
| CHAAL00000005390 | LOC106895107 | 0.189841416 | Autosome |
| CHAAL00000013652 | RAB3IL1 | 0.189566837 | Autosome |
| CHAAL00000008496 | ERICH6B | 0.189473684 | Autosome |
| CHAAL00000012057 | CCDC103 | 0.18939685 | Autosome |
| CHAAL00000015605 | PDIA3 | 0.189379699 | Autosome |
| CHAAL00000001441 | IL17RD | 0.189332391 | Autosome |
| CHAAL00000011572 | HARBI1 | 0.1888969 | Autosome |
| CHAAL00000000495 | RARS | 0.18883523 | Autosome |
| CHAAL00000001531 | LOC104287262 | 0.188822572 | Autosome |
| CHAAL00000002982 | PPP3CB | 0.188736172 | Autosome |
| CHAAL00000000398 | | 0.188572098 | Autosome |
| CHAAL00000002249 | | 0.18839154 | Autosome |
| CHAAL00000015173 | TBP | 0.188332276 | Autosome |
| CHAAL00000006050 | MPG | 0.188304094 | Autosome |
| CHAAL00000014931 | MOGAT2 | 0.188224799 | Autosome |
| CHAAL00000005833 | MCMDC2 | 0.188191882 | Autosome |
| CHAAL00000006672 | MICU1 | 0.188034188 | Autosome |
| CHAAL00000002453 | ABCA3 | 0.187825795 | Autosome |
| CHAAL00000005907 | FUBP1 | 0.18767981 | Autosome |

| | | | |
|---|---|---|---|
| CHAAL00000008366 | RPRD2 | 0.187643021 | Autosome |
| CHAAL00000010545 | Sep-03 | 0.187643021 | Autosome |
| CHAAL00000002322 | LOC104016420, LOC104643102 | 0.187566231 | Autosome |
| CHAAL00000011065 | LOC104835929 | 0.1875 | Autosome |
| CHAAL00000003192 | LOC104294485 | 0.187134503 | Autosome |
| CHAAL00000003377 | LOC101878579 | 0.187134503 | Autosome |
| CHAAL00000008152 | PLIN1 | 0.187134503 | Autosome |
| CHAAL00000007607 | LOC103912354 | 0.186602871 | Autosome |
| CHAAL00000000250 | SV2B | 0.186351706 | Autosome |
| CHAAL00000004733 | GPR6 | 0.1860085 | Autosome |
| CHAAL00000001295 | RIC8A | 0.185805423 | Autosome |
| CHAAL00000012608 | OXTR | 0.185555129 | Autosome |
| CHAAL00000009435 | LOC104291175 | 0.185463659 | Autosome |
| CHAAL00000011439 | | 0.185463659 | Autosome |
| CHAAL00000013389 | | 0.185463659 | Autosome |
| CHAAL00000004225 | RAD54L2 | 0.185309712 | Autosome |
| CHAAL00000010634 | LOC104158658 | 0.18494152 | Autosome |
| CHAAL00000003932 | KDELC2 | 0.184373978 | Autosome |
| CHAAL00000009446 | CRISPLD2 | 0.18431972 | Autosome |
| CHAAL00000000698 | LOC104283564 | 0.184210526 | Autosome |
| CHAAL00000002959 | NEK3 | 0.184210526 | Autosome |
| CHAAL00000004303 | PLPPR3 | 0.184210526 | Autosome |
| CHAAL00000013746 | | 0.184210526 | Autosome |
| CHAAL00000005545 | INTS11 | 0.184117318 | Autosome |

**Supplementary Table 19** High *Tajima's D* genes and gene descriptions for Kentish plover. Genes were considered to have high *Tajima's D* if they fell above the 95th percentile for positive genes (autosomes: *Tajima's D* > 1.565, Z: *Tajima's D* > 1.723). Gene symbols were assigned based on best reciprocal BLAST matches from the RefSeq protein database.

| Gene Name | Gene Symbol | *Tajima's D* | Chromosome |
|---|---|---|---|
| CHAAL00000001241 | | 3.02611691 | Autosome |
| CHAAL00000001242 | LOC106019565 | 3.02611691 | Autosome |
| CHAAL00000004928 | SERINC1 | 2.69431138 | Autosome |
| CHAAL00000003535 | | 2.61805574 | Z |
| CHAAL00000006800 | | 2.58422841 | Autosome |
| CHAAL00000007141 | CACNA1S | 2.55318369 | Autosome |
| CHAAL00000004454 | NUP35 | 2.51155149 | Autosome |
| CHAAL00000014607 | | 2.50486989 | Z |
| CHAAL00000003441 | | 2.44299688 | Z |

| | | | |
|---|---|---|---|
| CHAAL00000006302 | LOC104307831 | 2.43977177 | Autosome |
| CHAAL00000014877 | | 2.42764346 | Autosome |
| CHAAL00000011683 | CER1 | 2.42653515 | Z |
| CHAAL00000013967 | | 2.38848486 | Z |
| CHAAL00000012084 | CCT5 | 2.37542832 | Autosome |
| CHAAL00000002551 | PARG | 2.35711168 | Autosome |
| CHAAL00000005586 | GPR132 | 2.35357065 | Autosome |
| CHAAL00000004939 | LOC104640764 | 2.33412892 | Autosome |
| CHAAL00000008745 | GDF9 | 2.32595519 | Autosome |
| CHAAL00000014606 | LOC104272773 | 2.30298568 | Z |
| CHAAL00000008587 | | 2.27650121 | Autosome |
| CHAAL00000002766 | | 2.22867856 | Autosome |
| CHAAL00000015071 | | 2.22131404 | Autosome |
| CHAAL00000011113 | CAND1 | 2.21651314 | Autosome |
| CHAAL00000003123 | LOC104283033 | 2.21326376 | Autosome |
| CHAAL00000004551 | NCR3LG1 | 2.21326376 | Autosome |
| CHAAL00000003493 | LOC103529205 | 2.20168181 | Autosome |
| CHAAL00000006280 | FBXW11 | 2.18326569 | Autosome |
| CHAAL00000003440 | | 2.16664197 | Z |
| CHAAL00000014581 | | 2.15794845 | Z |
| CHAAL00000003902 | GRK6 | 2.15001824 | Autosome |
| CHAAL00000011405 | | 2.14045125 | Autosome |
| CHAAL00000002369 | | 2.12900599 | Autosome |
| CHAAL00000004833 | LOC104285445 | 2.11851032 | Autosome |
| CHAAL00000000479 | C1QTNF2 | 2.11652531 | Autosome |
| CHAAL00000011929 | TOGARAM2 | 2.11041019 | Autosome |
| CHAAL00000006143 | SENP7 | 2.0965062 | Autosome |
| CHAAL00000006758 | GFRAL | 2.09164034 | Autosome |
| CHAAL00000007032 | MTREX | 2.08352334 | Z |
| CHAAL00000005443 | DNAJC1 | 2.08352334 | Autosome |
| CHAAL00000010401 | MED8 | 2.08115314 | Autosome |
| CHAAL00000003647 | DDR2 | 2.0794178 | Autosome |
| CHAAL00000009789 | AHCTF1 | 2.06859784 | Autosome |
| CHAAL00000001524 | GTSE1 | 2.06542009 | Autosome |
| CHAAL00000008494 | SPERT | 2.06516339 | Autosome |
| CHAAL00000007746 | VEZT | 2.05755357 | Autosome |
| CHAAL00000015209 | ADAD1 | 2.04966488 | Autosome |
| CHAAL00000002610 | | 2.04796422 | Autosome |
| CHAAL00000014213 | RASSF5 | 2.03365216 | Autosome |
| CHAAL00000014690 | AP3M2 | 2.03365216 | Autosome |
| CHAAL00000014242 | FSTL1 | 2.03362146 | Autosome |
| CHAAL00000007506 | BID | 2.03336279 | Autosome |
| CHAAL00000011320 | EZR | 2.03336279 | Autosome |

| | | | |
|---|---|---|---|
| CHAAL00000008629 | ENTPD2 | 2.02511052 | Autosome |
| CHAAL00000011562 | CRY2 | 2.02511052 | Autosome |
| CHAAL00000008543 | NUP62 | 2.0174507 | Autosome |
| CHAAL00000001999 | BCLAF3 | 2.01147268 | Autosome |
| CHAAL00000003999 | RPS10 | 2.01147268 | Autosome |
| CHAAL00000009657 | SKA2 | 2.01147268 | Autosome |
| CHAAL00000012148 | | 2.01147268 | Autosome |
| CHAAL00000008016 | | 2.00097358 | Autosome |
| CHAAL00000006163 | | 2.00007669 | Autosome |
| CHAAL00000001752 | OTULINL | 1.99704434 | Autosome |
| CHAAL00000002458 | RIPK4 | 1.99082997 | Autosome |
| CHAAL00000008223 | | 1.98958257 | Z |
| CHAAL00000004941 | KPNA5 | 1.98958257 | Autosome |
| CHAAL00000007275 | LEKR1 | 1.98958257 | Autosome |
| CHAAL00000010549 | SLC16A3 | 1.98958257 | Autosome |
| CHAAL00000012908 | UPRT | 1.98958257 | Autosome |
| CHAAL00000014789 | | 1.98958257 | Autosome |
| CHAAL00000015531 | TSPAN5 | 1.98958257 | Autosome |
| CHAAL00000002026 | ASB11 | 1.98675486 | Autosome |
| CHAAL00000007198 | LOC104286641 | 1.98606298 | Autosome |
| CHAAL00000013165 | MAP3K1 | 1.98378099 | Z |
| CHAAL00000013364 | IFT22 | 1.98378099 | Autosome |
| CHAAL00000002192 | ANKRD13A | 1.98310711 | Autosome |
| CHAAL00000008248 | LOC104296055 | 1.97940312 | Z |
| CHAAL00000001690 | EIF6 | 1.97940312 | Autosome |
| CHAAL00000013382 | | 1.97102182 | Autosome |
| CHAAL00000012505 | KCNJ16 | 1.96797627 | Autosome |
| CHAAL00000006149 | TMEM45A | 1.96715726 | Autosome |
| CHAAL00000000806 | NCR3LG1 | 1.95787672 | Autosome |
| CHAAL00000014584 | LOC104287688 | 1.9565032 | Z |
| CHAAL00000002016 | | 1.95528877 | Autosome |
| CHAAL00000004328 | NCLN | 1.94580234 | Autosome |
| CHAAL00000005527 | LOC104008945 | 1.94580234 | Autosome |
| CHAAL00000004277 | AAMDC | 1.94512257 | Autosome |
| CHAAL00000003840 | TLR3 | 1.94220741 | Autosome |
| CHAAL00000000187 | AGTPBP1 | 1.92391222 | Z |
| CHAAL00000001585 | FER | 1.92391222 | Z |
| CHAAL00000001484 | SYCP2L | 1.92391222 | Autosome |
| CHAAL00000004146 | LOC101872941 | 1.92391222 | Autosome |
| CHAAL00000004253 | LOC101868153 | 1.92391222 | Autosome |
| CHAAL00000005943 | LOC101869757 | 1.92391222 | Autosome |
| CHAAL00000007325 | LMNB2 | 1.92391222 | Autosome |
| CHAAL00000010706 | ISM1 | 1.92391222 | Autosome |

| | | | |
|---|---|---|---|
| CHAAL00000010778 | | 1.92391222 | Autosome |
| CHAAL00000014224 | TMEM39A | 1.92391222 | Autosome |
| CHAAL00000014453 | HOMER2 | 1.92391222 | Autosome |
| CHAAL00000011336 | | 1.92382268 | Autosome |
| CHAAL00000001767 | ZNF536 | 1.92073558 | Autosome |
| CHAAL00000003646 | TOR4A | 1.91595615 | Autosome |
| CHAAL00000000508 | LOC101872127 | 1.90066236 | Autosome |
| CHAAL00000007548 | EFHC1 | 1.89786917 | Autosome |
| CHAAL00000002958 | CKAP2 | 1.89575797 | Autosome |
| CHAAL00000007576 | STRAP | 1.88893463 | Autosome |
| CHAAL00000000533 | HHATL | 1.88798832 | Autosome |
| CHAAL00000001083 | RASL11A | 1.88403863 | Autosome |
| CHAAL00000008959 | GALR2 | 1.88403863 | Autosome |
| CHAAL00000015183 | THBS2 | 1.87866549 | Autosome |
| CHAAL00000001686 | CEP250 | 1.86942496 | Autosome |
| CHAAL00000001414 | | 1.85888408 | Z |
| CHAAL00000001860 | PLA2G12A | 1.85824188 | Autosome |
| CHAAL00000002893 | KIAA1147 | 1.85824188 | Autosome |
| CHAAL00000004493 | | 1.85824188 | Autosome |
| CHAAL00000005231 | | 1.85824188 | Autosome |
| CHAAL00000012397 | TXNDC17 | 1.85824188 | Autosome |
| CHAAL00000011241 | TTC4 | 1.85506057 | Autosome |
| CHAAL00000014978 | LOC104033167 | 1.85479921 | Autosome |
| CHAAL00000002919 | PARP12 | 1.85131211 | Autosome |
| CHAAL00000010881 | OGFOD2 | 1.8479258 | Autosome |
| CHAAL00000002272 | LOC104252744 | 1.83635177 | Autosome |
| CHAAL00000006611 | ZFAND2B | 1.83635177 | Autosome |
| CHAAL00000009160 | TUBB6 | 1.83635177 | Autosome |
| CHAAL00000009234 | FOSL2 | 1.83635177 | Autosome |
| CHAAL00000013506 | SH3GL1 | 1.83635177 | Autosome |
| CHAAL00000003110 | LOC104293975 | 1.83487978 | Autosome |
| CHAAL00000003388 | GPD1 | 1.83416746 | Autosome |
| CHAAL00000011439 | | 1.83416746 | Autosome |
| CHAAL00000011473 | CARMIL1 | 1.83398021 | Autosome |
| CHAAL00000005871 | | 1.8228089 | Autosome |
| CHAAL00000003546 | HMGCR | 1.8215592 | Z |
| CHAAL00000003321 | LOC104693568 | 1.8215592 | Autosome |
| CHAAL00000002645 | LOC104293427 | 1.82136607 | Autosome |
| CHAAL00000010873 | ARPC3 | 1.82136607 | Autosome |
| CHAAL00000005964 | RPL5 | 1.81754373 | Autosome |
| CHAAL00000000165 | PSAT1 | 1.81446165 | Z |
| CHAAL00000002662 | VAMP1 | 1.81446165 | Autosome |
| CHAAL00000003390 | LIMA1 | 1.81446165 | Autosome |

| | | | |
|---|---|---|---|
| CHAAL00000012408 | DHRS11 | 1.81446165 | Autosome |
| CHAAL00000004361 | | 1.80732929 | Autosome |
| CHAAL00000015534 | ADH5 | 1.80582483 | Autosome |
| CHAAL00000010688 | CCT4 | 1.8051172 | Autosome |
| CHAAL00000007227 | MYH15 | 1.79617438 | Autosome |
| CHAAL00000002442 | | 1.79308149 | Autosome |
| CHAAL00000006781 | | 1.79122031 | Autosome |
| CHAAL00000005508 | | 1.79009178 | Autosome |
| CHAAL00000005862 | MTMR1 | 1.78514667 | Autosome |
| CHAAL00000006351 | BRAP | 1.78514667 | Autosome |
| CHAAL00000007828 | ST6GALNAC4 | 1.78429628 | Autosome |
| CHAAL00000004221 | DOCK3 | 1.78370928 | Autosome |
| CHAAL00000004354 | FGF14 | 1.77966129 | Autosome |
| CHAAL00000007549 | PAQR8 | 1.77197101 | Autosome |
| CHAAL00000006929 | BTBD10 | 1.76767256 | Autosome |
| CHAAL00000012795 | TRAK2 | 1.76767256 | Autosome |
| CHAAL00000004127 | AMPD1 | 1.76731121 | Autosome |
| CHAAL00000013361 | RPAIN | 1.76731121 | Autosome |
| CHAAL00000013008 | RP1 | 1.76563529 | Autosome |
| CHAAL00000010227 | LOC102043728 | 1.76229297 | Autosome |
| CHAAL00000004008 | RPL10A | 1.75104883 | Autosome |
| CHAAL00000004464 | LANCL1 | 1.75104883 | Autosome |
| CHAAL00000015173 | TBP | 1.75104883 | Autosome |
| CHAAL00000002712 | TTC29 | 1.75086612 | Autosome |
| CHAAL00000011788 | IGSF9 | 1.75086612 | Autosome |
| CHAAL00000001687 | GDF5 | 1.75075917 | Autosome |
| CHAAL00000006959 | LOC104291707 | 1.75075917 | Autosome |
| CHAAL00000000753 | VGLL1 | 1.74879131 | Autosome |
| CHAAL00000002825 | HIBADH | 1.74879131 | Autosome |
| CHAAL00000002982 | PPP3CB | 1.74879131 | Autosome |
| CHAAL00000004072 | LOC104573748 | 1.74879131 | Autosome |
| CHAAL00000004189 | LOC104286901 | 1.74879131 | Autosome |
| CHAAL00000005479 | ADAM22 | 1.74879131 | Autosome |
| CHAAL00000012796 | CASP10 | 1.74879131 | Autosome |
| CHAAL00000013256 | TPST1, LOC101870184 | 1.74879131 | Autosome |
| CHAAL00000013590 | NR5A1 | 1.74879131 | Autosome |
| CHAAL00000013845 | | 1.74879131 | Autosome |
| CHAAL00000015399 | LRRC59 | 1.74879131 | Autosome |
| CHAAL00000004772 | SLC17A5 | 1.74630607 | Autosome |
| CHAAL00000006720 | STYK1 | 1.74289264 | Autosome |
| CHAAL00000014264 | WDR3 | 1.73791942 | Autosome |
| CHAAL00000014360 | KIF17 | 1.73753566 | Autosome |
| CHAAL00000004977 | REPS1 | 1.73442511 | Autosome |

| | | | |
|---|---|---|---|
| CHAAL00000007872 | LOC104289104 | 1.72801242 | Autosome |
| CHAAL00000014195 | | 1.72677007 | Autosome |
| CHAAL00000010937 | | 1.7204478 | Autosome |
| CHAAL00000010327 | PHF13 | 1.71999485 | Autosome |
| CHAAL00000006888 | | 1.7195 | Autosome |
| CHAAL00000009771 | | 1.71721784 | Autosome |
| CHAAL00000000278 | | 1.71668137 | Autosome |
| CHAAL00000002667 | FAM162A | 1.71325636 | Autosome |
| CHAAL00000012378 | | 1.71300019 | Autosome |
| CHAAL00000006411 | ABCD4 | 1.71246517 | Autosome |
| CHAAL00000004394 | LOC101867830 | 1.71122444 | Autosome |
| CHAAL00000008370 | PTPN14 | 1.70993564 | Autosome |
| CHAAL00000005496 | PEX1 | 1.70948691 | Autosome |
| CHAAL00000012961 | LOC106023452 | 1.70912763 | Autosome |
| CHAAL00000002321 | RAD54B | 1.70735193 | Autosome |
| CHAAL00000003867 | TBC1D9 | 1.70551005 | Autosome |
| CHAAL00000012398 | KIAA0753 | 1.70551005 | Autosome |
| CHAAL00000009858 | AMFR | 1.69974265 | Autosome |
| CHAAL00000010876 | CCDC62 | 1.69974265 | Autosome |
| CHAAL00000001244 | PLA2G4F | 1.69644961 | Autosome |
| CHAAL00000002015 | SYAP1 | 1.69408354 | Autosome |
| CHAAL00000014165 | MARK3 | 1.69373187 | Autosome |
| CHAAL00000002910 | | 1.69368362 | Autosome |
| CHAAL00000015514 | GK5 | 1.6926299 | Autosome |
| CHAAL00000006796 | | 1.68779932 | Autosome |
| CHAAL00000012221 | | 1.68455393 | Autosome |
| CHAAL00000000754 | HTATSF1 | 1.68312097 | Autosome |
| CHAAL00000002346 | | 1.68312097 | Autosome |
| CHAAL00000005027 | CLSTN2 | 1.68271226 | Autosome |
| CHAAL00000010057 | RUNDC1 | 1.68271226 | Autosome |
| CHAAL00000004355 | | 1.68271052 | Autosome |
| CHAAL00000006306 | LOC103526522 | 1.6761428 | Autosome |
| CHAAL00000007060 | SLC24A3 | 1.67087817 | Autosome |
| CHAAL00000004077 | LOC104310750 | 1.67087671 | Autosome |
| CHAAL00000000209 | DPP8 | 1.66955663 | Autosome |
| CHAAL00000007191 | LOC104286660 | 1.66422741 | Autosome |
| CHAAL00000012563 | LOC105400195 | 1.66422741 | Autosome |
| CHAAL00000008590 | | 1.66342127 | Autosome |
| CHAAL00000007930 | MRPS2 | 1.66287698 | Autosome |
| CHAAL00000002044 | HAUS2 | 1.66123085 | Autosome |
| CHAAL00000002501 | KCNE2 | 1.66123085 | Autosome |
| CHAAL00000002868 | IPO8 | 1.66123085 | Autosome |
| CHAAL00000003141 | MAML2 | 1.66123085 | Autosome |

| | | | |
|---|---|---|---|
| CHAAL00000007562 | LOC104281720 | 1.66123085 | Autosome |
| CHAAL00000007770 | LOC104284336 | 1.66123085 | Autosome |
| CHAAL00000010270 | TMOD2 | 1.66123085 | Autosome |
| CHAAL00000011526 | PPCS | 1.66123085 | Autosome |
| CHAAL00000013359 | DHX33 | 1.65945132 | Autosome |
| CHAAL00000005589 | CLBA1 | 1.65920151 | Autosome |
| CHAAL00000007359 | CFAP58 | 1.65920151 | Autosome |
| CHAAL00000010302 | ATRIP | 1.65622961 | Autosome |
| CHAAL00000007131 | KDM5B | 1.65528709 | Autosome |
| CHAAL00000013479 | CAPN8 | 1.65523912 | Autosome |
| CHAAL00000006255 | DECR1 | 1.65490609 | Autosome |
| CHAAL00000000292 | NT5C1B | 1.65130648 | Autosome |
| CHAAL00000011410 | COL22A1 | 1.65130648 | Autosome |
| CHAAL00000002155 | TRIP11 | 1.65085311 | Autosome |
| CHAAL00000003281 | FGA | 1.64849436 | Autosome |
| CHAAL00000008443 | LOC106890872 | 1.64849436 | Autosome |
| CHAAL00000009143 | COL4A1 | 1.64815233 | Autosome |
| CHAAL00000015535 | TEKT3 | 1.64551062 | Autosome |
| CHAAL00000006908 | | 1.6430417 | Autosome |
| CHAAL00000011089 | INCENP | 1.64184694 | Autosome |
| CHAAL00000014702 | MTERF2 | 1.64106117 | Autosome |
| CHAAL00000001997 | LOC103540408 | 1.63934074 | Autosome |
| CHAAL00000002438 | | 1.63934074 | Autosome |
| CHAAL00000002923 | SYCP3 | 1.63934074 | Autosome |
| CHAAL00000003359 | | 1.63934074 | Autosome |
| CHAAL00000004906 | TMEM106B | 1.63934074 | Autosome |
| CHAAL00000008690 | ECHDC3 | 1.63934074 | Autosome |
| CHAAL00000009049 | KCTD5 | 1.63934074 | Autosome |
| CHAAL00000009722 | LOC104283176 | 1.63934074 | Autosome |
| CHAAL00000012140 | MSRB1 | 1.63934074 | Autosome |
| CHAAL00000013010 | LOC104019791 | 1.63934074 | Autosome |
| CHAAL00000006448 | EAPP | 1.63468275 | Autosome |
| CHAAL00000007449 | CPN1 | 1.63468275 | Autosome |
| CHAAL00000008355 | AGAP3 | 1.63468275 | Autosome |
| CHAAL00000014853 | CD83 | 1.63468275 | Autosome |
| CHAAL00000008500 | GTF2F2 | 1.63217408 | Autosome |
| CHAAL00000003392 | DIP2B | 1.62963797 | Autosome |
| CHAAL00000012079 | MTRR | 1.62693541 | Autosome |
| CHAAL00000010753 | CSE1L | 1.62517078 | Autosome |
| CHAAL00000003158 | PTPN21 | 1.62352035 | Autosome |
| CHAAL00000009056 | ANKS4B | 1.62320643 | Autosome |
| CHAAL00000009868 | MBTPS1 | 1.62108198 | Autosome |
| CHAAL00000000846 | MRPL45 | 1.61866037 | Autosome |

| | | | |
|---|---|---|---|
| CHAAL00000005537 | MRPL20 | 1.61866037 | Autosome |
| CHAAL00000009582 | OPCML | 1.61866037 | Autosome |
| CHAAL00000011085 | PDGFD | 1.61866037 | Autosome |
| CHAAL00000000581 | | 1.61805903 | Autosome |
| CHAAL00000006725 | ZNF512 | 1.61805903 | Autosome |
| CHAAL00000003925 | ALKBH8 | 1.61374393 | Autosome |
| CHAAL00000008340 | CHODL | 1.61374393 | Autosome |
| CHAAL00000000545 | ZDHHC18 | 1.61328879 | Autosome |
| CHAAL00000007233 | LOC104077359 | 1.61328879 | Autosome |
| CHAAL00000015379 | PAXIP1 | 1.61328879 | Autosome |
| CHAAL00000002838 | ANKRD28 | 1.60916175 | Autosome |
| CHAAL00000008015 | | 1.60740595 | Autosome |
| CHAAL00000007193 | LOC104286645 | 1.60534372 | Autosome |
| CHAAL00000011855 | FAM221A | 1.60514665 | Autosome |
| CHAAL00000000916 | CCM2 | 1.60266949 | Autosome |
| CHAAL00000004821 | LOC104290067 | 1.60231708 | Autosome |
| CHAAL00000011643 | CPNE2 | 1.60231708 | Autosome |
| CHAAL00000005248 | POSTN | 1.59720912 | Autosome |
| CHAAL00000002462 | MNT | 1.59556051 | Autosome |
| CHAAL00000003462 | LPXN | 1.59556051 | Autosome |
| CHAAL00000008105 | LOC104020516 | 1.59556051 | Autosome |
| CHAAL00000009169 | LOC106630465 | 1.59345351 | Autosome |
| CHAAL00000003119 | CEP295 | 1.593167 | Autosome |
| CHAAL00000000395 | LOC104281694 | 1.59299273 | Autosome |
| CHAAL00000000804 | NGLY1 | 1.59163294 | Autosome |
| CHAAL00000005906 | NEXN | 1.59089023 | Autosome |
| CHAAL00000007672 | LOC104282654 | 1.59089023 | Autosome |
| CHAAL00000011438 | NELL1 | 1.59089023 | Autosome |
| CHAAL00000014094 | AIFM2 | 1.59089023 | Autosome |
| CHAAL00000001717 | AP5Z1 | 1.58843871 | Autosome |
| CHAAL00000012926 | ACSL4 | 1.58795686 | Autosome |
| CHAAL00000002246 | MCF2L | 1.58481158 | Autosome |
| CHAAL00000004080 | LOC103539723 | 1.58481158 | Autosome |
| CHAAL00000001020 | SLC35G2 | 1.58353587 | Autosome |
| CHAAL00000001441 | IL17RD | 1.58353587 | Autosome |
| CHAAL00000008012 | UROC1 | 1.58274672 | Autosome |
| CHAAL00000003834 | GP9 | 1.57946338 | Autosome |
| CHAAL00000015397 | ACSF2 | 1.57946338 | Autosome |
| CHAAL00000001539 | CBLL1 | 1.57811923 | Autosome |
| CHAAL00000010926 | CHFR | 1.57811923 | Autosome |
| CHAAL00000010768 | RIPOR3 | 1.57811175 | Autosome |
| CHAAL00000004519 | EXOC4 | 1.57361824 | Autosome |
| CHAAL00000007210 | RIPK1 | 1.57361824 | Autosome |

| | | | |
|---|---|---|---|
| CHAAL00000015668 | | 1.5708979 | Autosome |
| CHAAL00000001417 | FDFT1 | 1.56818785 | Autosome |
| CHAAL00000006456 | DTD2 | 1.56818785 | Autosome |
| CHAAL00000008841 | SLC36A1 | 1.56818785 | Autosome |
| CHAAL00000006342 | DNAJB13 | 1.56803653 | Autosome |
| CHAAL00000011459 | ZNF407 | 1.56759703 | Autosome |
| CHAAL00000000082 | LMO1 | 1.56502749 | Autosome |
| CHAAL00000000291 | RDH14 | 1.56502749 | Autosome |
| CHAAL00000000730 | DIAPH1 | 1.56502749 | Autosome |
| CHAAL00000001017 | PCCB | 1.56502749 | Autosome |
| CHAAL00000001236 | NMRAL1 | 1.56502749 | Autosome |
| CHAAL00000001628 | TMPPE | 1.56502749 | Autosome |
| CHAAL00000001750 | ANKH | 1.56502749 | Autosome |
| CHAAL00000002907 | LOC104288579 | 1.56502749 | Autosome |
| CHAAL00000003115 | MTNR1B | 1.56502749 | Autosome |
| CHAAL00000003795 | | 1.56502749 | Autosome |
| CHAAL00000004124 | LOC101881780 | 1.56502749 | Autosome |
| CHAAL00000004611 | OTOP2 | 1.56502749 | Autosome |
| CHAAL00000005429 | GPR158 | 1.56502749 | Autosome |
| CHAAL00000005658 | ITPA | 1.56502749 | Autosome |
| CHAAL00000005809 | KXD1 | 1.56502749 | Autosome |
| CHAAL00000006504 | LOC104285104 | 1.56502749 | Autosome |
| CHAAL00000006511 | | 1.56502749 | Autosome |
| CHAAL00000007070 | RHOA | 1.56502749 | Autosome |
| CHAAL00000008011 | SNX16 | 1.56502749 | Autosome |
| CHAAL00000008103 | LOC104832078 | 1.56502749 | Autosome |
| CHAAL00000008158 | BTLA | 1.56502749 | Autosome |
| CHAAL00000008646 | AMBP | 1.56502749 | Autosome |
| CHAAL00000009161 | | 1.56502749 | Autosome |
| CHAAL00000009282 | DRGX | 1.56502749 | Autosome |
| CHAAL00000009429 | CMIP | 1.56502749 | Autosome |
| CHAAL00000009739 | | 1.56502749 | Autosome |
| CHAAL00000010236 | | 1.56502749 | Autosome |
| CHAAL00000010409 | EBNA1BP2 | 1.56502749 | Autosome |
| CHAAL00000011192 | CHMP2B | 1.56502749 | Autosome |
| CHAAL00000011899 | NUCKS1 | 1.56502749 | Autosome |
| CHAAL00000012074 | | 1.56502749 | Autosome |
| CHAAL00000012353 | LOC104124394 | 1.56502749 | Autosome |
| CHAAL00000012598 | RHO | 1.56502749 | Autosome |
| CHAAL00000012755 | LOC104291063 | 1.56502749 | Autosome |
| CHAAL00000012761 | CD28 | 1.56502749 | Autosome |
| CHAAL00000012853 | GPR180 | 1.56502749 | Autosome |
| CHAAL00000013367 | LOC104027930 | 1.56502749 | Autosome |

| | | | |
|---|---|---|---|
| CHAAL00000013368 | CUX1 | 1.56502749 | Autosome |
| CHAAL00000013729 | LOC101875497 | 1.56502749 | Autosome |
| CHAAL00000014138 | DCUN1D4 | 1.56502749 | Autosome |
| CHAAL00000014773 | CSNK1E | 1.56502749 | Autosome |

**Supplementary Table 20** High *Tajima's D* genes and gene descriptions for white-faced plover. Genes were considered to have high *Tajima's D* if they fell above the 95th percentile for positive genes (autosomes: *Tajima's D* ≥ 1.669, Z: *Tajima's D* ≥ 1.835). Gene symbols were assigned based on best reciprocal BLAST matches from the RefSeq protein database.

| Gene Name | Gene Symbol | *Tajima's D* | Chromosome |
|---|---|---|---|
| CHAAL00000001241 | | 3.01958733 | Autosome |
| CHAAL00000001242 | LOC106019565 | 3.01958733 | Autosome |
| CHAAL00000015182 | | 2.94306018 | Autosome |
| CHAAL00000014607 | | 2.65962448 | Z |
| CHAAL00000003535 | | 2.61317338 | Z |
| CHAAL00000006800 | | 2.53307169 | Autosome |
| CHAAL00000013967 | | 2.51192775 | Z |
| CHAAL00000014606 | LOC104272773 | 2.46770478 | Z |
| CHAAL00000007548 | EFHC1 | 2.45804386 | Autosome |
| CHAAL00000005575 | LOC104327277 | 2.45087973 | Autosome |
| CHAAL00000010929 | ANKLE2 | 2.44790396 | Autosome |
| CHAAL00000000114 | PTPN5 | 2.43647711 | Autosome |
| CHAAL00000015181 | WDR27 | 2.40544372 | Autosome |
| CHAAL00000004770 | | 2.37312718 | Autosome |
| CHAAL00000012149 | LOC105403338 | 2.34719404 | Autosome |
| CHAAL00000001686 | CEP250 | 2.34189646 | Autosome |
| CHAAL00000012613 | TMEM161B | 2.34128405 | Z |
| CHAAL00000008246 | TOPORS | 2.33488718 | Z |
| CHAAL00000003440 | | 2.33363546 | Z |
| CHAAL00000001563 | SLC26A5 | 2.32735876 | Autosome |
| CHAAL00000005027 | CLSTN2 | 2.32137346 | Autosome |
| CHAAL00000003493 | LOC103529205 | 2.30577963 | Autosome |
| CHAAL00000001977 | LOC104380646 | 2.29963177 | Autosome |
| CHAAL00000005393 | DHRS9 | 2.29935491 | Autosome |
| CHAAL00000014278 | ERCC6L2 | 2.28300804 | Z |
| CHAAL00000014597 | RIT2 | 2.26731861 | Z |
| CHAAL00000014405 | IWS1 | 2.26731861 | Autosome |
| CHAAL00000010149 | CYP1A5 | 2.26638432 | Autosome |
| CHAAL00000012614 | | 2.26164301 | Z |
| CHAAL00000010787 | LOC104293944 | 2.25498979 | Autosome |

| | | | |
|---|---|---|---|
| CHAAL00000012227 | SNTB1 | 2.24976059 | Autosome |
| CHAAL00000007198 | LOC104286641 | 2.23866412 | Autosome |
| CHAAL00000006467 | | 2.23061708 | Autosome |
| CHAAL00000004077 | LOC104310750 | 2.22085599 | Autosome |
| CHAAL00000014264 | WDR3 | 2.21682457 | Autosome |
| CHAAL00000003089 | LOC104293991 | 2.21651314 | Autosome |
| CHAAL00000006773 | LOC102049827 | 2.20279499 | Autosome |
| CHAAL00000005577 | | 2.19359693 | Autosome |
| CHAAL00000005578 | LOC104072859 | 2.19359693 | Autosome |
| CHAAL00000009306 | | 2.18380169 | Autosome |
| CHAAL00000008035 | GPR78 | 2.18326569 | Autosome |
| CHAAL00000000268 | LOC104613886 | 2.17272262 | Autosome |
| CHAAL00000009794 | QPCT | 2.17272262 | Autosome |
| CHAAL00000003397 | | 2.16664197 | Autosome |
| CHAAL00000000112 | SPTY2D1 | 2.16223271 | Autosome |
| CHAAL00000001840 | LOC104282992 | 2.16185555 | Autosome |
| CHAAL00000004603 | SLC16A5 | 2.15875891 | Autosome |
| CHAAL00000006302 | LOC104307831 | 2.15359274 | Autosome |
| CHAAL00000009570 | ATP12A | 2.15001824 | Autosome |
| CHAAL00000013381 | | 2.14552164 | Autosome |
| CHAAL00000007153 | Mar-11 | 2.13657132 | Autosome |
| CHAAL00000007154 | | 2.13657132 | Autosome |
| CHAAL00000001570 | FAM185A | 2.13218148 | Autosome |
| CHAAL00000012584 | USP19 | 2.13218148 | Autosome |
| CHAAL00000015071 | | 2.11287609 | Autosome |
| CHAAL00000001572 | | 2.10767752 | Autosome |
| CHAAL00000003053 | ZBTB46 | 2.10515405 | Autosome |
| CHAAL00000004480 | LOC104288717 | 2.10515405 | Autosome |
| CHAAL00000010514 | LOC103908400 | 2.10515405 | Autosome |
| CHAAL00000006908 | | 2.10509846 | Autosome |
| CHAAL00000015209 | ADAD1 | 2.10509846 | Autosome |
| CHAAL00000013811 | SAMD7 | 2.09712243 | Autosome |
| CHAAL00000011477 | ALG2 | 2.09367161 | Autosome |
| CHAAL00000002223 | PTAFR | 2.08352334 | Autosome |
| CHAAL00000007230 | LOC104289547 | 2.08352334 | Autosome |
| CHAAL00000006921 | SPON1 | 2.08224476 | Autosome |
| CHAAL00000002442 | | 2.07554392 | Autosome |
| CHAAL00000008587 | | 2.07081791 | Autosome |
| CHAAL00000015451 | NELFCD | 2.07081791 | Autosome |
| CHAAL00000008745 | GDF9 | 2.06910358 | Autosome |
| CHAAL00000001363 | ZNF518B | 2.0645595 | Autosome |
| CHAAL00000014739 | C1QTNF6 | 2.06204079 | Autosome |
| CHAAL00000002617 | RAP1GAP, LOC106891180 | 2.0510992 | Autosome |

| | | | |
|---|---|---|---|
| CHAAL00000014690 | AP3M2 | 2.0510992 | Autosome |
| CHAAL00000015017 | TES | 2.05027589 | Autosome |
| CHAAL00000008013 | ZXDC | 2.04968705 | Autosome |
| CHAAL00000006418 | FAM161B | 2.04966488 | Autosome |
| CHAAL00000001409 | DMXL1 | 2.04837322 | Z |
| CHAAL00000001690 | EIF6 | 2.04796422 | Autosome |
| CHAAL00000010052 | BRCA1 | 2.04644908 | Autosome |
| CHAAL00000001431 | CHDH | 2.03758548 | Autosome |
| CHAAL00000005884 | RGCC | 2.03758548 | Autosome |
| CHAAL00000015379 | PAXIP1 | 2.03758548 | Autosome |
| CHAAL00000003197 | LONP2 | 2.03653737 | Autosome |
| CHAAL00000007860 | NUP188 | 2.03653737 | Autosome |
| CHAAL00000013921 | COG1 | 2.0360288 | Autosome |
| CHAAL00000000372 | LOC101879987 | 2.03365216 | Autosome |
| CHAAL00000006909 | EIF3M | 2.03365216 | Autosome |
| CHAAL00000007429 | GOLGA7B | 2.03365216 | Autosome |
| CHAAL00000012016 | ACTR5 | 2.03365216 | Autosome |
| CHAAL00000001294 | SIRT3 | 2.03336279 | Autosome |
| CHAAL00000007570 | | 2.03336279 | Autosome |
| CHAAL00000010861 | RHOF | 2.03336279 | Autosome |
| CHAAL00000002988 | CHCHD1 | 2.01702844 | Autosome |
| CHAAL00000009796 | NDUFAF7 | 2.01702844 | Autosome |
| CHAAL00000001531 | LOC104287262 | 2.01147268 | Autosome |
| CHAAL00000008550 | GABRE | 2.01147268 | Autosome |
| CHAAL00000011210 | LOC101870554 | 2.01147268 | Autosome |
| CHAAL00000013484 | UPP2 | 2.01147268 | Autosome |
| CHAAL00000014821 | CCDC9B | 2.01147268 | Autosome |
| CHAAL00000005508 | | 2.00412398 | Autosome |
| CHAAL00000003091 | TPX2 | 1.99704434 | Autosome |
| CHAAL00000003525 | ZNF366 | 1.98958257 | Z |
| CHAAL00000004048 | SLC16A4 | 1.98958257 | Autosome |
| CHAAL00000009820 | | 1.98958257 | Autosome |
| CHAAL00000013854 | NUDT9 | 1.98958257 | Autosome |
| CHAAL00000008376 | GPATCH2 | 1.98378099 | Autosome |
| CHAAL00000011855 | FAM221A | 1.98353063 | Autosome |
| CHAAL00000005823 | ARMC6 | 1.98348637 | Autosome |
| CHAAL00000013595 | GOLGA1 | 1.98297459 | Autosome |
| CHAAL00000013225 | STYXL1 | 1.98024141 | Autosome |
| CHAAL00000006516 | EXD2 | 1.97350714 | Autosome |
| CHAAL00000005623 | | 1.95679589 | Autosome |
| CHAAL00000002576 | ANXA11 | 1.95654942 | Autosome |
| CHAAL00000003158 | PTPN21 | 1.95596131 | Autosome |
| CHAAL00000002854 | FARSB | 1.95494731 | Autosome |

| | | | |
|---|---|---|---|
| CHAAL00000012488 | CACNG1 | 1.95053354 | Autosome |
| CHAAL00000004909 | | 1.94976755 | Autosome |
| CHAAL00000002481 | NIM1K | 1.94580234 | Z |
| CHAAL00000002995 | ARMC4 | 1.94580234 | Autosome |
| CHAAL00000005829 | LOC104286936 | 1.94580234 | Autosome |
| CHAAL00000009160 | TUBB6 | 1.94580234 | Autosome |
| CHAAL00000014731 | NCF4 | 1.94580234 | Autosome |
| CHAAL00000015531 | TSPAN5 | 1.94580234 | Autosome |
| CHAAL00000002984 | MYOZ1 | 1.94298949 | Autosome |
| CHAAL00000009485 | SPG7 | 1.94298949 | Autosome |
| CHAAL00000001767 | ZNF536 | 1.94057086 | Autosome |
| CHAAL00000004939 | LOC104640764 | 1.93190367 | Autosome |
| CHAAL00000002958 | CKAP2 | 1.93187997 | Autosome |
| CHAAL00000011681 | TTC39B | 1.92391222 | Z |
| CHAAL00000000165 | PSAT1 | 1.92391222 | Z |
| CHAAL00000000442 | BTBD6 | 1.92391222 | Autosome |
| CHAAL00000002867 | TMTC1 | 1.92391222 | Autosome |
| CHAAL00000004483 | RAB17 | 1.92391222 | Autosome |
| CHAAL00000004638 | LOC103907554 | 1.92391222 | Autosome |
| CHAAL00000006285 | GABRP | 1.92391222 | Autosome |
| CHAAL00000008305 | BIRC5 | 1.92391222 | Autosome |
| CHAAL00000009122 | NUBP1 | 1.92391222 | Autosome |
| CHAAL00000010046 | DUSP3 | 1.92391222 | Autosome |
| CHAAL00000011110 | TMBIM4 | 1.92391222 | Autosome |
| CHAAL00000013364 | IFT22 | 1.92391222 | Autosome |
| CHAAL00000015280 | LOC104284949 | 1.92391222 | Autosome |
| CHAAL00000015549 | TACR1 | 1.92391222 | Autosome |
| CHAAL00000002910 | | 1.92226887 | Autosome |
| CHAAL00000005525 | NADK | 1.91728609 | Autosome |
| CHAAL00000007726 | UHRF1BP1L | 1.91596206 | Autosome |
| CHAAL00000006357 | | 1.91381897 | Autosome |
| CHAAL00000013859 | AFF1 | 1.91084202 | Autosome |
| CHAAL00000015371 | NOM1 | 1.90808963 | Autosome |
| CHAAL00000002327 | INTS8 | 1.90500394 | Autosome |
| CHAAL00000013068 | LOC104283236 | 1.90244835 | Autosome |
| CHAAL00000015122 | BAP1 | 1.89941517 | Autosome |
| CHAAL00000003645 | ANAPC2 | 1.89674274 | Autosome |
| CHAAL00000011351 | | 1.89235658 | Autosome |
| CHAAL00000009789 | AHCTF1 | 1.89225797 | Autosome |
| CHAAL00000002835 | PLEKHA8 | 1.88893463 | Autosome |
| CHAAL00000006955 | SCUBE2 | 1.88893463 | Autosome |
| CHAAL00000012448 | ABCA1 | 1.88798832 | Z |
| CHAAL00000015335 | ZNF438 | 1.88416985 | Autosome |

| | | | |
|---|---|---|---|
| CHAAL00000014916 | RHOBTB3 | 1.88403863 | Z |
| CHAAL00000004530 | ZDHHC6 | 1.88403863 | Autosome |
| CHAAL00000014978 | LOC104033167 | 1.88337875 | Autosome |
| CHAAL00000014048 | LOC105412744 | 1.88106503 | Autosome |
| CHAAL00000000554 | UBXN11 | 1.87114739 | Autosome |
| CHAAL00000014400 | INPP5D | 1.87114739 | Autosome |
| CHAAL00000005013 | TAAR5 | 1.8693301 | Autosome |
| CHAAL00000001491 | SLCO4C1 | 1.86741491 | Z |
| CHAAL00000002113 | GGACT | 1.86741491 | Autosome |
| CHAAL00000011916 | EHD3 | 1.86741491 | Autosome |
| CHAAL00000005017 | ENPP1 | 1.86513462 | Autosome |
| CHAAL00000006720 | STYK1 | 1.86219534 | Autosome |
| CHAAL00000003335 | LOC104146587 | 1.86190851 | Autosome |
| CHAAL00000007905 | GFI1B | 1.86190721 | Autosome |
| CHAAL00000010895 | TCTN2 | 1.86089049 | Autosome |
| CHAAL00000003524 | PTCD2 | 1.85824188 | Z |
| CHAAL00000000091 | LOC104010676 | 1.85824188 | Autosome |
| CHAAL00000003412 | CCDC65 | 1.85824188 | Autosome |
| CHAAL00000006400 | DLST | 1.85824188 | Autosome |
| CHAAL00000007959 | STOM | 1.85824188 | Autosome |
| CHAAL00000009401 | NOD1 | 1.85824188 | Autosome |
| CHAAL00000009782 | ABHD10 | 1.85824188 | Autosome |
| CHAAL00000010375 | LOC104352314 | 1.85824188 | Autosome |
| CHAAL00000015649 | SRSF6 | 1.85824188 | Autosome |
| CHAAL00000006796 | | 1.85771559 | Autosome |
| CHAAL00000014359 | RSC1A1 | 1.85771559 | Autosome |
| CHAAL00000006367 | VIPAS39 | 1.85370777 | Autosome |
| CHAAL00000009119 | LOC105407359 | 1.85370777 | Autosome |
| CHAAL00000003627 | LOC104286690 | 1.85116188 | Autosome |
| CHAAL00000001437 | ERC2 | 1.84278057 | Autosome |
| CHAAL00000003630 | SF3B6 | 1.84228092 | Autosome |
| CHAAL00000007576 | STRAP | 1.84228092 | Autosome |
| CHAAL00000008806 | RUFY1 | 1.84228092 | Autosome |
| CHAAL00000010873 | ARPC3 | 1.84228092 | Autosome |
| CHAAL00000008821 | PDGFRB | 1.84175952 | Autosome |
| CHAAL00000014877 | | 1.84079103 | Autosome |
| CHAAL00000012653 | N4BP2 | 1.83914246 | Autosome |
| CHAAL00000011698 | BUD23 | 1.83635177 | Autosome |
| CHAAL00000014224 | TMEM39A | 1.83635177 | Autosome |
| CHAAL00000003122 | TAF1D | 1.83487978 | Autosome |
| CHAAL00000004344 | PPM1H | 1.83487978 | Autosome |
| CHAAL00000008500 | GTF2F2 | 1.83416746 | Autosome |
| CHAAL00000011140 | ADRA1D | 1.83416746 | Autosome |

| | | | |
|---|---|---|---|
| CHAAL00000011353 | LOC106899559 | 1.83416746 | Autosome |
| CHAAL00000013087 | HMCES | 1.83416746 | Autosome |
| CHAAL00000001183 | THSD7B | 1.8265215 | Autosome |
| CHAAL00000007141 | CACNA1S | 1.82155787 | Autosome |
| CHAAL00000014858 | POMGNT2 | 1.82155787 | Autosome |
| CHAAL00000004362 | LOC104282449 | 1.82136607 | Autosome |
| CHAAL00000003743 | CASK | 1.81942722 | Autosome |
| CHAAL00000013023 | PENK | 1.81754373 | Autosome |
| CHAAL00000000292 | NT5C1B | 1.81446165 | Autosome |
| CHAAL00000001844 | CSGALNACT1 | 1.81446165 | Autosome |
| CHAAL00000002439 | LRRC4C | 1.81446165 | Autosome |
| CHAAL00000005015 | MOXD1 | 1.81446165 | Autosome |
| CHAAL00000006149 | TMEM45A | 1.81446165 | Autosome |
| CHAAL00000006885 | CENPQ | 1.81446165 | Autosome |
| CHAAL00000013506 | SH3GL1 | 1.81446165 | Autosome |
| CHAAL00000015259 | FAM131A | 1.81446165 | Autosome |
| CHAAL00000006127 | HHLA2 | 1.81199283 | Autosome |
| CHAAL00000012354 | ATCAY | 1.81164156 | Autosome |
| CHAAL00000010699 | ANKEF1 | 1.80785235 | Autosome |
| CHAAL00000011471 | GMNN | 1.80785235 | Autosome |
| CHAAL00000014453 | HOMER2 | 1.80785235 | Autosome |
| CHAAL00000008678 | DCLK3 | 1.80172392 | Autosome |
| CHAAL00000014123 | PDCL2 | 1.80172392 | Autosome |
| CHAAL00000005428 | MYO3A | 1.80046626 | Autosome |
| CHAAL00000003906 | | 1.79798243 | Autosome |
| CHAAL00000007197 | SERPINB10 | 1.79795831 | Autosome |
| CHAAL00000014737 | TMPRSS6 | 1.79795831 | Autosome |
| CHAAL00000002769 | NCOA7 | 1.79015812 | Autosome |
| CHAAL00000007131 | KDM5B | 1.79015812 | Autosome |
| CHAAL00000007397 | KAZALD1 | 1.78429628 | Autosome |
| CHAAL00000005496 | PEX1 | 1.78370928 | Autosome |
| CHAAL00000009047 | LOC104382800 | 1.78222526 | Autosome |
| CHAAL00000001295 | RIC8A | 1.78188864 | Autosome |
| CHAAL00000002017 | CTPS2 | 1.78188864 | Autosome |
| CHAAL00000002623 | EPHA1 | 1.78082493 | Autosome |
| CHAAL00000003098 | HCK | 1.78082493 | Autosome |
| CHAAL00000004551 | NCR3LG1 | 1.78082493 | Autosome |
| CHAAL00000011298 | FBXO5 | 1.78082493 | Autosome |
| CHAAL00000008344 | | 1.7752616 | Autosome |
| CHAAL00000004924 | | 1.77435874 | Autosome |
| CHAAL00000009439 | KCNG4 | 1.77371982 | Autosome |
| CHAAL00000010247 | NEDD4 | 1.77371982 | Autosome |
| CHAAL00000010933 | TRMT2A | 1.77197101 | Autosome |

| | | | |
|---|---|---|---|
| CHAAL00000014251 | | 1.77197101 | Autosome |
| CHAAL00000000398 | | 1.77155894 | Autosome |
| CHAAL00000000394 | BORCS5 | 1.76767256 | Autosome |
| CHAAL00000004080 | LOC103539723 | 1.76767256 | Autosome |
| CHAAL00000012084 | CCT5 | 1.76767256 | Autosome |
| CHAAL00000014033 | LOC104571010 | 1.76767256 | Autosome |
| CHAAL00000012478 | CEP95 | 1.7669523 | Autosome |
| CHAAL00000002192 | ANKRD13A | 1.76649221 | Autosome |
| CHAAL00000013671 | ANO1 | 1.76649221 | Autosome |
| CHAAL00000015663 | | 1.76384689 | Autosome |
| CHAAL00000015621 | LOC104071961 | 1.76229297 | Autosome |
| CHAAL00000012056 | EFTUD2 | 1.76136664 | Autosome |
| CHAAL00000002919 | PARP12 | 1.75517383 | Autosome |
| CHAAL00000010881 | OGFOD2 | 1.75507648 | Autosome |
| CHAAL00000010937 | | 1.7543528 | Autosome |
| CHAAL00000005728 | DRC3 | 1.7537975 | Autosome |
| CHAAL00000004559 | KCNK18 | 1.75126997 | Autosome |
| CHAAL00000008422 | YAE1D1 | 1.75104883 | Autosome |
| CHAAL00000000793 | LOC104291268 | 1.75086612 | Autosome |
| CHAAL00000008760 | FNIP1 | 1.75086612 | Autosome |
| CHAAL00000000789 | ACAA1 | 1.74879131 | Autosome |
| CHAAL00000002304 | RBM17 | 1.74879131 | Autosome |
| CHAAL00000002441 | | 1.74879131 | Autosome |
| CHAAL00000004444 | JCHAIN | 1.74879131 | Autosome |
| CHAAL00000004799 | EDN1 | 1.74879131 | Autosome |
| CHAAL00000005517 | PEX10 | 1.74879131 | Autosome |
| CHAAL00000005770 | LOC104283360 | 1.74879131 | Autosome |
| CHAAL00000006640 | RBMS1 | 1.74879131 | Autosome |
| CHAAL00000006666 | PPA1 | 1.74879131 | Autosome |
| CHAAL00000007825 | NAIF1 | 1.74879131 | Autosome |
| CHAAL00000009825 | CMAS | 1.74879131 | Autosome |
| CHAAL00000009837 | MED20 | 1.74879131 | Autosome |
| CHAAL00000012305 | PNOC | 1.74879131 | Autosome |
| CHAAL00000012361 | KLF5 | 1.74879131 | Autosome |
| CHAAL00000013145 | PTDSS1 | 1.74879131 | Autosome |
| CHAAL00000013743 | RPL27 | 1.74879131 | Autosome |
| CHAAL00000015493 | LOC106019833 | 1.74879131 | Autosome |
| CHAAL00000001215 | CCT3 | 1.74028379 | Autosome |
| CHAAL00000010667 | CCDC88A | 1.73943927 | Autosome |
| CHAAL00000007319 | KLHL33 | 1.73905089 | Autosome |
| CHAAL00000011241 | TTC4 | 1.73905089 | Autosome |
| CHAAL00000006938 | GALNT18 | 1.73753566 | Autosome |
| CHAAL00000015661 | RHNO1 | 1.73442511 | Autosome |

| | | | |
|---|---|---|---|
| CHAAL00000000806 | NCR3LG1 | 1.73384075 | Autosome |
| CHAAL00000015494 | MYO1G | 1.73230045 | Autosome |
| CHAAL00000001634 | RPSA | 1.72801242 | Autosome |
| CHAAL00000009868 | MBTPS1 | 1.72683699 | Autosome |
| CHAAL00000002900 | ADCK2 | 1.72677007 | Autosome |
| CHAAL00000010896 | ATP6V0A2 | 1.72677007 | Autosome |
| CHAAL00000002838 | ANKRD28 | 1.72020184 | Autosome |
| CHAAL00000004933 | | 1.71999485 | Autosome |
| CHAAL00000006041 | CFH | 1.71929307 | Autosome |
| CHAAL00000006145 | | 1.71929307 | Autosome |
| CHAAL00000009106 | ERCC4 | 1.71929307 | Autosome |
| CHAAL00000000395 | LOC104281694 | 1.71874236 | Autosome |
| CHAAL00000009638 | APPBP2 | 1.71780138 | Autosome |
| CHAAL00000004507 | FRMD4A | 1.71658557 | Autosome |
| CHAAL00000006095 | LOC104251070 | 1.71658557 | Autosome |
| CHAAL00000008032 | HTRA3 | 1.71658557 | Autosome |
| CHAAL00000003692 | RWDD2A | 1.71246517 | Autosome |
| CHAAL00000007813 | COQ9 | 1.70356003 | Autosome |
| CHAAL00000004610 | OTOP3 | 1.70254754 | Autosome |
| CHAAL00000012213 | FAM91A1 | 1.70254754 | Autosome |
| CHAAL00000004477 | FN1 | 1.70245403 | Autosome |
| CHAAL00000000479 | C1QTNF2 | 1.70117766 | Autosome |
| CHAAL00000011960 | MTPAP | 1.70117766 | Autosome |
| CHAAL00000000774 | IL13RA1 | 1.69974265 | Autosome |
| CHAAL00000000810 | EOMES | 1.69974265 | Autosome |
| CHAAL00000005846 | NCOA2 | 1.69974265 | Autosome |
| CHAAL00000005987 | SNX7 | 1.69974265 | Autosome |
| CHAAL00000010686 | XPO1 | 1.69974265 | Autosome |
| CHAAL00000010887 | KMT5A | 1.69974265 | Autosome |
| CHAAL00000015511 | GRK7 | 1.69974265 | Autosome |
| CHAAL00000010751 | PREX1 | 1.69809569 | Autosome |
| CHAAL00000004800 | HIVEP1 | 1.69532988 | Autosome |
| CHAAL00000000695 | | 1.69373187 | Autosome |
| CHAAL00000004405 | SLC7A6 | 1.69373187 | Autosome |
| CHAAL00000005109 | WNT16 | 1.68782698 | Autosome |
| CHAAL00000009819 | LRMP | 1.68782698 | Autosome |
| CHAAL00000007191 | LOC104286660 | 1.68755397 | Autosome |
| CHAAL00000000581 | | 1.68622893 | Autosome |
| CHAAL00000004097 | GET4 | 1.68622893 | Autosome |
| CHAAL00000007210 | RIPK1 | 1.68455393 | Autosome |
| CHAAL00000009867 | HSDL1 | 1.68455393 | Autosome |
| CHAAL00000001704 | GABRA2 | 1.68312097 | Autosome |
| CHAAL00000005315 | UBE2E3 | 1.68312097 | Autosome |

| | | | |
|---|---|---|---|
| CHAAL00000007518 | WNT5B | 1.68312097 | Autosome |
| CHAAL00000008555 | CHST10 | 1.68312097 | Autosome |
| CHAAL00000013746 | | 1.68312097 | Autosome |
| CHAAL00000003935 | | 1.68230502 | Autosome |
| CHAAL00000006742 | SPSB3 | 1.68230502 | Autosome |
| CHAAL00000011335 | NMBR | 1.68230502 | Autosome |
| CHAAL00000011624 | CAB39L | 1.68230502 | Autosome |
| CHAAL00000009671 | MRM3 | 1.68102439 | Autosome |
| CHAAL00000002572 | RET | 1.67449338 | Autosome |
| CHAAL00000006649 | | 1.67199609 | Autosome |
| CHAAL00000009115 | LOC103917624 | 1.67087817 | Autosome |
| CHAAL00000013042 | GPATCH1 | 1.67087817 | Autosome |
| CHAAL00000006990 | HDAC10 | 1.67038526 | Autosome |

**Supplementary Table 21** Low *Tajima's D* genes and gene descriptions. Genes were considered to have low *Tajima's D* if they fell below the 5$^{th}$ percentile for negative genes (autosomes: *Tajima's D* < -1.179, Z: *Tajima's D* < -1.234). Gene symbols were assigned based on best reciprocal BLAST matches from the RefSeq protein database.

| Gene Name | Gene Symbol | *Tajima's D* | Chromosome |
|---|---|---|---|
| CHAAL00000000621 | | -2.0562375 | Autosome |
| CHAAL00000002674 | LOC104293163 | -1.9229072 | Autosome |
| CHAAL00000000904 | PRKCH | -1.8678777 | Autosome |
| CHAAL00000004811 | IFNGR2 | -1.8678777 | Autosome |
| CHAAL00000014286 | | -1.8678777 | Autosome |
| CHAAL00000000241 | FAM169B | -1.7233098 | Autosome |
| CHAAL00000001274 | LOC106894171 | -1.7233098 | Autosome |
| CHAAL00000001723 | GNA12 | -1.7233098 | Autosome |
| CHAAL00000001791 | GJA8 | -1.7233098 | Autosome |
| CHAAL00000002088 | | -1.7233098 | Autosome |
| CHAAL00000002229 | UPF3A | -1.7233098 | Autosome |
| CHAAL00000004499 | ADCYAP1R1 | -1.7233098 | Autosome |
| CHAAL00000010311 | KCNA4 | -1.7233098 | Autosome |
| CHAAL00000010729 | LOC104290288 | -1.7233098 | Autosome |
| CHAAL00000011484 | MSANTD3 | -1.7233098 | Autosome |
| CHAAL00000013552 | ZRANB1 | -1.7233098 | Autosome |

| | | | |
|---|---|---|---|
| CHAAL00000013182 | PRLR | -1.7233098 | Z |
| CHAAL00000008549 | CNGA2 | -1.6916942 | Autosome |
| CHAAL00000004071 | DAGLB | -1.6381446 | Autosome |
| CHAAL00000004649 | LOC104350811 | -1.6381446 | Autosome |
| CHAAL00000004787 | | -1.6381446 | Autosome |
| CHAAL00000006679 | | -1.6381446 | Autosome |
| CHAAL00000008432 | | -1.6381446 | Autosome |
| CHAAL00000008858 | CNOT8 | -1.6381446 | Autosome |
| CHAAL00000011013 | LOC104289611 | -1.6381446 | Autosome |
| CHAAL00000012533 | LMF2 | -1.6381446 | Autosome |
| CHAAL00000001969 | | -1.6099437 | Autosome |
| CHAAL00000014756 | TRIOBP | -1.5900289 | Autosome |
| CHAAL00000008686 | | -1.585774 | Autosome |
| CHAAL00000009591 | SWT1 | -1.585774 | Autosome |
| CHAAL00000010352 | PRDM16 | -1.585774 | Autosome |
| CHAAL00000015191 | SEC24D | -1.5702732 | Autosome |
| CHAAL00000005089 | SELENOI | -1.551485 | Autosome |
| CHAAL00000000413 | FAM118A | -1.5128357 | Autosome |
| CHAAL00000000529 | LOC104282407 | -1.5128357 | Autosome |
| CHAAL00000000716 | HDX | -1.5128357 | Autosome |
| CHAAL00000001568 | ARMC10 | -1.5128357 | Autosome |
| CHAAL00000002287 | SOWAHC | -1.5128357 | Autosome |
| CHAAL00000002532 | PHYHIPL | -1.5128357 | Autosome |
| CHAAL00000002540 | LDB3 | -1.5128357 | Autosome |
| CHAAL00000003104 | NOL4L | -1.5128357 | Autosome |
| CHAAL00000003148 | SEL1L | -1.5128357 | Autosome |
| CHAAL00000003303 | MBOAT4 | -1.5128357 | Autosome |
| CHAAL00000003494 | | -1.5128357 | Autosome |
| CHAAL00000003921 | GUCY1A2 | -1.5128357 | Autosome |
| CHAAL00000004346 | USP15 | -1.5128357 | Autosome |
| CHAAL00000004396 | LOC104125202 | -1.5128357 | Autosome |
| CHAAL00000005106 | MYOCD | -1.5128357 | Autosome |
| CHAAL00000005178 | FAM167B | -1.5128357 | Autosome |
| CHAAL00000005263 | SLC20A1 | -1.5128357 | Autosome |
| CHAAL00000005576 | LOC104259182 | -1.5128357 | Autosome |
| CHAAL00000005904 | USP33 | -1.5128357 | Autosome |
| CHAAL00000006370 | TMED8 | -1.5128357 | Autosome |
| CHAAL00000006459 | AP4S1 | -1.5128357 | Autosome |
| CHAAL00000006506 | ARG2 | -1.5128357 | Autosome |
| CHAAL00000007073 | PUDP | -1.5128357 | Autosome |
| CHAAL00000007259 | LYPD6B | -1.5128357 | Autosome |
| CHAAL00000007425 | MORN4 | -1.5128357 | Autosome |
| CHAAL00000007852 | ZER1 | -1.5128357 | Autosome |

| | | | |
|---|---|---|---|
| CHAAL00000007987 | LOC101876505 | -1.5128357 | Autosome |
| CHAAL00000007997 | ZC2HC1A | -1.5128357 | Autosome |
| CHAAL00000008071 | CTBP1 | -1.5128357 | Autosome |
| CHAAL00000008295 | LOC104285964 | -1.5128357 | Autosome |
| CHAAL00000008305 | BIRC5 | -1.5128357 | Autosome |
| CHAAL00000008488 | ADAM11 | -1.5128357 | Autosome |
| CHAAL00000008558 | AFF3 | -1.5128357 | Autosome |
| CHAAL00000008852 | MFAP3 | -1.5128357 | Autosome |
| CHAAL00000009398 | EAF1 | -1.5128357 | Autosome |
| CHAAL00000009983 | LOC104282979 | -1.5128357 | Autosome |
| CHAAL00000010053 | RND2 | -1.5128357 | Autosome |
| CHAAL00000010575 | KLC4 | -1.5128357 | Autosome |
| CHAAL00000010597 | SLC35A4 | -1.5128357 | Autosome |
| CHAAL00000010913 | STX2 | -1.5128357 | Autosome |
| CHAAL00000011009 | SLC5A1 | -1.5128357 | Autosome |
| CHAAL00000011169 | ZC3H15 | -1.5128357 | Autosome |
| CHAAL00000011527 | ZMYND12 | -1.5128357 | Autosome |
| CHAAL00000011758 | PIP4K2B | -1.5128357 | Autosome |
| CHAAL00000011764 | | -1.5128357 | Autosome |
| CHAAL00000012178 | LOC108508529 | -1.5128357 | Autosome |
| CHAAL00000012342 | LOC101910757 | -1.5128357 | Autosome |
| CHAAL00000013335 | LOC104829944 | -1.5128357 | Autosome |
| CHAAL00000013419 | LACTBL1 | -1.5128357 | Autosome |
| CHAAL00000013501 | HNRNPM | -1.5128357 | Autosome |
| CHAAL00000013708 | PQLC3 | -1.5128357 | Autosome |
| CHAAL00000014418 | | -1.5128357 | Autosome |
| CHAAL00000014684 | GINS4 | -1.5128357 | Autosome |
| CHAAL00000014965 | LOC104282713 | -1.5128357 | Autosome |
| CHAAL00000003547 | MED18 | -1.5128357 | Z |
| CHAAL00000013968 | TMED7 | -1.5128357 | Z |
| CHAAL00000005705 | LOC106888801 | -1.4923492 | Autosome |
| CHAAL00000004698 | AGRN | -1.4696085 | Autosome |
| CHAAL00000006270 | LOC104292474 | -1.4600787 | Autosome |
| CHAAL00000008179 | NEDD4L | -1.4600787 | Z |
| CHAAL00000004491 | AGAP1 | -1.4413439 | Autosome |
| CHAAL00000000647 | GLRA4 | -1.4407064 | Autosome |
| CHAAL00000001152 | TMX1 | -1.4407064 | Autosome |
| CHAAL00000001640 | SLC37A1 | -1.4407064 | Autosome |
| CHAAL00000003086 | REM1 | -1.4407064 | Autosome |
| CHAAL00000005151 | LOC103903537 | -1.4407064 | Autosome |
| CHAAL00000005574 | LOC103918196 | -1.4407064 | Autosome |
| CHAAL00000005747 | TLK2 | -1.4407064 | Autosome |
| CHAAL00000006064 | TLR4 | -1.4407064 | Autosome |

| | | | |
|---|---|---|---|
| CHAAL00000006618 | TUBA4A | -1.4407064 | Autosome |
| CHAAL00000007276 | CCNL1 | -1.4407064 | Autosome |
| CHAAL00000007598 | ETNK2 | -1.4407064 | Autosome |
| CHAAL00000007615 | | -1.4407064 | Autosome |
| CHAAL00000008707 | BACE1 | -1.4407064 | Autosome |
| CHAAL00000008796 | NME5 | -1.4407064 | Autosome |
| CHAAL00000008907 | DDX41 | -1.4407064 | Autosome |
| CHAAL00000009055 | LOC101880026 | -1.4407064 | Autosome |
| CHAAL00000009801 | LOC104076975 | -1.4407064 | Autosome |
| CHAAL00000010452 | TMEM38A | -1.4407064 | Autosome |
| CHAAL00000010518 | TNRC6B | -1.4407064 | Autosome |
| CHAAL00000012956 | NKRF | -1.4407064 | Autosome |
| CHAAL00000013013 | TMEM68, LOC106894137 | -1.4407064 | Autosome |
| CHAAL00000013360 | C1QBP | -1.4407064 | Autosome |
| CHAAL00000013481 | SLITRK3 | -1.4407064 | Autosome |
| CHAAL00000013502 | YJU2 | -1.4407064 | Autosome |
| CHAAL00000013807 | MECOM | -1.4407064 | Autosome |
| CHAAL00000014068 | | -1.4407064 | Autosome |
| CHAAL00000014221 | LOC106896931 | -1.4407064 | Autosome |
| CHAAL00000014246 | GTF2E1 | -1.4407064 | Autosome |
| CHAAL00000015554 | FER1L5 | -1.4407064 | Autosome |
| CHAAL00000013880 | TAF1C | -1.4407064 | Z |
| CHAAL00000013188 | SKP2 | -1.4407064 | Z |
| CHAAL00000000149 | LOC104832430 | -1.4407064 | Z |
| CHAAL00000011824 | XRCC4 | -1.4407064 | Z |
| CHAAL00000015124 | LOC104387136 | -1.4354389 | Autosome |
| CHAAL00000003587 | SREK1 | -1.4354389 | Z |
| CHAAL00000002069 | MED6 | -1.4143713 | Autosome |
| CHAAL00000002190 | TCHP | -1.4143713 | Autosome |
| CHAAL00000004182 | LOC104289051 | -1.4143713 | Autosome |
| CHAAL00000013121 | ZC3H10 | -1.4143713 | Autosome |
| CHAAL00000015188 | | -1.4143713 | Z |
| CHAAL00000000083 | RIC3 | -1.4084114 | Autosome |
| CHAAL00000000864 | SIGIRR | -1.4084114 | Autosome |
| CHAAL00000001676 | ERBB2 | -1.4084114 | Autosome |
| CHAAL00000002559 | GHITM | -1.4084114 | Autosome |
| CHAAL00000002722 | BACH1 | -1.4084114 | Autosome |
| CHAAL00000002950 | LHFPL6 | -1.4084114 | Autosome |
| CHAAL00000003073 | PSMF1 | -1.4084114 | Autosome |
| CHAAL00000004237 | AMT | -1.4084114 | Autosome |
| CHAAL00000005127 | INSRR | -1.4084114 | Autosome |
| CHAAL00000007988 | GDAP1 | -1.4084114 | Autosome |
| CHAAL00000010019 | ALDH4A1 | -1.4084114 | Autosome |

| | | | |
|---|---|---|---|
| CHAAL00000014700 | RIC8B | -1.4084114 | Autosome |
| CHAAL00000015185 | SLC12A2 | -1.4084114 | Z |
| CHAAL00000000231 | LINS1 | -1.4016733 | Autosome |
| CHAAL00000000756 | ADGRG4 | -1.4016733 | Autosome |
| CHAAL00000013343 | GTF2IRD1 | -1.3935003 | Autosome |
| CHAAL00000000665 | ZC3H12B | -1.3915176 | Autosome |
| CHAAL00000012215 | KLHL38 | -1.3915176 | Autosome |
| CHAAL00000014518 | RFX5 | -1.381838 | Autosome |
| CHAAL00000001829 | | -1.3757198 | Autosome |
| CHAAL00000004366 | | -1.3486986 | Autosome |
| CHAAL00000011694 | LOC104281733 | -1.3415432 | Autosome |
| CHAAL00000001826 | | -1.3386743 | Autosome |
| CHAAL00000005425 | APBB1IP | -1.3322498 | Autosome |
| CHAAL00000001626 | SUSD5 | -1.3029915 | Autosome |
| CHAAL00000003310 | TNKS | -1.2970258 | Autosome |
| CHAAL00000013446 | LOC104276419 | -1.2913717 | Autosome |
| CHAAL00000009884 | TPO | -1.2899324 | Autosome |
| CHAAL00000011912 | COQ8A | -1.2771021 | Autosome |
| CHAAL00000006030 | ZNF281 | -1.272744 | Autosome |
| CHAAL00000013616 | LOC106899268, LOC104035844 | -1.272744 | Autosome |
| CHAAL00000005000 | MYB | -1.2707146 | Autosome |
| CHAAL00000000057 | FGF23 | -1.2658222 | Autosome |
| CHAAL00000005214 | HDAC11 | -1.2658222 | Autosome |
| CHAAL00000011565 | LARGE2 | -1.2658222 | Autosome |
| CHAAL00000014999 | TMEM200C | -1.2658222 | Autosome |
| CHAAL00000015503 | DAW1 | -1.2658222 | Autosome |
| CHAAL00000014323 | PCDH1 | -1.2619442 | Autosome |
| CHAAL00000001205 | CHRNB2 | -1.2597606 | Autosome |
| CHAAL00000008710 | TMPRSS13 | -1.2597606 | Autosome |
| CHAAL00000013835 | PPM1K | -1.2529087 | Autosome |
| CHAAL00000011693 | | -1.2444034 | Autosome |
| CHAAL00000006092 | LOC104264308 | -1.2429911 | Autosome |
| CHAAL00000008505 | GGT7 | -1.2429685 | Autosome |
| CHAAL00000010993 | LOC104293691 | -1.2429685 | Autosome |
| CHAAL00000012718 | SIK1 | -1.2429685 | Autosome |
| CHAAL00000008279 | GRIN3A | -1.2385197 | Z |
| CHAAL00000000309 | MOB3C | -1.2330735 | Autosome |
| CHAAL00000000668 | MSN | -1.2330735 | Autosome |
| CHAAL00000003900 | SLC34A1 | -1.2330735 | Autosome |
| CHAAL00000003919 | KBTBD3 | -1.2330735 | Autosome |
| CHAAL00000015471 | HRH3 | -1.2330735 | Autosome |
| CHAAL00000009430 | PLCG2 | -1.2306531 | Autosome |
| CHAAL00000005679 | ALDH9A1 | -1.2287952 | Autosome |

| | | | |
|---|---|---|---|
| CHAAL00000015153 | ACTA1 | -1.2268625 | Autosome |
| CHAAL00000003804 | SHMT1 | -1.2227748 | Autosome |
| CHAAL00000000780 | AGTR2 | -1.2201148 | Autosome |
| CHAAL00000000902 | TRMT5 | -1.2201148 | Autosome |
| CHAAL00000004356 | TPP2 | -1.2201148 | Autosome |
| CHAAL00000006063 | BRINP1 | -1.2201148 | Autosome |
| CHAAL00000007641 | | -1.2201148 | Autosome |
| CHAAL00000007995 | PEX2 | -1.2201148 | Autosome |
| CHAAL00000008102 | MRPL21 | -1.2201148 | Autosome |
| CHAAL00000009279 | LRRC18 | -1.2201148 | Autosome |
| CHAAL00000009374 | CXCR6 | -1.2201148 | Autosome |
| CHAAL00000014997 | EPB41L3 | -1.2201148 | Autosome |
| CHAAL00000013090 | RPN1 | -1.218107 | Autosome |
| CHAAL00000015154 | | -1.2132382 | Autosome |
| CHAAL00000009238 | TTC7A | -1.2115775 | Autosome |
| CHAAL00000014748 | GGA1 | -1.2107342 | Autosome |
| CHAAL00000001330 | SF3A2 | -1.2057057 | Autosome |
| CHAAL00000002174 | SSH1 | -1.2057057 | Autosome |
| CHAAL00000005193 | KIAA1522 | -1.2057057 | Autosome |
| CHAAL00000006532 | BAALC | -1.2057057 | Autosome |
| CHAAL00000007177 | | -1.2057057 | Autosome |
| CHAAL00000007793 | | -1.2057057 | Autosome |
| CHAAL00000011338 | MATN3 | -1.2057057 | Autosome |
| CHAAL00000011552 | DLL4 | -1.2057057 | Autosome |
| CHAAL00000013389 | | -1.2057057 | Autosome |
| CHAAL00000013972 | CTTNBP2NL | -1.2057057 | Autosome |
| CHAAL00000014986 | LOC104281974 | -1.2057057 | Autosome |
| CHAAL00000013585 | | -1.2005513 | Autosome |
| CHAAL00000006951 | ZNF143 | -1.1985183 | Autosome |
| CHAAL00000004367 | | -1.1972611 | Autosome |
| CHAAL00000006115 | TRIM41 | -1.1972611 | Autosome |
| CHAAL00000007238 | ABHD18 | -1.1972611 | Autosome |
| CHAAL00000008448 | CLN5 | -1.1972611 | Autosome |
| CHAAL00000002020 | ZRSR2 | -1.1913506 | Autosome |
| CHAAL00000003079 | SRXN1 | -1.1913506 | Autosome |
| CHAAL00000004107 | SRSF11 | -1.1913506 | Autosome |
| CHAAL00000004638 | LOC103907554 | -1.1913506 | Autosome |
| CHAAL00000005112 | TSPAN12 | -1.1913506 | Autosome |
| CHAAL00000005358 | SP3 | -1.1913506 | Autosome |
| CHAAL00000005875 | OLFM4 | -1.1913506 | Autosome |
| CHAAL00000006395 | ACYP1 | -1.1913506 | Autosome |
| CHAAL00000006958 | ASCL3 | -1.1913506 | Autosome |
| CHAAL00000007261 | MMADHC | -1.1913506 | Autosome |

| | | | |
|---|---|---|---|
| CHAAL00000008413 | CCKBR | -1.1913506 | Autosome |
| CHAAL00000008491 | CPB2 | -1.1913506 | Autosome |
| CHAAL00000009163 | IMPA2 | -1.1913506 | Autosome |
| CHAAL00000009941 | GATA6 | -1.1913506 | Autosome |
| CHAAL00000010956 | YDJC | -1.1913506 | Autosome |
| CHAAL00000011594 | FAM180B | -1.1913506 | Autosome |
| CHAAL00000012060 | KIF18B | -1.1913506 | Autosome |
| CHAAL00000012304 | LOC104295673 | -1.1913506 | Autosome |
| CHAAL00000013333 | RBM12 | -1.1913506 | Autosome |
| CHAAL00000013493 | ZBTB7A | -1.1913506 | Autosome |
| CHAAL00000013564 | CLRN3 | -1.1913506 | Autosome |
| CHAAL00000014392 | GOLPH3L | -1.1913506 | Autosome |
| CHAAL00000015148 | SPRTN | -1.1913506 | Autosome |
| CHAAL00000015261 | LOC101880644 | -1.1913506 | Autosome |
| CHAAL00000003483 | RBBP8NL | -1.1843502 | Autosome |
| CHAAL00000002244 | PROZ | -1.1830105 | Autosome |

**Supplementary Table 22** Low *Tajima's D* genes and gene descriptions. Genes were considered to have low *Tajima's D* if they fell below the 5$^{th}$ percentile for negative genes (autosomes: *Tajima's D* ≤ -1.191, Z: *Tajima's D* ≤ -1.445). Gene symbols were assigned based on best reciprocal BLAST matches from the RefSeq protein database.

| Gene Name | Gene Symbol | *Tajima's D* | Chromosome |
|---|---|---|---|
| CHAAL00000002390 | | -1.8678777 | Autosome |
| CHAAL00000014524 | TMOD4 | -1.8678777 | Autosome |
| CHAAL00000012049 | | -1.8206424 | Autosome |
| CHAAL00000001210 | HAX1 | -1.7233098 | Autosome |
| CHAAL00000002278 | RFT1 | -1.7233098 | Autosome |
| CHAAL00000003609 | SDC1 | -1.7233098 | Autosome |
| CHAAL00000006378 | CIPC | -1.7233098 | Autosome |
| CHAAL00000006438 | NFKBIA | -1.7233098 | Autosome |
| CHAAL00000009321 | DHX36 | -1.7233098 | Autosome |
| CHAAL00000009809 | INTS13 | -1.7233098 | Autosome |
| CHAAL00000009832 | LOC104289366 | -1.7233098 | Autosome |
| CHAAL00000010950 | PPIL2 | -1.7233098 | Autosome |
| CHAAL00000011911 | PSEN2 | -1.7233098 | Autosome |
| CHAAL00000014316 | HDAC3 | -1.7233098 | Autosome |
| CHAAL00000014665 | SH2B3 | -1.7233098 | Autosome |

| | | | |
|---|---|---|---|
| CHAAL00000003441 | | -1.7233098 | Z |
| CHAAL00000015584 | EYS | -1.7190377 | Autosome |
| CHAAL00000011167 | FAM171B | -1.6741534 | Autosome |
| CHAAL00000004203 | LOC104291592 | -1.6381446 | Autosome |
| CHAAL00000005780 | AK7 | -1.6381446 | Autosome |
| CHAAL00000009079 | COQ7 | -1.6381446 | Autosome |
| CHAAL00000011338 | MATN3 | -1.6381446 | Autosome |
| CHAAL00000012871 | DOCK9 | -1.6381446 | Autosome |
| CHAAL00000012912 | MAGT1 | -1.6381446 | Autosome |
| CHAAL00000012964 | LONRF3 | -1.6381446 | Autosome |
| CHAAL00000004572 | | -1.6099437 | Autosome |
| CHAAL00000006975 | | -1.60012 | Autosome |
| CHAAL00000003384 | LOC104258112 | -1.5960355 | Autosome |
| CHAAL00000008632 | ABCA2 | -1.5902928 | Autosome |
| CHAAL00000000455 | NEURL1B | -1.585774 | Autosome |
| CHAAL00000007837 | CERCAM | -1.585774 | Autosome |
| CHAAL00000012583 | LAMB2 | -1.585774 | Autosome |
| CHAAL00000012175 | LOC104020593 | -1.5702732 | Autosome |
| CHAAL00000008034 | TRMT44 | -1.5504379 | Autosome |
| CHAAL00000011817 | AIFM1 | -1.5504379 | Autosome |
| CHAAL00000001593 | | -1.5339538 | Z |
| CHAAL00000001383 | TRIM36 | -1.5301909 | Z |
| CHAAL00000001384 | | -1.5301909 | Z |
| CHAAL00000000252 | KLHL25 | -1.5128357 | Autosome |
| CHAAL00000000750 | RBMX | -1.5128357 | Autosome |
| CHAAL00000000855 | CDK5RAP3 | -1.5128357 | Autosome |
| CHAAL00000001110 | IFT88 | -1.5128357 | Autosome |
| CHAAL00000001112 | LOC104291399 | -1.5128357 | Autosome |
| CHAAL00000001523 | TRMU | -1.5128357 | Autosome |
| CHAAL00000001835 | LOC104283002 | -1.5128357 | Autosome |
| CHAAL00000001912 | ARRDC5 | -1.5128357 | Autosome |
| CHAAL00000002175 | DAO | -1.5128357 | Autosome |
| CHAAL00000002185 | MMAB | -1.5128357 | Autosome |
| CHAAL00000002515 | REEP3 | -1.5128357 | Autosome |
| CHAAL00000003307 | PPP1R3B | -1.5128357 | Autosome |
| CHAAL00000003407 | CCNT1 | -1.5128357 | Autosome |
| CHAAL00000003731 | CISH | -1.5128357 | Autosome |
| CHAAL00000004120 | NGF | -1.5128357 | Autosome |
| CHAAL00000004320 | MIER2 | -1.5128357 | Autosome |
| CHAAL00000004822 | KRT222 | -1.5128357 | Autosome |
| CHAAL00000005470 | NMT2 | -1.5128357 | Autosome |
| CHAAL00000005523 | TMEM52 | -1.5128357 | Autosome |
| CHAAL00000005592 | RBPJL | -1.5128357 | Autosome |

| | | | |
|---|---|---|---|
| CHAAL00000005675 | LMX1A | -1.5128357 | Autosome |
| CHAAL00000005925 | LPAR3 | -1.5128357 | Autosome |
| CHAAL00000006101 | NDUFS8 | -1.5128357 | Autosome |
| CHAAL00000006427 | SSTR1 | -1.5128357 | Autosome |
| CHAAL00000007007 | SLC13A1 | -1.5128357 | Autosome |
| CHAAL00000007607 | LOC103912354 | -1.5128357 | Autosome |
| CHAAL00000007843 | LOC104023141 | -1.5128357 | Autosome |
| CHAAL00000007888 | PLPP7 | -1.5128357 | Autosome |
| CHAAL00000008191 | MAPK1IP1L | -1.5128357 | Autosome |
| CHAAL00000008975 | LGALS8 | -1.5128357 | Autosome |
| CHAAL00000009127 | USP7 | -1.5128357 | Autosome |
| CHAAL00000009356 | TRAK1 | -1.5128357 | Autosome |
| CHAAL00000009421 | CDYL2 | -1.5128357 | Autosome |
| CHAAL00000009507 | UTP4 | -1.5128357 | Autosome |
| CHAAL00000009756 | LPGAT1 | -1.5128357 | Autosome |
| CHAAL00000009785 | | -1.5128357 | Autosome |
| CHAAL00000009998 | MYCL | -1.5128357 | Autosome |
| CHAAL00000010003 | LOC104284436 | -1.5128357 | Autosome |
| CHAAL00000010150 | CSK | -1.5128357 | Autosome |
| CHAAL00000010151 | ULK3 | -1.5128357 | Autosome |
| CHAAL00000010349 | TPRG1L | -1.5128357 | Autosome |
| CHAAL00000010372 | MECR | -1.5128357 | Autosome |
| CHAAL00000010864 | ORAI1 | -1.5128357 | Autosome |
| CHAAL00000011369 | CDON | -1.5128357 | Autosome |
| CHAAL00000011579 | ARFGAP2 | -1.5128357 | Autosome |
| CHAAL00000012644 | TMEM33 | -1.5128357 | Autosome |
| CHAAL00000012900 | LOC101939167 | -1.5128357 | Autosome |
| CHAAL00000013011 | XKR4 | -1.5128357 | Autosome |
| CHAAL00000013819 | RPL22L1 | -1.5128357 | Autosome |
| CHAAL00000014306 | MRPS26 | -1.5128357 | Autosome |
| CHAAL00000014340 | TNNT3 | -1.5128357 | Autosome |
| CHAAL00000014356 | TMEM82 | -1.5128357 | Autosome |
| CHAAL00000014824 | IVD | -1.5128357 | Autosome |
| CHAAL00000015321 | | -1.5128357 | Autosome |
| CHAAL00000015348 | KLF6 | -1.5128357 | Autosome |
| CHAAL00000015431 | CCDC124 | -1.5128357 | Autosome |
| CHAAL00000015440 | ANO8 | -1.5128357 | Autosome |
| CHAAL00000015571 | BMP15 | -1.5128357 | Autosome |
| CHAAL00000003591 | TRAPPC13 | -1.5128357 | Z |
| CHAAL00000008261 | | -1.5128357 | Z |
| CHAAL00000011659 | ARRDC3 | -1.5128357 | Z |
| CHAAL00000014556 | DNAJB5 | -1.5128357 | Z |
| CHAAL00000015060 | LMNB1 | -1.5128357 | Z |

| | | | |
|---|---|---|---|
| CHAAL00000012442 | | -1.4703947 | Z |
| CHAAL00000000404 | TTC38 | -1.4600787 | Autosome |
| CHAAL00000003269 | | -1.4600787 | Autosome |
| CHAAL00000009771 | | -1.4556897 | Autosome |
| CHAAL00000000696 | KIF4A | -1.4407064 | Autosome |
| CHAAL00000001130 | PODXL | -1.4407064 | Autosome |
| CHAAL00000001160 | GPR137C | -1.4407064 | Autosome |
| CHAAL00000001179 | NXPH2 | -1.4407064 | Autosome |
| CHAAL00000002366 | MYADML2 | -1.4407064 | Autosome |
| CHAAL00000002722 | BACH1 | -1.4407064 | Autosome |
| CHAAL00000003052 | TPD52L2 | -1.4407064 | Autosome |
| CHAAL00000003353 | PARM1 | -1.4407064 | Autosome |
| CHAAL00000003361 | KCNV1 | -1.4407064 | Autosome |
| CHAAL00000005083 | MYO15A | -1.4407064 | Autosome |
| CHAAL00000005247 | SUPT20H | -1.4407064 | Autosome |
| CHAAL00000005644 | LOC106888974 | -1.4407064 | Autosome |
| CHAAL00000005901 | PIGK | -1.4407064 | Autosome |
| CHAAL00000006084 | DAGLA | -1.4407064 | Autosome |
| CHAAL00000007369 | CALHM1 | -1.4407064 | Autosome |
| CHAAL00000007709 | | -1.4407064 | Autosome |
| CHAAL00000007802 | ZNF319 | -1.4407064 | Autosome |
| CHAAL00000008439 | BLVRA | -1.4407064 | Autosome |
| CHAAL00000009103 | CPPED1 | -1.4407064 | Autosome |
| CHAAL00000009262 | SBF1 | -1.4407064 | Autosome |
| CHAAL00000009630 | MFSD9 | -1.4407064 | Autosome |
| CHAAL00000009907 | IAH1 | -1.4407064 | Autosome |
| CHAAL00000010343 | DFFB | -1.4407064 | Autosome |
| CHAAL00000010501 | LOC104017488 | -1.4407064 | Autosome |
| CHAAL00000011092 | APLNR | -1.4407064 | Autosome |
| CHAAL00000011217 | GPX7 | -1.4407064 | Autosome |
| CHAAL00000012393 | WSCD1 | -1.4407064 | Autosome |
| CHAAL00000013122 | PA2G4 | -1.4407064 | Autosome |
| CHAAL00000014286 | | -1.4407064 | Autosome |
| CHAAL00000003925 | ALKBH8 | -1.4354389 | Autosome |
| CHAAL00000006882 | RHAG | -1.4354389 | Autosome |
| CHAAL00000007459 | ZNF518A | -1.4354389 | Autosome |
| CHAAL00000007809 | ADGRG1 | -1.4354389 | Autosome |
| CHAAL00000011236 | TCEANC2 | -1.4354389 | Autosome |
| CHAAL00000008150 | WDR93 | -1.4285819 | Autosome |
| CHAAL00000006077 | TKFC | -1.4215086 | Autosome |
| CHAAL00000006011 | RNPC3 | -1.4143713 | Autosome |
| CHAAL00000007420 | ZDHHC16 | -1.4143713 | Autosome |
| CHAAL00000010704 | LOC101880985 | -1.4143713 | Autosome |

| | | | |
|---|---|---|---|
| CHAAL00000014436 | NR2C2AP | -1.4143713 | Autosome |
| CHAAL00000004365 | | -1.4110411 | Autosome |
| CHAAL00000000528 | | -1.4084114 | Autosome |
| CHAAL00000000553 | CRYBG2 | -1.4084114 | Autosome |
| CHAAL00000001878 | INTS12 | -1.4084114 | Autosome |
| CHAAL00000003035 | EEF1A2 | -1.4084114 | Autosome |
| CHAAL00000003456 | LOC101876685 | -1.4084114 | Autosome |
| CHAAL00000006023 | STXBP3 | -1.4084114 | Autosome |
| CHAAL00000010238 | ALDH1A2 | -1.4084114 | Autosome |
| CHAAL00000007661 | DPYSL2 | -1.4016733 | Autosome |
| CHAAL00000007935 | INPP5E | -1.3935003 | Autosome |
| CHAAL00000011386 | LOC104291555 | -1.3935003 | Autosome |
| CHAAL00000013502 | YJU2 | -1.3935003 | Autosome |
| CHAAL00000002557 | LRIT2 | -1.3915176 | Autosome |
| CHAAL00000003997 | LOC106894806 | -1.3915176 | Autosome |
| CHAAL00000004753 | TTK | -1.3915176 | Autosome |
| CHAAL00000007875 | FNBP1 | -1.3915176 | Autosome |
| CHAAL00000008817 | LOC104293149 | -1.3915176 | Autosome |
| CHAAL00000014042 | OSBP2 | -1.3915176 | Autosome |
| CHAAL00000007128 | RNPEP | -1.3722505 | Autosome |
| CHAAL00000010675 | | -1.3160507 | Autosome |
| CHAAL00000006013 | | -1.312249 | Autosome |
| CHAAL00000011413 | | -1.3095211 | Autosome |
| CHAAL00000001981 | | -1.3073667 | Autosome |
| CHAAL00000006896 | LAP3 | -1.2970258 | Autosome |
| CHAAL00000005444 | MLLT10 | -1.2935853 | Autosome |
| CHAAL00000006562 | PLD4 | -1.2925793 | Autosome |
| CHAAL00000003621 | | -1.279485 | Autosome |
| CHAAL00000006206 | XRCC2 | -1.2778523 | Autosome |
| CHAAL00000013723 | | -1.2739665 | Autosome |
| CHAAL00000000392 | | -1.2716464 | Autosome |
| CHAAL00000013985 | GUCA1A | -1.2707146 | Autosome |
| CHAAL00000002172 | SELPLG | -1.2658222 | Autosome |
| CHAAL00000005725 | RAI1 | -1.2658222 | Autosome |
| CHAAL00000014931 | MOGAT2 | -1.2658222 | Autosome |
| CHAAL00000007449 | CPN1 | -1.2597606 | Autosome |
| CHAAL00000002644 | GPR162 | -1.2529087 | Autosome |
| CHAAL00000004303 | PLPPR3 | -1.2529087 | Autosome |
| CHAAL00000005764 | LOC106885651 | -1.2529087 | Autosome |
| CHAAL00000008875 | UBLCP1 | -1.2485631 | Autosome |
| CHAAL00000000564 | EXTL1 | -1.2444034 | Autosome |
| CHAAL00000012071 | NSUN2 | -1.2444034 | Autosome |
| CHAAL00000015052 | STAT3 | -1.2444034 | Autosome |

| | | | |
|---|---|---|---|
| CHAAL00000001145 | MAP4K5 | -1.2442254 | Autosome |
| CHAAL00000001629 | GLB1 | -1.2429685 | Autosome |
| CHAAL00000009842 | LOC103919816 | -1.2330735 | Autosome |
| CHAAL00000000665 | ZC3H12B | -1.2201148 | Autosome |
| CHAAL00000001150 | PYGL | -1.2201148 | Autosome |
| CHAAL00000002715 | TMEM184C | -1.2201148 | Autosome |
| CHAAL00000006024 | GPSM2 | -1.2201148 | Autosome |
| CHAAL00000006661 | ADAMTS14 | -1.2201148 | Autosome |
| CHAAL00000009076 | TMC5 | -1.2201148 | Autosome |
| CHAAL00000011322 | RSPH3 | -1.2201148 | Autosome |
| CHAAL00000014358 | PLEKHM2 | -1.2201148 | Autosome |
| CHAAL00000015067 | PNPLA8 | -1.2201148 | Autosome |
| CHAAL00000004217 | CLDN34 | -1.2149201 | Autosome |
| CHAAL00000006350 | ACAD10 | -1.2132382 | Autosome |
| CHAAL00000010819 | ZSWIM1 | -1.2132382 | Autosome |
| CHAAL00000001927 | APOBEC2 | -1.2057057 | Autosome |
| CHAAL00000003990 | TLL2 | -1.2057057 | Autosome |
| CHAAL00000006540 | GRHL2 | -1.2057057 | Autosome |
| CHAAL00000007930 | MRPS2 | -1.2057057 | Autosome |
| CHAAL00000007981 | RDH10 | -1.2057057 | Autosome |
| CHAAL00000009595 | EDEM3 | -1.2057057 | Autosome |
| CHAAL00000010919 | PUS1 | -1.2057057 | Autosome |
| CHAAL00000008432 | | -1.1972611 | Autosome |
| CHAAL00000008685 | | -1.1972611 | Autosome |
| CHAAL00000014103 | KIF1BP | -1.1972611 | Autosome |
| CHAAL00000000216 | SLC51B | -1.1913506 | Autosome |
| CHAAL00000001031 | | -1.1913506 | Autosome |
| CHAAL00000001836 | | -1.1913506 | Autosome |
| CHAAL00000002954 | SLC25A15 | -1.1913506 | Autosome |
| CHAAL00000003274 | GUCY1A1 | -1.1913506 | Autosome |
| CHAAL00000004279 | KCTD14 | -1.1913506 | Autosome |
| CHAAL00000006283 | RANBP17 | -1.1913506 | Autosome |
| CHAAL00000008866 | FNDC9 | -1.1913506 | Autosome |
| CHAAL00000011277 | LOC105413758 | -1.1913506 | Autosome |
| CHAAL00000011314 | SERAC1 | -1.1913506 | Autosome |
| CHAAL00000011488 | EXOC3 | -1.1913506 | Autosome |
| CHAAL00000014023 | ASCC2 | -1.1913506 | Autosome |
| CHAAL00000014806 | | -1.1913506 | Autosome |
| CHAAL00000015167 | ARV1 | -1.1913506 | Autosome |

19. Conesa A, Götz S, García-Gómez JM, Terol J, Talón M, Robles M. Blast2GO: a

universal tool for annotation, visualization and analysis in functional genomics research. Bioinformatics. 2005;21:3674–6.
20. Götz S, García-Gómez JM, Terol J, Williams TD, Nagaraj SH, Nueda MJ, et al. High-throughput functional annotation and data mining with the Blast2GO suite. Nucleic Acids Res. 2008;36:3420–35.
21. Zhang G, Li B, Li C, Gilbert MTP, Jarvis ED, Wang J, et al. Comparative genomic data of the Avian Phylogenomics Project. Gigascience. 2014;3:26.
22. Altschul SF, Gish W, Miller W, Myers EW, Lipman DJ. Basic Local Alignment Search Tool. J Mol Biol. 1990;215:403–10.
23. Edgar RC. MUSCLE: multiple sequence alignment with high accuracy and high throughput. Nucleic Acids Res. 2004;32:1792–7.
24. Capella-Gutiérrez S, Silla-Martínez JM, Gabaldón T. trimAl: a tool for automated alignment trimming in large-scale phylogenetic analyses. Bioinformatics. 2009;25:1972–3.
25. Stamatakis A. RAxML-VI-HPC: maximum likelihood-based phylogenetic analyses with thousands of taxa and mixed models. Bioinformatics. 2006;22:2688–90.
26. Stamatakis A, Hoover P, Rougemont J. A rapid bootstrap algorithm for the RAxML web servers. Syst Biol. 2008;57:758–71.
27. Paradis E, Claude J, Strimmer K. APE: Analyses of Phylogenetics and Evolution in R language. Bioinformatics. 2004;20:289–90.
28. Yang Z. PAML 4: Phylogenetic analysis by maximum likelihood. Mol Biol Evol. 2007;24:1586–91.
29. DePristo MA, Banks E, Poplin R, Garimella K V, Maguire JR, Hartl C, et al. A framework for variation discovery and genotyping using next-generation DNA sequencing data. Nat Genet. 2011;43:491–8.
